# Robust nucleation control via crisscross polymerization of DNA slats

**DOI:** 10.1101/2019.12.11.873349

**Authors:** Dionis Minev, Christopher M. Wintersinger, Anastasia Ershova, William M. Shih

**Author notes:** denotes equal contribution (Dionis Minev, Christopher M. Wintersinger).

## Abstract

Natural biomolecular assemblies such as actin filaments or microtubules polymerize in a nucleation-limited fashion^1,2^. The barrier to nucleation arises in part from chelate cooperativity, where stable capture of incoming monomers requires straddling multiple subunits on a filament end^3^. For programmable self-assembly from building blocks such as synthetic DNA^4–23^, it is likewise desirable to be able to suppress spontaneous nucleation^24–31^. However, existing approaches that exploit just a low level of cooperativity can limit spontaneous nucleation only for slow growth, near-equilibrium conditions^32^. Here we introduce ultracooperative assembly of ribbons densely woven from single-stranded DNA slats. An inbound “crisscross” slat snakes over and under six or more previously captured slats on a growing ribbon end, forming weak but specific half-duplex interactions with each. We demonstrate growth of crisscross ribbons with distinct widths and twists to lengths representing many thousands of slat additions. Strictly seed-initiated extension is attainable over a broad range of temperatures, divalent-cation concentrations, and free-slat concentrations, without unseeded ribbons arising even after a hundred hours to the limit of agarose-gel detection. We envision that crisscross assembly will be broadly enabling for all-or-nothing formation of microstructures with nanoscale features, algorithmic self-assembly, and signal amplification in diagnostic applications requiring extreme sensitivity.

## Introduction

DNA tiles or bricks have been shown to self-assemble non-periodic nanostructures much larger than what has been demonstrated thus far with any single DNA-origami scaffold, while maintaining an excess of components during the assembly reaction^15–17,21^. This is enabled by folding in a cooperative regime, where nucleation is relatively slow and growth is comparatively fast. Rate-limiting spontaneous nucleation can be relied upon to initiate growth, however control over the copy number of structures then will be limited. For periodic assembly, rate-limiting nucleation will lead to a wide distribution of sizes, which is disadvantageous when a relatively monodisperse product is desired. Introduction of seeds could provide a burst of controlled nucleation, nevertheless seed-independent nucleation will coincide under conditions that favor rapid growth (i.e. high concentrations of monomers and well below the reversible temperature), such that a subpopulation of smaller or incomplete assemblies will arise. In the case of algorithmic assembly, where each seed kinetically triggers a particular pattern of tile accretion, contamination from unseeded growths can be especially problematic^24–32^. Tile assembly faces a particularly stringent test for application in single-molecule amplification schemes for point-of-care diagnostics, where the speed of growth is at a premium while, conversely, the limit of detection is bounded by the rate of spontaneous nucleation.

We present a crisscross architecture implemented with single strands of DNA (ssDNA) acting as elongated slats that polymerize with alternating orthogonal orientations to weave ribbons of programmable widths and twists. The ability of crisscross slats to pass over or under each other allows for a large number of binding partners to be engaged simultaneously — beyond just nearest neighbors — and therefore creates the potential for an extraordinary degree of cooperativity to be realized. We demonstrate designs where stable recruitment of an incoming slat requires that it cross-bind either six or eight previously captured slats on a ribbon-end. Such a high level of cooperativity enables simultaneous realization of two often prized characteristics of programmable self-assembly that otherwise would be mutually exclusive: rapid, irreversible growth from introduced seeds along with near-complete suppression of spontaneous nucleation.

### How crisscross polymerization enables robust control over nucleation

In crisscross polymerization, a slat is a linear array of 2*n* weak-binding sites, each of equal strength and specific to a single conjugate site on one of 2*n* distinct cross-binding slats; *n* = 6 in the example in Figure 1a. Specificities are arranged such that alternating x-slats (blue) and y-slats (gold) can add sequentially to the ribbon end by securing *n* consecutive cross-binding interactions each (Figure 1a, bottom, Supplementary Information section S1.1). Near the reversible temperature for polymerization, all *n* of these cross-binding interactions are required for stable recruitment, and the critical nucleus for initiating growth consists of *n* pairs of x-slats and y-slats. Here the energetic credits from the *n*^2^ cross-binding interactions will offset only half the energetic debits (e.g. entropic fixation, electrostatic repulsion) from capture of 2*n* slats out of bulk solution (Figure 1b and Supplementary Information section S1.1). Therefore the mean time for spontaneous nucleation near the reversible temperature will scale exponentially with *n*; for example, if the bulk slat concentration is 1 µM, and the effective concentration of captured slats is 20 M (i.e. entropic initiation penalty of −6 cal mol^-1^ K^-1^ estimated for duplex formation^33^) and ignoring enthalpic initiation penalties, then each designed increment of *n* yields up to a 20 million-fold slower rate of spontaneous nucleation (i.e. 20 M / 10^-6^ M).

**Figure 1.**
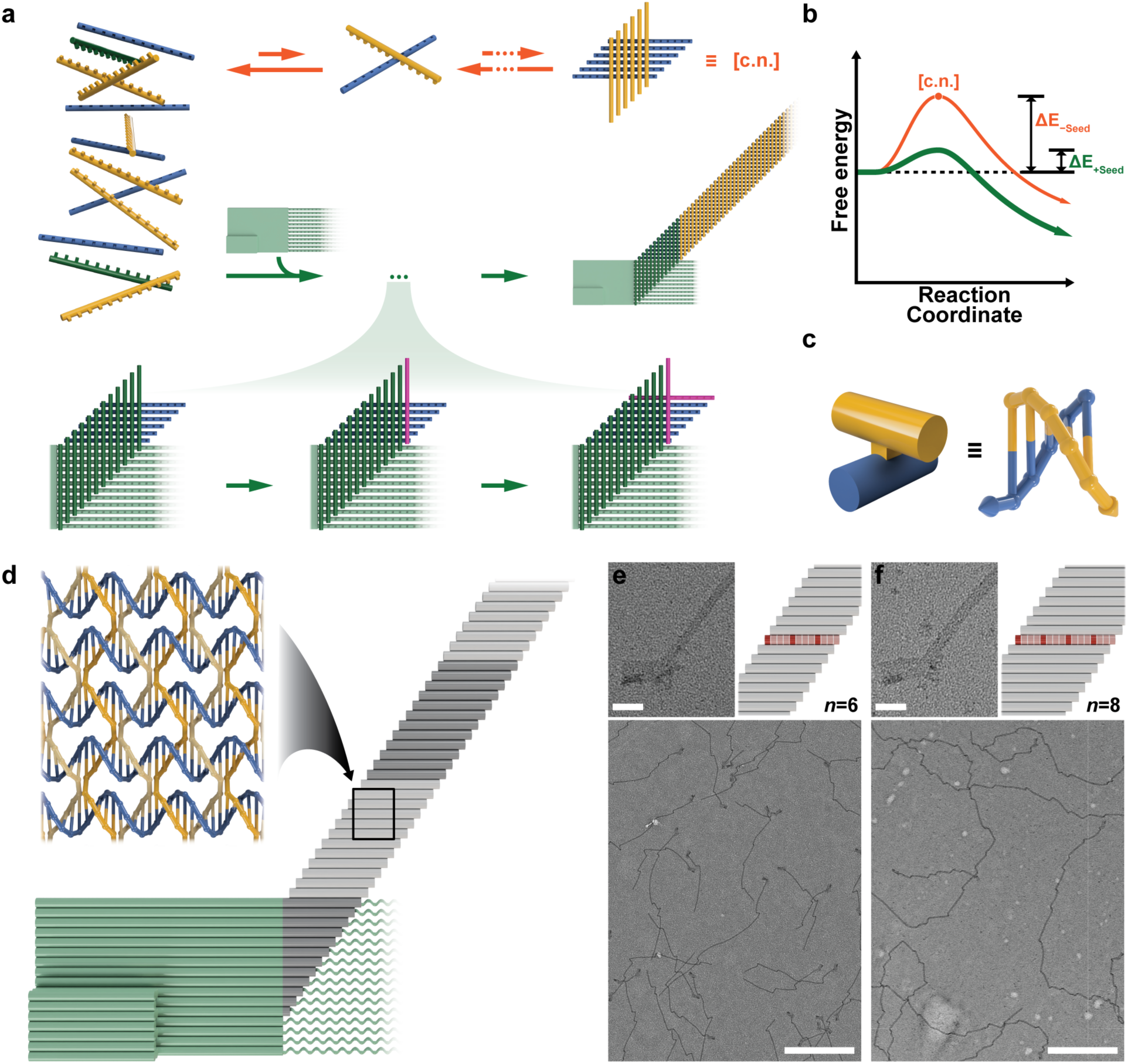
Crisscross polymerization of slats into ribbons. **a**, Abstract cartoon representation of assembly from x-slats (blue) and y-slats (gold) each with 12 binding domains (i.e. *n* = 6), at a temperature where six bonds are needed for stable attachment. The upper pathway (orange arrows) shows transient, unstable association between free slats. Formation of a critical nucleus (c.n.) requires an extended series of slat additions where, at each step, the growing intermediate is subject to highly favorable disintegration. Therefore, spontaneous nucleation is very rare. In contrast, the lower pathway (green arrows) exploits a seed (light green) for stable capture of twelve nuc-y-slats (dark green) along with six x-slats. Subsequent slats of orthogonal orientation (magenta) are recruited in alternating fashion to perpetuate ribbon growth. **b**, Energy landscape for seed-independent (orange) versus seed-dependent (green) nucleation and growth. **c**, Implementation of crisscross slats with a half-turn of ssDNA per binding domain. **d**, Rendering of crisscross polymerization of ssDNA slats (blue and gold) as triggered by a DNA-origami seed (light green). Straight cylinders represent double helices. **e**, Negative-stain TEM images of v6 (*n* = 6) ribbons. **f**, Negative-stain TEM images of v8 (*n* = 8) ribbons. In *e* and *f*, the renderings highlight the width difference of v6 versus v8 designs. Each boxed cell is a half-turn domain, with dark-red and light-red cells representing six and five base pairs, respectively. Scale bars are 50 nm for the top versus 800 nm for the bottom.

Most importantly, if the length of the crisscross slats is incremented by two segments (i.e. *n*’ = *n* + 1) but the temperature, salt, and slat concentrations are maintained such that still only *n* cross-bonds are needed for stable recruitment, then irreversible growth can be achieved while maintaining the identical-height barrier to nucleation. More generally, any target barrier height and degree of growth irreversibility can be engineered for a given high concentration of monomers simply by designing slats with a sufficiently large *n* and adjusting temperature accordingly (Supplementary Information sections S1.1–3). Conversely, for square tiles that attach to an existing assembly with a maximum of two bonds (i.e. analogous to *n* = 2), the limited cooperativity allows for very little suppression of spontaneous nucleation under rapid growth conditions. For example, at 1 µM each square tile and 100× growth:shrinkage, and assuming 20 M effective concentration of captured tiles and no enthalpic initiation penalty, the critical nucleus is only a nominal 2.3×2.3 tiles in size. Furthermore, the free energy for assembling equivalent-size critical nuclei can be up to twice the magnitude for crisscross compared to square tiles. Indeed, Woods et al found they were unable to achieve seeded growth with satisfactory suppression of spontaneous nucleation for square tiles at 1 µM each (see Supplementary Information A, Section S5.1.1 and Figure S27A from Woods et al.^31^).

### Crisscross implementation with ssDNA slats

We sought the most compact slat architecture that we could conceive for satisfying the requirements of crisscross polymerization as described above, as this would enable economical use of the highest concentrations of monomers for the fastest growth. Here we report that a single strand of DNA, conceptualized as a linear array of domains half a turn long, can serve well as a crisscross slat (Figure 1c, d). For this study, we examined crisscross growth from “v6” slats (*n* = 6, Figure 1e) versus “v8” slats (*n* = 8, Figure 1f) (Supplementary Information section S2.1). Multiple ssDNA slats can assemble into crisscross ribbons comprised of staggered parallel double helices connected by antiparallel crossovers that occur every half turn (Figure 1d). Single-stranded slats alternate threading over and under consecutive cross-binding partners, instead of passing entirely above or below as depicted in our abstract-cartoon crisscross model (cf. Figure 1a versus Figure 1d). Base stacking propagates along the x-axis (i.e. parallel to the helices of the seed as in Figure 1d), such that x-slats never cross over and therefore follow a right-handed helical path, whereas y-slats cross over every half turn and therefore follow a left-handed helical path (Figure 1d).

We sought to investigate whether, with v6 versus v8 slats, robust periodic growth of crisscross ribbons from introduced seeds could be achieved without appreciable spontaneous nucleation over a wide range of slat concentrations, temperatures, and Mg^2+^ concentrations. To initiate programmed nucleation, we designed a DNA-origami seed that presents single-stranded scaffold loops pre-organized to cooperate in recruiting *n* unique nucleating y-slats (i.e. nuc-y-slats, dark green in Figure 1a; Supplementary Information section S2.2), followed by an additional *n* unique nuc-y-slats that alternate addition with *n* x-slats. This complex then can nucleate periodic growth from repeating sets of 2*n* alternating pairs of x- and y-slats. We additionally designed multiple DNA slat sequence variants for v6 and v8 (Supplementary Information section S2.3). For v8, we also exploited symmetry for repeating sets of only *n* pairs of x- and y-slats, or even only *n*/2 pairs, useful for reducing material costs while maintaining the same speed of growth (Supplementary Information section S2.4).

### Observation of seeded versus unseeded crisscross growth

We first verified that seeds could nucleate crisscross assembly with either v6 or v8 slats by observing ribbons with negative-stain transmission electron microscopy (TEM, Figure 1e, f, Supplementary Information section S3.1). We attribute the kinked appearance of the ribbons to trapping of twisted configurations on the TEM grids. We next compared relative ribbon-length distributions via agarose gel electrophoresis to determine the impact of varying temperature and divalent cation (i.e. MgCl_2_) concentration. With slats at 0.2 µM each and the seed at 2 nM, we grew ribbons for 16 hours using various v6 (v6.1, v6.2, v6.3; Figure 2a, Supplementary Information section S3.2.1) or v8 sequence variants (v8.1, v8.2; Figure 2b, Supplementary Information section S3.2.2). We define “optimal” growth conditions as those that produce ribbons with the largest and most uniform length distributions, as evidenced by a tight, slowly migrating band on an agarose gel. Slats of v6 design in 12, 14, or 16 mM MgCl_2_ formed ribbons optimally at temperatures from 44°C–50°C, whereas slats of v8 design gave optimal growth at temperatures 2°C–6°C higher. We confirmed that copy number of filaments is controlled precisely by the number of starting seeds added to the reaction (Supplementary Information section S3.3). Adding blocking strands sequestering one or more of the nuc-y-slats led to termination of assembly under certain growth conditions (Supplementary Information section S3.4).

**Figure 2:**
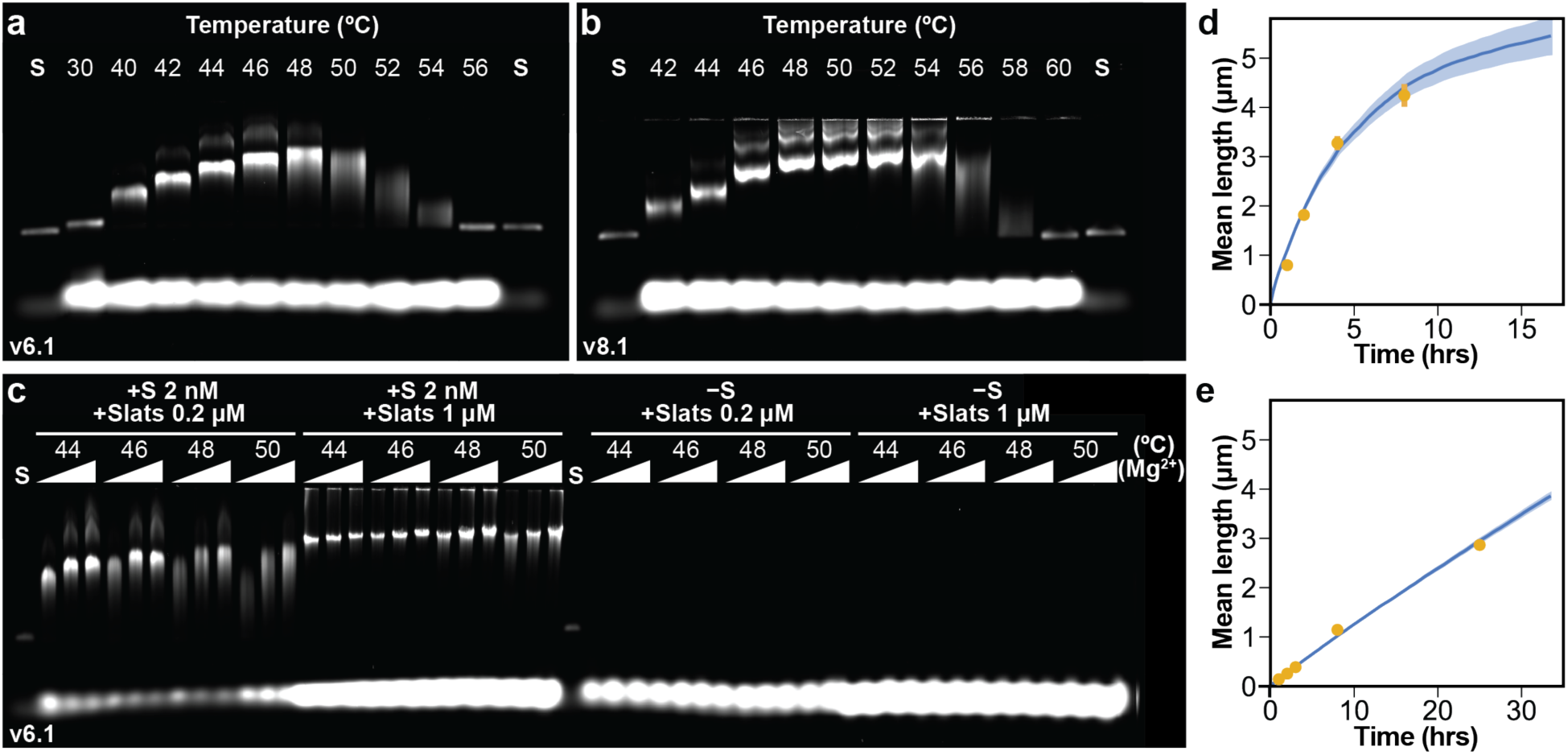
Characterization of DNA slat growth with and without the seed. Seeded ribbons from v6.1 (**a**) and v8.1 (**b**) slats, where different isothermal assembly temperatures are compared on agarose gels (0.2 µM each slat, 2 nM seed, 14 mM MgCl_2_, ∼16 hours growth). Lane (S) is the seed only. **c**, Agarose gels of DNA slats incubated isothermally ∼16 hours with (left) or without (right) the seed at optimal assembly temperatures. Concentration of slats was either 0.2 µM or 1 µM each slat with, as indicated by **⊿**, 12, 14, or 16 mM MgCl_2_. No assembly was observed in the absence of the seed. **d–e**, Direct length measurements of v6.1 ribbons at timepoints (gold) as observed in TEM, with length predictions derived from the stochastic model (blue). *d* shows faster assembly with optimal growth conditions (50°C, 14 mM MgCl_2_, 1 µM each slat, 2 nM seed) whereas *e* shows slower assembly with suboptimal lower temperature conditions (40°C, 20 mM MgCl_2_, 1 µM each slat, 2 nM seed). For panel *d* and *e*, each gold data point is the mean ±SEM length (N=130–168 ribbons measured per point; note that error bars are obscured for most of the data points), and N=150 ribbons for the model (shaded range is ±SEM).

To assay for spontaneous nucleation, we next increased the concentration of slats to 1 µM each and incubated them for 16 hours at the temperature and MgCl_2_ ranges optimal for growth described above. We estimated our limit of detection at ∼65 pg per agarose-gel lane, corresponding to a half-million 5 µm ribbons (assuming ∼3300 slats contributing 1.5 nm extension each), or ∼200 fM for a 4 µL reaction volume (Supplementary Information section S4.1). Notably, seeded reactions with 1 µM versus 0.2 µM each slat always yielded more uniform length distributions. Spurious (i.e. unseeded) ribbons were undetectable on agarose gels for v6.1 and v6.2 slats (Figure 2c, Supplementary Information section S4.2); however, they were observed for select conditions for v6.3 slats (Supplementary Information section S4.3). Furthermore, the v8 sequence variants demonstrated no observable spontaneous nucleation (Supplementary Information section S4.4). We next incubated v6.1 and v6.2 slats for an extended 100 hour period, at temperatures from 46°C– 52°C with 16 mM MgCl_2_ and 1 µM each slat, and observed no spurious ribbons, except for a faint band near the gel detection limit for v6.1 slats at 46°C (Supplementary Information section S4.5). Next, we experimentally ascertained that the reversible temperature for binding of the v6.1 and v6.2 slats is ∼56°C at 16 mM MgCl_2_ and 1 µM each slat (Supplementary Information section S4.6). Hence, we concluded that slats at 1 µM concentration can be seeded into ribbons with spurious assembly below our gel-detection limit (<200 fM after 100 hours) under rapid, highly irreversible growth conditions (i.e. 10°C below the reversible temperature).

We also incubated 1 µM each v6.1 slat at suboptimal growth temperatures (i.e. 34°C–44°C) for 16 hours at 16 mM MgCl_2_ and observed increasing occurrence of spurious ribbons, albeit with progressively slower growth as the temperature was lowered (Supplementary Information section S5.1). Conversely, for v6.1 slats we found that increasing temperature (i.e. from 41.0 to 44.7°C), decreasing the concentration of slats (i.e. from 1.0 to 0.2 µM), or decreasing MgCl_2_ (i.e. from 16 to 10 mM) could each individually lessen relative spurious ribbons between 100–1000 fold (Supplementary Information section S5.2). Furthermore, we found that v6.2 was particularly resilient to spontaneous nucleation with 10–100 fold fewer spontaneously formed ribbons compared to v6.1 at a given temperature (Supplementary Information section S5.3). However, this decreased rate of spontaneous nucleation came at the cost of decreased uniformity of growth, as shown by gel bands that are less tight.

Next, we characterized the kinetics of slat assembly by directly measuring ribbon lengths in TEM micrographs. We characterized growth with the optimized conditions noted above (i.e. 50°C, 16 mM MgCl_2_, 1 µM each slat; Figure 2d, Supplementary Information section S6.1.1). Growth of v6.1 slats for one hour yielded ribbons with a mean length of ∼800 nm. After eight hours, the mean length exceeded 4 µm. We further characterized growth of v6.1 ribbons at an assembly temperature and MgCl_2_ concentration outside of the optimal range (40°C, 20 mM MgCl_2_, 1 µM slats; as shown in Figure 2e, Supplementary Information section S6.1.2). The rate of ribbon growth was much slower, with mean length of ∼140 nm after one hour and a mean length of only ∼3 µm after 24 hours. Furthermore, we observed spurious nucleation of ribbons at this suboptimal lower temperature and high MgCl_2_ condition (Supplementary Information section S6.2). In an attempt to account for how an interplay of nucleation, extension, and termination might describe our observations, we developed a stochastic model and derived an analytical solution for DNA slat assembly. Our model is in general agreement with length measurements obtained from TEM images and gel data for both optimal and suboptimal growth conditions (Figure 2d–e, Supplementary Information section S6.1–5).

To illustrate how crisscross polymerization could be used for detection of a real-world analyte, we designed a set of six nuc-y-slats that folds a 192 nt segment of a ssDNA viral genome (M13 in this case); this structure then can bind six x-slats and thereby form a nucleus for periodic v6 ribbon assembly (Supplementary Information section S7). For smaller analytes with insufficient binding energy to recruit six nuc-y-slats to high local concentration, linking crisscross growth to target detection will likely require a different approach.

A diversity of ribbon shapes could prove especially helpful for multiplexed detection. Therefore we further generalized crisscross morphology by designing slats that assemble into tubes and coiled ribbons (Figure 3a). To achieve this, we tested v8 slat variants where either the pattern of 5 and 6 nt domains is shifted, or the number of base pairs per turn for each four-unit set of domains is increased (i.e. from 10.5 bp/turn to 11bp/turn; Supplementary Information section S8.1). Shifting the arrangement of the domains in v8.1 slats, while maintaining a 10.5 bp/turn reciprocal twist, led to a loose coiling of the ribbons (Figure 3d, Supplementary Information section S8.2). Underwinding the DNA to 11.0 bp/turn in v8.3 slats led to more tightly coiled ribbons (Figure 3b, Supplementary Information section S8.2). We added short sticky ends to the slats to increase the propensity of closure of the coiled sheets into tubes of varied diameters (Figure 3c and e, Supplementary Information section S8.3). Interestingly, v8.3 slats exhibited an especially rapid rate of addition (estimated second-order rate constant 10^6^ M^-1^s^-1^ for v8.3 versus 0.5×10^6^ M^-1^s^-1^ for v6.1; Supplementary Information sections S6.1, S8.4) and resulted in a constant narrow-tube diameter, while maintaining no observable spurious nucleation under optimal growth conditions (Supplementary Information section S8.5).

**Figure 3:**
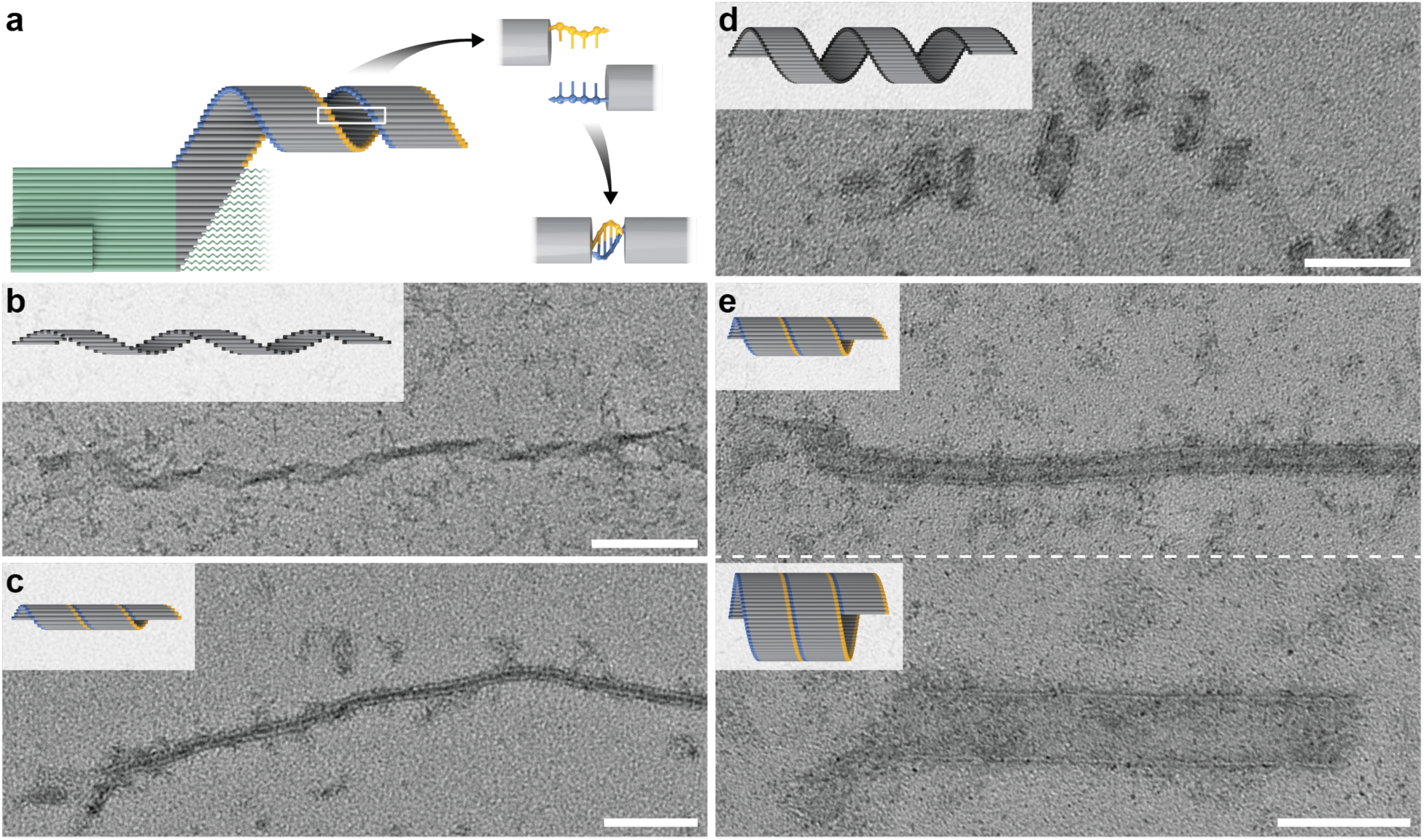
Formation of tubes and coiled ribbons. **a**, Seeded growth of a coiled ribbon. Coiled ribbons can be closed with sticky-end overhangs, as indicated with gold and blue. **b**, v8.3 slats without sticky ends form open, tightly coiled ribbons. **c**, v8.3 slats with 3 nt sticky ends form closed tubes of constant diameter. **d**, v8.1 slats without sticky ends form open and loosely coiled ribbons (note the difference from the coiled ribbon in *b*). **e**, v8.1 slats, with 4 nt sticky ends, form closed tubes of varied diameters in a single reaction. Scale bars are 100 nm.

## Summary and Conclusions

Crisscross polymerization combines the unbounded size of DNA brick or tile nanoconstructions with the fast folding and copy-number control of DNA origami. The remarkable robustness of DNA-origami folding obviates the need for negative sequence design, strand purity, and precise tuning of strand stoichiometries, and makes DNA origami extremely attractive for general use. Likewise, crisscross assembly with sufficiently long slats offers robust nucleation control over a wide range of temperatures and slat concentrations. Similar to past developments with other motifs in structural DNA nanotechnology^9,12,15,16^, crisscross assembly with ssDNA slats should prove adaptable to 2D and 3D growth architectures. We also envision the exploration of dendrimeric building blocks with reach-over appendages that facilitate ultracooperativity with alternate assembly geometries, as observed with naturally occurring clathrin triskelions^34,35^. Moreover, crisscross polymerization will likely prove generalizable to slats constructed from other macromolecules or supramolecular assemblies, across a wide range of length scales, in cases where weak, specific binding sites can be programmed to occur in a linear arrangement.

## Materials and Methods

### DNA slats sequence design

DNA slat sequences were designed using custom Python scripts (available upon request); base-pairing energies were assessed using Unafold^36^. All sequences were designed to have minimal self-structure. The v6.1 slats were designed to have stronger base-stacking in the y-direction than in the x-direction, as assessed using Unafold. The v6.3 and v8.1 slats were designed to have stronger base-stacking in the y-direction than in the x-direction, based on stacking energies reported in Protozanova et al^37^.

### DNA slats denaturing polyacrylamide gel electrophoresis (PAGE) purification

Dehydrated DNA slat oligonucleotides were purchased from Integrated DNA Technologies (IDT) at 10 or 100 nmole scale. SequaGel UreaGel System (National Diagnostics, EC-833) reagents were used to prepare 15% denaturing PAGE gels in empty plastic 1.5 mm mini-gel cassettes (Invitrogen Novex™, NC2015). Each DNA slat was rehydrated at 100 or 700 µM (assuming 70 nmole/well for 100 nmole dry IDT oligo order) in Milli-Q water, slats were combined into pools (for nuc-y-slats, x-slats, and y-slats respectively), and pools were mixed 1:1 by volume with 95% formamide, 0.025% (w/v) bromophenol blue, 0.025% (w/v) xylene cyanol, and 5mM EDTA. Samples were loaded into the gels and run at 200–300 V for ∼40–50 minutes, until the bromophenol blue dye front had run off the gel. Bands were identified on the gels by shadowing with UV light, so that bands for each pool could be excised with a razor blade. Gel slices were crushed in a 2 mL round-bottom tube with a pestle, soaked at room temperature on a shaking incubator at 1500 rpm in an excess of 1X TE buffer (5 mM Tris and 1 mM EDTA) overnight, at which point waste acrylamide was separated from the aqueous slats solution with Freeze N’ Squeeze spin columns (Bio-Rad, 732-6166). The DNA slats were further purified by precipitation in isopropanol, two washes in cold ethanol, and final resuspension in water, with the volume sufficient to attain an approximate concentration of 10 µM per each slat in the pool. A Nanodrop 2000c Spectrophotometer (Thermo Scientific™) was used to determine the final concentration. All DNA slat sequences are reported in Supplementary Data S1.

### Agarose gel electrophoresis

Gel characterization of the origami seed or assembled slat filaments was performed using either the Thermo Scientific™ Owl™ EasyCast™ B2 or D3-14 electrophoresis system. UltraPure agarose (Life technologies, 16500500) was melted in 0.5X TBE (45 mM Tris, 45 mM boric acid, 0.78 mM EDTA, 12–16 mM MgCl_2_ to match amount in slats experiment) to a concentration of 0.5–2.0 % (w/v). The 0.5–1% gels were typically used to characterize larger assemblies of slats, whereas 2% gels for smaller structures such as the origami seed. The molten agarose was cooled to 65°C and gel stain was added. All slat assembly reactions where characterized with 10000x SYBR Gold (Thermo Fisher, S-11494) gel stain added to a final concentration of ∼0.183x (i.e. 3 µL per 160 mL molten agarose). Origami seed folding was characterized with 6.25×10^-5^ % (w/v) ethidium bromide (i.e. 10 µL per 160 mL molten agarose; Bio Rad, 1610433). Gels were covered with aluminum foil during solidification and running to lessen exposure to ambient light. DNA assemblies were mixed in an excess of agarose gel loading buffer (5 mM Tris, 1 mM EDTA, 30% w/v glycerol, 0.025% w/v xylene cyanol, 12–16 mM MgCl_2_; with typically 10 µL loading buffer added to 4–10 µL of each assembly). The mixed samples were loaded onto the gel and separated for 3–4 hours at 60 V at room temperature. Control samples for size and densitometry included one or both of the following: first, 0.5–11.2 fmoles of folded DNA-origami seed; or second, 0.00625–0.5 µg of Gene Ruler 1kb Plus DNA Ladder (Thermo Scientific™ SM1331). Gel images were captured on a GE Typhoon FLA 9500 fluorescent imager using the SYBR-Gold parameters as given in the Typhoon control software. The photo-multiplier tube (PMT) was set to 300–500V, with the value varied depending on the amount of sample loaded. Densitometry to quantify relative assembly of DNA bands was performed with ImageJ (v2.0.0-rc-69/1.52i)^38^. Background subtraction with a rolling ball radius of 30–60 pixels was performed on linear TIFF images. The GelAnalyzer plugin in ImageJ and wand tool was used to integrate total pixel intensities from lanes of interest. Gel absorbency data collected on different agarose gels were compared to one other by normalizing them with respect to the DNA-origami seed control, as well as the volume of reaction loaded.

### DNA-origami seed

Dehydrated staple oligonucleotides were purchased from Integrated DNA Technologies (IDT) at 10 nmole scale. The p8064 scaffold strand was produced from M13 phage replication in *Escherichia coli*. The DNA-origami seed was folded with 10 nM p8064 scaffold and 100 nM of each staple strand in 1X TE buffer (5 mM Tris and 1 mM EDTA) containing 6 mM MgCl_2_. The reaction was incubated on a PTC-225 Peltier Thermal Cycler (MJ Research) with the following temperature program: 90°C for 2 min, cool to 55°C and decrease to 50°C over 18 hours by decreasing the temperature at a rate of −1°C/3 hours, and then holding the temperature at 4°C thereafter. The folded reaction was analyzed on agarose gel and by observation with TEM. The folded seed was stored at 4°C in the raw folding mixture. The seed was noted to remain stable in 4°C storage for up to a year and was used directly from the raw folding reaction in slat assembly experiments. Staple and scaffold DNA sequences are reported in Supplementary Data S1.

### DNA slat filament and tube assembly reactions for sequence variants with *n* = 6 domains, or *n* = 8 domains

10X reaction buffers were prepared for all reactions as follows: 35 mM Tris, 7 mM EDTA, 0.1% Tween-20 (Sigma Aldrich #P9416), 108–200 mM MgCl_2_ depending on the final MgCl_2_ in the reaction. The use of 10X reaction buffers was preferred to lessen variability in the final concentration of MgCl_2_ and Tween-20 between experiments. PAGE purified DNA slat pools and the DNA-origami seed were added to diluted reaction buffer to achieve the final concentrations: 0.2–1 µM DNA slats, 0.02–2 nM seed, and 1X reaction buffer. It is noteworthy that PAGE purification of the slats was critical to achieve filament growth. Attempts to assemble the raw slats as purchased from Integrated DNA Technologies (IDT) did not result in appreciable growth (data not shown). The reactions were assembled isothermally on a PTC-225 Peltier Thermal Cycler (MJ Research) or a Tetrad 2 Peltier thermal cycler (Bio-Rad) at a suitable growth temperature (e.g. 46–56°C) for various times (e.g. 1–100 hours, but overnight, 16 h growth, was most typical). We also assembled reactions using two-step temperature incubations (see Supplementary Information section S5) where a lower nucleation temperature (e.g. <46°C) was followed with a higher, more favorable growth temperature (e.g. 46–56°C). While spurious background assembly from room temperature conditions, when the nuc-y-, x-, and y-slats were mixed, was not typically identifiable, we either prepared reactions where the seed was not initially added so the slats could be denatured at 85°C for 10 minutes or added all components with the exception of the x-slats to the reaction and incubated at 60°C for 3–4 min. These reactions were then cooled to the isothermal growth temperature, at which point the reaction was briefly uncapped so that the seed or x-slats could be mixed into the reaction. This approach was used in slat characterization experiments where any spurious background assembly might have skewed the results. Alternatively, experiments where we tested for spurious assembly in the absence of a seed were prepared as above, but without adding the seed and denaturing the slats at 85°C for 10 minutes prior to incubations at lower temperatures.

### Note about poly-thymidine (poly-T) brushes and DNA slat filament assembly reactions for sequence variants 8.1 and 8.2

We found that the enzymatic addition of poly-T brushes on the 3’–ends of the v8 slats with terminal transferase (TdT) are important to ensure assembled filaments did not aggregate with one another. We used TdT to add a poly-T brush to the 3’–end of each slat, since slats were already 90 nt in length (further length increases result in significant upcharge when purchased from a commercial vendor). In all of the v6 slats, the poly-T brush was designed into the commercially purchased slat sequence. For the v8 slats used to form tubes, aggregation was less prevalent because their tube morphology concealed the aggregation-prone edges of the assembled sheet of slats to associate with themselves. We found variants 8.1 and 8.2 especially prone to aggregation when grown into long filaments, as with overnight growth at temperatures ≥ 46°C. TdT poly-T brush reaction: PAGE purified x- and y-slats were separately combined with dTTPs (Fisher Scientific, R0171) at a molar ratio of 1:125, terminal transferase (NEB, M0315L) in 1X terminal transferase reaction buffer (NEB, M0315L) and 0.25 mM CoCl_2_ (NEB, M0315L). The reaction was incubated for 30 minutes at 37°C, followed by heat inactivation of the enzyme at 70°C for 10 minutes. The slats were then separated from the reaction buffer by precipitating them in isopropanol, washed twice in cold ethanol, and resuspending them in water at the desired concentration. The DNA slat assembly reactions were then prepared as described previously.

### Transmission electron microscopy (TEM)

Filament samples were prepared by diluting slat assembly reactions 1:10–1:200 in 1X reaction buffer matching the buffer in the initial assembly reaction. The DNA-origami seed was diluted in 1X TE buffer with 6 mM MgCl_2_. FCF400-CU-50 TEM grids (Fisher Scientific, 5026034) were negatively glow discharged at 15 mA for 25 s in a PELCO easiGlow. The diluted sample (3 µL) was applied to the glow discharged grid, incubated for 2 minutes, and wicked off completely into Whatman paper (Fisher Scientific, 09-874-16B). Immediately after, 3 µL of 2% aqueous filtered uranyl formate, which in certain instances was pH adjusted with 25 mM sodium hydroxide, was applied and incubated for 1–2 seconds. The uranyl formate was wicked off completely into Whatman paper. All imaging was performed at 80 kV on a JEOL JEM 1400 plus microscope.

### Measurements and statistics

TEM filament length measurements were obtained manually using the ‘segmented line’ tool in ImageJ (v2.0.0-rc-69/1.52i)^38^. At each condition of interest, lengths of 130–168 filaments were recorded from TEM images showing distinguishable single filaments where the origami seed was identifiable.

## General

The authors would like to thank Richard Guerra, Jaeseung Hahn and Rasmus Sørensen for fruitful discussions, Jonathan F. Berengut for helpful insights into 3D modelling tools and providing his 3D ball-and-stick DNA model, and Erik Winfree for discussions on comparing crisscross versus square-tile nucleation.

## Author contributions

D.M. and W.M.S. designed the DNA slats architecture. D.M. and W.M.S. wrote the theoretical comparison. D.M., A.E., and W.M.S. designed the DNA slat sequences. D.M., C.M.W., and W.M.S. designed the experiments. D.M. and C.M.W. performed the experiments. D.M., C.M.W., A.E., and W.M.S. analyzed the data. D.M., C.M.W., A.E., and W.M.S. designed the stochastic model. A.E. wrote and performed the computations regarding the stochastic model. A.E. designed, wrote, and performed the computations regarding the analytical solution for the stochastic model. D.M., C.M.W., A.E. and W.M.S. wrote the paper.

