## Supplementary Information for "Robust nucleation control via crisscross polymerization of DNA slats"

**Title**

[#]denotes equal contribution

### Table of contents

**S1—Principles of crisscross nucleation** **4**
S1.1—Energetic landscape for spontaneous and seed-initiated nucleation of DNA slats 5
S1.2—Comparison of barrier height for crisscross versus square tile nucleation 6
S1.3—Derivation of equations 11

**S2—ssDNA slat architecture** **15**
S2.1—Critical nuclei comprised of ssDNA slats 15
S2.1.1—v6 critical nucleus 15
S2.1.2—v8 critical nucleus 16
S2.2—DNA-origami seed design and characterization 17
S2.3—Sequence design of DNA slats 18
S2.4—Sequence symmetry of DNA slats versus number of repeating slats 19

**S3—Determining optimal growth conditions for seeded ribbons** **21**
S3.1—Additional TEM images of v6 and v8 ribbons 21
S3.1.1—v6.1 ribbons 21
S3.1.2—v8.2 ribbons 22
S3.1.3—v8.7 high-sequence symmetry ribbons 22
S3.2—Growth of ribbons under optimal conditions 23
S3.2.1—v6 designs 23
S3.2.1.1—v6.1, v6.2, v6.3 0.2 µM each slat 23
S3.2.1.2—v6.1 1 µM each slat 24
S3.2.2—v8 designs 24
S3.2.2.1—v8.2 0.2 µM each slat 24
S3.2.2.2—v8.1 1 µM each slat 25
S3.3—Copy number control of ribbons with the seed 26
S3.4—Blocking nuc-y slats impedes ribbon assembly 27

**S4—Assay for spontaneous nucleation under optimal growth conditions** **29**
S4.1—SYBR-Gold gel detection limit of ribbons 29
S4.2—Spurious nucleation not observable for v6.2 with optimal growth conditions 31
S4.3—Spurious nucleation is observable for v6.3 under select optimal growth conditions 31
S4.3.1—Gel characterization of spurious nucleation of v6.3 31
S4.3.2—TEM validation that spuriously formed v6.3 gel bands are indeed ribbons 32
S4.4—Spurious nucleation is not observable for v8.1 and v8.2 under optimal growth conditions 33
S4.5—Extended 100 hour incubation of v6.1 and v6.2 under optimal growth conditions 33
S4.6—Determination of reversible temperature for binding v6.1 and v6.2 slats 34

**S5—Quantification of spontaneous nucleation under suboptimal low-temperature growth conditions** **36**

S5.1—Increased spurious nucleation observed at low temperatures 36
S5.2—Reducing spurious nucleation by increasing temperature, decreasing concentration of slats, or decreasing $MgCl_2$ concentration 40
S5.3—Reducing spurious nucleation by changing the sequence design from v6.1 to v6.2 42

**S6—Ribbon assembly characteristics by experimental observation and modelling 44**
S6.1—Kinetics by TEM observation versus model fit 46
S6.1.1—v6.1 at optimal growth conditions 47
S6.1.2—v6.1 at sub-optimal growth conditions 48
S6.1.3—v6.2 at optimal growth conditions 49
S6.1.4—v6.2 at alternative optimal growth conditions with 1.5 µM each slat 50
S6.2—Observation of spurious ribbons on agarose gels versus model fit 51
S6.3—Analytical solution of ratiometric gel analysis 54
S6.4—Analytical solution derivation 55
S6.5—Consideration of delayed seed initiation 56

**S7—Crisscross polymerization to sense nucleic acid sequences 59**

**S8—Twisted ribbons and tubes 60**
S8.1—Design notes on arrangement of binding sites to change ribbon morphology 61
S8.2—Results from shifting arrangement of binding sites 62
S8.3—Results from adding short sticky ends to close the coiled sheets into tubes 64
S8.4—Faster rate of assembly for 11.0 bp/turn design for underwound v8.3 65
S8.5—No observable spurious nucleation for v8.3 under optimal growth conditions (≥52ºC for 1 µM each slat) 67

**References 69**

#### S1—Principles of crisscross nucleation

**Overview**: In **S1.1**, we illustrate graphically the free-energy profile for spontaneous assembly of the critical nucleus for crisscross polymerization. In **S1.2**, we show that the barrier height ($G_{k*}$) is simply the size of the critical nucleus ($k^*$ pairs of x-slats and y-slats) multiplied by the free energy required to capture a slat from bulk solution into a ribbon ($G_{mc}$). The size of the critical nucleus $k^*$ is equivalent to the number of cross-binding interactions needed for stable attachment of a slat to the critical nucleus at the given temperature, up to the maximum of half the number of cross-binding sites in a slat (*n)*. Once the ribbon has passed the nucleation phase and is in the extension phase, then the excess cross-binding interactions ($n$ - $k^*$) promote the irreversibility of growth ($\boldsymbol{\varepsilon} = G_{mc}(n - k^*)/k^*$)). Thus any desired barrier height $G_{k*}$ and irreversibility $\boldsymbol{\varepsilon}$ can be achieved by scaling $n$ and then adjusting temperature to distribute $n$ between $k^*$ (i.e. size of critical nucleus and therefore height of nucleation barrier) and $n$ - $k^*$ (i.e. growth irreversibility) as desired.

| | | |
|---|---|---|
| $k^* = n/(1 + \boldsymbol{\varepsilon}/G_{mc})$ | or | $\boldsymbol{\varepsilon} = G_{mc}\,(n/k^* - 1)$ |
| $G_{k*} = k^* G_{mc}$ | or | $G_{k*} = [n/(1 + \boldsymbol{\varepsilon}/G_{mc})]G_{mc}$ |

- $k^*$ describes the size of the critical nucleus (i.e. $k^*$ pairs of x-slats and y-slats);
- $n$ describes the length of the slat (i.e. $2n$ binding sites; note that $k^* \le n$) ;
- $\boldsymbol{\varepsilon}$ is a dimensionless term representing the excess free energy from capture of a slat with $n$ binding interactions;
- $G_{mc}$ is a dimensionless term representing the free energy required to capture a free slat from bulk solution into a crisscross ribbon;
- $G_{k*}$ is a dimensionless term representing the free energy required to assemble the critical nucleus

Square tiles can provide substantial suppression of spontaneous nucleation at low tile concentration (i.e. high $G_{mc}$) and low irreversibility (i.e. low $\boldsymbol{\varepsilon}$). In **S1.2**, we remind the reader that this collapses at high tile concentration (i.e. low $G_{mc}$) and high irreversiblity (i.e. high $\boldsymbol{\varepsilon}$). We demonstrate that for an example case with 1 µM each square tile, 100× growth:shrinkage, and captured-tile effective concentration of 20 M, the nominal size $k^*$ of the critical nucleus for square tiles is 2.3 (i.e. 2.3 × 2.3). In contrast, crisscross under the equivalent conditions with $n$ = 6 yields $k^*$ = 4.7 and ~1e23 slower nucleation, and with $n$ = 8 yields $k^*$ = 6.3 and ~1e35 slower nucleation.

We also show that for a given sized critical nucleus $k^*$, the ratio of barrier heights comparing crisscross slats versus square tiles approaches a factor of two as $k^*$ increases: $G_{k*\_cc}/G_{k*\_st} = 2 - 1/k^*$

#### S1.1—Energetic landscape for spontaneous and seed-initiated nucleation of DNA slats

In **Supplementary Figure S1**, we illustrate the free-energy profile for the unfavorable spurious birth versus favorable seeded birth of a crisscross ribbon, using our abstract cartoon model for crisscross architecture. In the absence of a seed, DNA slats must form a rate-limiting critical nucleus, shown in step 11, before the capture of new slats to the end of ribbons is energetically favorable. The leading edge of the critical nucleus has sufficiently many overhanging binding sites to cooperatively engage new slats. All steps prior to 11 are energetically unfavorable because the entropic penalty of recruiting slats is larger with respect to energetic gains of binding single or small numbers of weak half-turn binding sites. The seed has multiple single stranded binding sites fixed proximally to cooperatively engage initial nuc-y-slats, as shown in step 1 (seeded pathway, **Supplementary Figure S1b**). The kinetic barrier to nucleation is much lower with the seed and repeating sets of x- and y-slats are now added to grow ribbons.

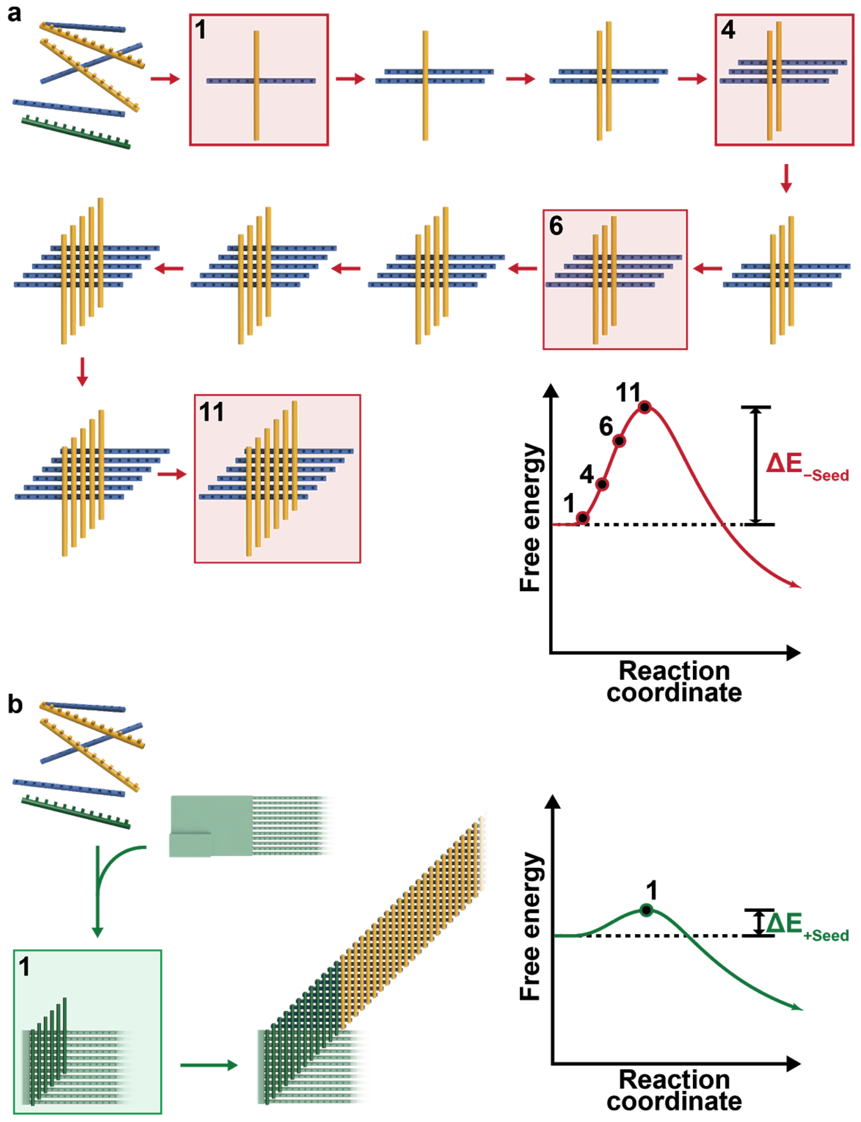


**Supplementary Figure S1**: Qualitative energetic landscape for spontaneous versus seed-initiated nucleation of crisscross polymerization. **a**, Spurious nucleation must overcome a large kinetic barrier to assembly. **b**, Seed-initiated nucleation provides an alternate lower barrier route to assembly.

In **Supplementary Figure S2**, we illustrate that the maximum barrier height for crisscross polymerization is proportional to *n*, therefore the barrier for v8 ($n$ = 8) is 33% higher than that of v6 ($n$ = 6). The size of the critical nucleus is made larger by using longer slats with more stringent assembly conditions. There are four more sequentially arranged binding sites per slat with v8 versus v6 slats. More binding energy is therefore available to capture new slats at the end of v8 slat ribbons. The consequence is that optimal growth is attainable with conditions less prone to spurious nucleation, such as using higher reaction temperatures. The v6 versus v8 slats have critical nuclei that are 6+6 and 8+8 arrays of x- and y-slats respectively, so long as assembly is carried out at the appropriately high reversible temperatures. In principle, DNA slats could be further elongated to increase the kinetic barrier to assembly using reaction conditions more stringent than characterized here (e.g. lower $MgCl_2$ concentration, higher temperature, presence of chemical denaturant such as formamide, etc).

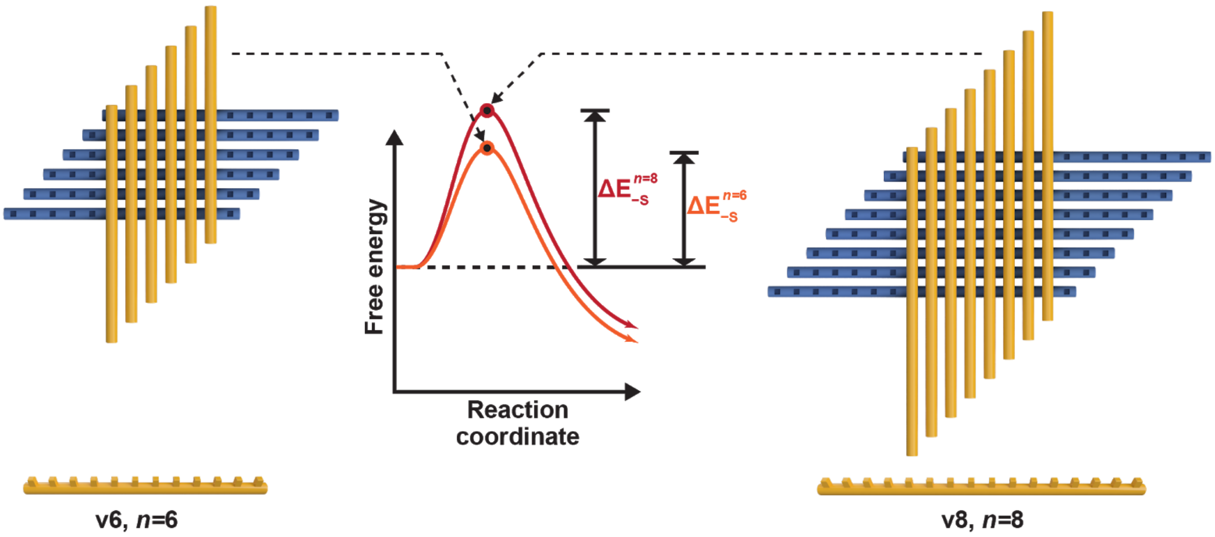


**Supplementary Figure S2**: Critical nuclei for v6 (n = 6) versus v8 (n = 8) slat assemblies. $\Delta E_{-S}$ denotes the energetic barrier without a seed being present in the assembly.

#### S1.2—Comparison of barrier height for crisscross versus square tile nucleation

In this section we adapt the previously developed kinetic tile assembly model (kTAM)[1,2] to crisscross assembly. We highlight the comparison of crisscross versus square-tile assembly primarily for far-from-equilibrium, fast growth conditions. We focus solely on the homogenous tile system with a single tile type that can form a total of four bonds ($n = 2$). For this we adapted the previously reported equations for square tiles (see section 4 Nucleation, equation 12 and figure 8, from Evans & Winfree[2]) to crisscross assembly (see **S1.3** below and **Supplementary Figure S6**). It is noteworthy, that other variants of square-tile assembly, such as zig-zag tile assembly[3–5] or square tiles that assemble into tubes[6,7] are potentially preferred in practice, since they allow for more uniform growth (i.e. each incoming tile adds with two bonds after the critical nucleus has formed) at near reversible conditions (small values of $\varepsilon$). However, here we focus on the key advantage of crisscross over square-tile assembly, namely the maintenance of strictly seed-dependent growth and suppression of spurious nucleation, in far-from-equilibrium, fast growth conditions. Under these assembly conditions (large values of $\varepsilon$ — e.g. $\varepsilon = 2$) the zig-zag tile assembly system does not display any advantages over homogenous square-tile assembly. Zig-zag tile assembly is only advantageous to use in near-equilibrium assembly conditions.

**The following assumptions were made for the square tile versus crisscross assembly comparison.**

We set a single energy unit to $2.3\ k_B T$, where $k_B$ is the Boltzmann constant and $T$ the temperature. We chose this to yield a $10\times$ per-unit change for variations of the ratio of $growth : shrinkage$ (equation 2) and variations in the DNA slat and square-tile concentration. Consequently equations 1 and 2 (adapted from equation 3 with $\varepsilon = nG_{se} - G_{mc}$, from Evans & Winfree[2]) define the relationship between the dimensionless parameter $\varepsilon$ and the $growth : shrinkage$ of the assembly. Equations 3 and 4 are a further adaptation from Evans & Winfree[2] (see section 2 Basic models, sentence above equation 2) defining the free monomer concentration $G_{mc}$.

$$r_f/r_{r,n} = growth : shrinkage = e^{2.3(nG_{se}-G_{mc})} \approx 10^{(nG_{se}-G_{mc})} = 10^{\varepsilon} \quad (1)$$

$$\varepsilon = log_{10}\left(\frac{r_f}{r_{r,n}}\right) \quad (2)$$

$$[C] = u_0 e^{-2.3(G_{mc}+\alpha)} \approx u_0 10^{-(G_{mc}+\alpha)} \quad (3)$$

$$G_{mc} = \alpha - log_{10}\left(\frac{[c]}{u_0}\right) \quad (4)$$

Where $r_f/r_{r,n}$ is the ratio for $growth : shrinkage$, $n$ is the number of correct bonds formed, $G_{se}$ is the free energy for a single bond, $u_0$ is a standard concentration of 1 M, $[c]$ the concentration per DNA slat or square tile monomer, and $\alpha$ a constant unitless parameter accounting for entropic fixation and electrostatic repulsion. For this section we assume $\alpha = log_{10}(20\ M/1\ M) = 1.3$ corresponding to entropy of initiation of -6 cal mol$^{-1}$ K$^{-1}$ [8]. For the purpose of this example, we set the free monomer concentration $G_{mc} = 1.3 - log_{10}\left(\frac{10^{-6}M}{1M}\right) = 7.3$, i.e. 1 µM each DNA slat or square tile monomer.

For an example of a fast, far-from-equilibrium growth condition, we pick the ratio of $growth : shrinkage = r_f/r_{r,n} = 100$ (i.e. $\varepsilon = log_{10}(100) = 2$). Both $n = 6$ and $n = 8$ crisscross assembly display a much larger nucleation barrier (~1e23 and ~1e35 fold slower rate of nucleation, respectively) and critical-nucleus size ($k^* = 4.7$ and 6.3, respectively) when compared to square-tile assembly ($k^* = 2.3$) under these conditions (**Supplementary Figure S3**). We explore next how crisscross maintains a larger critical nucleus in comparison to square-tile assembly for increasingly large values of $\varepsilon$ (**Supplementary Figure S4**). Only for $\varepsilon$ values smaller than 0.737 (for crisscross $n = 6$) and 0.524 (for crisscross $n = 8$) is the critical nucleus for square tiles assembly larger than crisscross, due to the asymptotic nature of equation 10 for small $\varepsilon$ values (see **S1.3** below). Note that the irreversibility for square-tile filament growth is the $\varepsilon$ of two-bond addition minus that for reversible growth of a filament of that width, the latter of which is nonzero due to corner tiles being captured with just a single bond. Zig-zag and tubular tile assembly systems eliminate these corner tiles through specific edge-tiles or circular arrangement (see section 4 Nucleation, figure 9, from Evans & Winfree[2]).

Even when crisscross and square tile assembly have identically sized critical nuclei, crisscross maintains an up to $2\times$ larger free energy barrier to assembly; the ratio $G_{k^*\_cc}/G_{k^*\_st}$ ($cc$ = crisscross, $st$ = square tiles) at the critical nucleus ($k^*$), is $2 - 1/k^*$ for $k \in \mathrm{N}_{>0}$ (**Supplementary Figure S5**, see section Derivation of equations below).

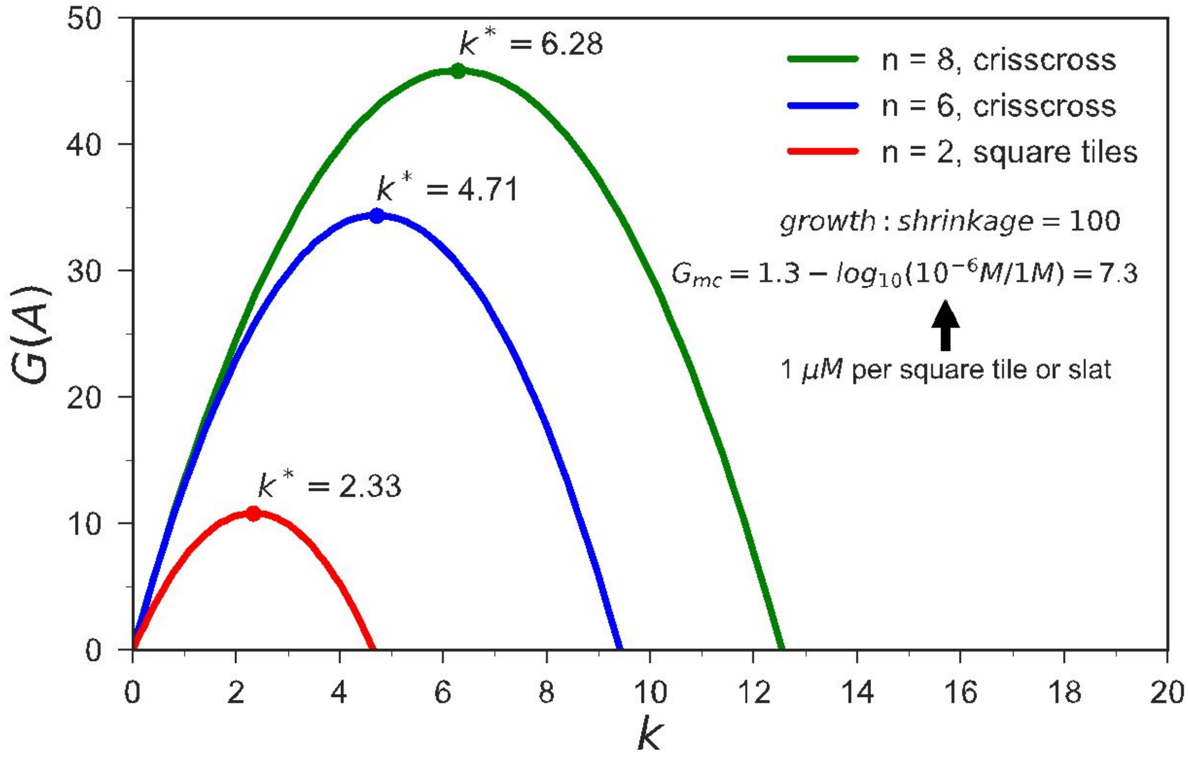


**Supplementary Figure S3**: Free energy $G(A)$ versus $k$ (i.e. $k \times k$ square-tile assembly or $2k$ crisscross assembly) for square tile ($n = 2$) and crisscross ($n = 6$ and $8$) assembly. Free energy of the monomer concentration $G_{mc} = 1.3 - log_{10}\left(\frac{10^{-6}M}{1M}\right) = 7.3$ (i.e. 1 μM each DNA slat or square tile and an effective concentration of bound slats or square tiles of 20 M (i.e. $\alpha = \log_{10}(20\ \text{M}/1\ \text{M})$) and $growth : shrinkage = 10^{\varepsilon} = 100$. Reminder from above: "Note that the irreversibility for square-tile filament growth is the ε of two-bond addition minus that for reversible growth of a filament of that width, the latter of which is nonzero due to corner tiles being captured with just a single bond. Zig-zag and tubular tile assembly systems eliminate these corner tiles through specific edge-tiles or circular arrangement (see section 4 Nucleation, figure 9, from Evans & Winfree[2])."

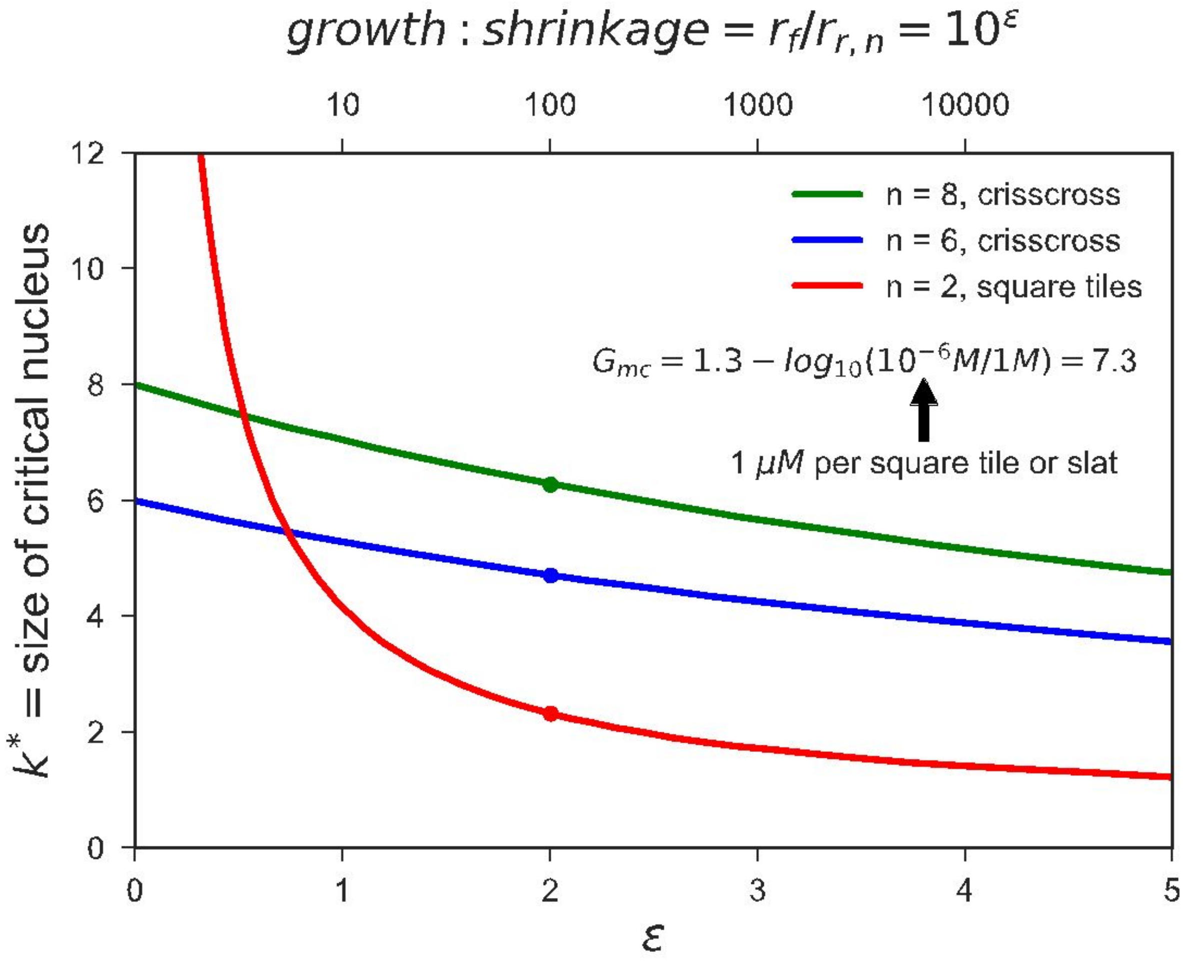


**Supplementary Figure S4**: Critical-nucleus size $k^*$ versus $\varepsilon$ for square tile ( $n = 2$ ) and crisscross ( $n = 6$ and $n = 8$ ) assembly. Free energy of the monomer concentration $G_{mc} = 1.3 - log_{10}\left(\frac{10^{-6}M}{1M}\right) = 7.3$ (i.e. 1 μM each DNA slat or square tile monomer and an effective concentration of bound slats or square tiles of 20 M (i.e. $\alpha = \log_{10}(20\ M/1\ M)$). Reminder from above: "Note that the irreversibility for square-tile filament growth is the $\varepsilon$ of two-bond addition minus that for reversible growth of a filament of that width, the latter of which is nonzero due to corner tiles being captured with just a single bond. Zig-zag and tubular tile assembly systems eliminate these corner tiles through specific edge-tiles or circular arrangement (see section 4 Nucleation, figure 9, from Evans & Winfree[2])."

**a**

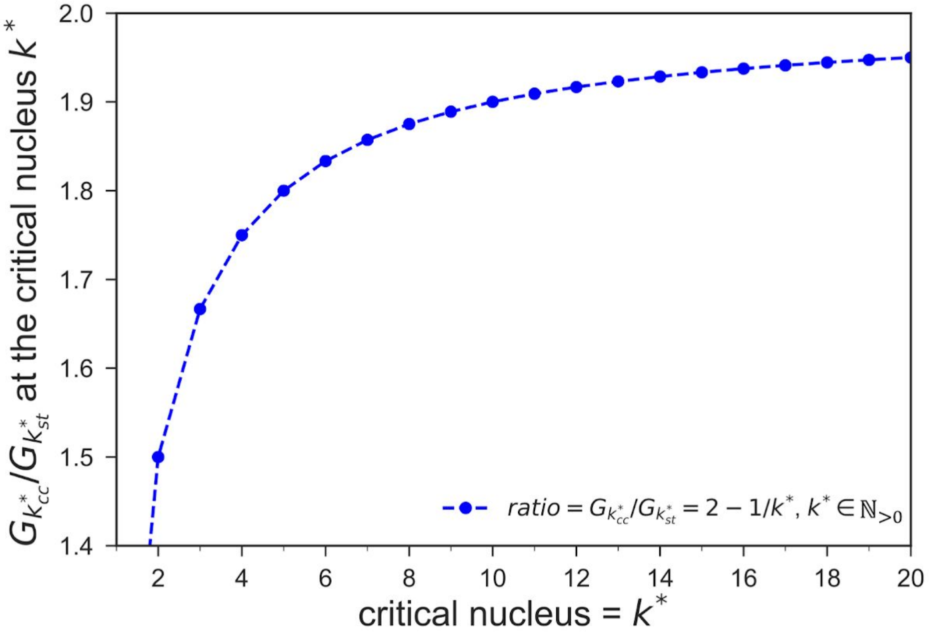


**b**

| $k^*$ | $\varepsilon_{cc}$ for $n = k^* + 1$ | $\varepsilon_{st}$ |
|---|---|---|
| 2 | 3.65 | 2.433 |
| 4 | 1.825 | 1.043 |
| 6 | 1.217 | 0.664 |
| 8 | 0.912 | 0.487 |
| 10 | 0.73 | 0.384 |
| 12 | 0.608 | 0.317 |
| 14 | 0.521 | 0.27 |
| 16 | 0.456 | 0.235 |
| 18 | 0.406 | 0.209 |
| 20 | 0.365 | 0.187 |

$\varepsilon_{cc}$ and $\varepsilon_{st}$ calculated with $G_{mc} = 7.3$

**Supplementary Figure S5**: Free-energy ratio $G_{k^*\_cc}/G_{k^*\_st}$ ($cc$ = crisscross, $st$ = square tiles) at the critical nucleus $k^*$. **a**, Plot of $G_{k^*\_cc}/G_{k^*\_st} = 2 - \frac{1}{k^*}$ is shown for $k \in \mathrm{N}_{>0}$. For large $k^*$, $G_{k^*\_cc}/G_{k^*\_st} \to 2$. **b**, ε values ($growth : shrinkage = 10^{\varepsilon}$) at critical nucleus ($k^*$) for both crisscross and square tile assembly, with $G_{mc} = 1.3 - log_{10}\left(\frac{10^{-6}M}{1M}\right) = 7.3$ (i.e. 1 μM each DNA slat or square tile monomer and an effective concentration of bound slats or square tiles of 20 M (i.e. $\alpha = \log_{10}(20\text{ M}/1\text{ M})$) and $n = k^* + 1$ for crisscross. Reminder from above: "Note that the irreversibility for square-tile filament growth is the ε of two-bond addition minus that for reversible growth of a filament of that width, the latter of which is nonzero due to corner tiles being captured with just a single bond. Zig-zag and tubular tile assembly systems eliminate these corner tiles through specific edge-tiles or circular arrangement (see section 4 Nucleation, figure 9, from Evans & Winfree[2])."

#### S1.3—Derivation of equations

##### Free energy of the assembly

Following Evans and Winfree's review[2] (see section 4 Nucleation, sentence above equation 12), the free energy of a square-tile or crisscross assembly is denoted as:

$$G(A) = NG_{mc} - BG_{se} \qquad (5)$$

Where the free energy of a "sticky end" bond is $G_{se}$, the energy from the free monomer concentration is $G_{mc}$, $N$ is the number of tiles in the assembly, and $B$ is the number of bonds formed in the assembly (**Supplementary Figure S6a,b**).

##### Square-tile assembly equations

The number of tiles (N) and bonds (B) for a $k{\times}k$ square-tile assembly (**Supplementary Figure S6a**) can be described as

$$N = k^2 \qquad (6)$$

$$B = 2k(k-1) \qquad (7)$$

Where $k \in \mathrm{N}_{>0}$. Inserting equations 6 and 7 into 5, results in the free energy

$$G(A) = k^2G_{mc} - 2k(k-1)G_{se} \qquad (8)$$

With $G_{se} = (G_{mc} + \varepsilon)/2$ , the free energy is

$$G(A) = k^2G_{mc} - k(k-1)(G_{mc} + \varepsilon) \qquad (9)$$

The critical nucleus ( $k^*$ ) is at the maximum free energy and can therefore be described as

$$k^* = \frac{G_{mc}}{2\varepsilon} + \frac{1}{2} \qquad (10)$$

**Crisscross assembly equations**

The number of tiles (N) and bonds (B) for crisscross assembly (**Supplementary Figure S6b**) can be described as

$$N = 2k \tag{11}$$

$$B = k^2 \tag{12}$$

Where $k \in N_{>0}$. Inserting equations 11 and 12 into 5, results in the free energy

$$G(A) = 2kG_{mc} - k^2 G_{se} \tag{13}$$

With $G_{se} = (G_{mc} + \varepsilon)/n$, where $2n$ is the number of bonds that a crisscross slat can form (**Supplementary Figure S6c**), the free energy is

$$G(A) = 2kG_{mc} - k^2 \frac{(G_{mc}+\varepsilon)}{n} \tag{14}$$

The critical nucleus ( $k^*$ ) is at the maximum free energy and can therefore be described as

$$k^* = \frac{n}{1+\frac{\varepsilon}{G_{mc}}} \tag{15}$$

**Crisscross:square tiles ratio of maximum free energy**

The maximum free energy for crisscross (cc) and square tile (st) assembly is at the critical nucleus ($k^*$) and given by equations 16 and 17

$$G_{k^*\_cc} = 2k^* G_{mc} - k^{*2} G_{mc} \frac{1}{n} \frac{n}{k^*}$$

$$G_{k^*\_cc} = k^* G_{mc} \tag{16}$$

$$G_{k^*\_st} = k^{*2} G_{mc} - k^*(k^* - 1) G_{mc} \left(1 + \frac{1}{2k^*-1}\right)$$

$$G_{k^*\_st} = G_{mc} \frac{k^{*2}}{2k^*-1} \tag{17}$$

Subsequently the ratio of $G_{k^*\_cc}/G_{k^*\_st}$ results in

$$G_{k^*\_cc}/G_{k^*\_st} = 2 - \frac{1}{k^*} \qquad (18)$$

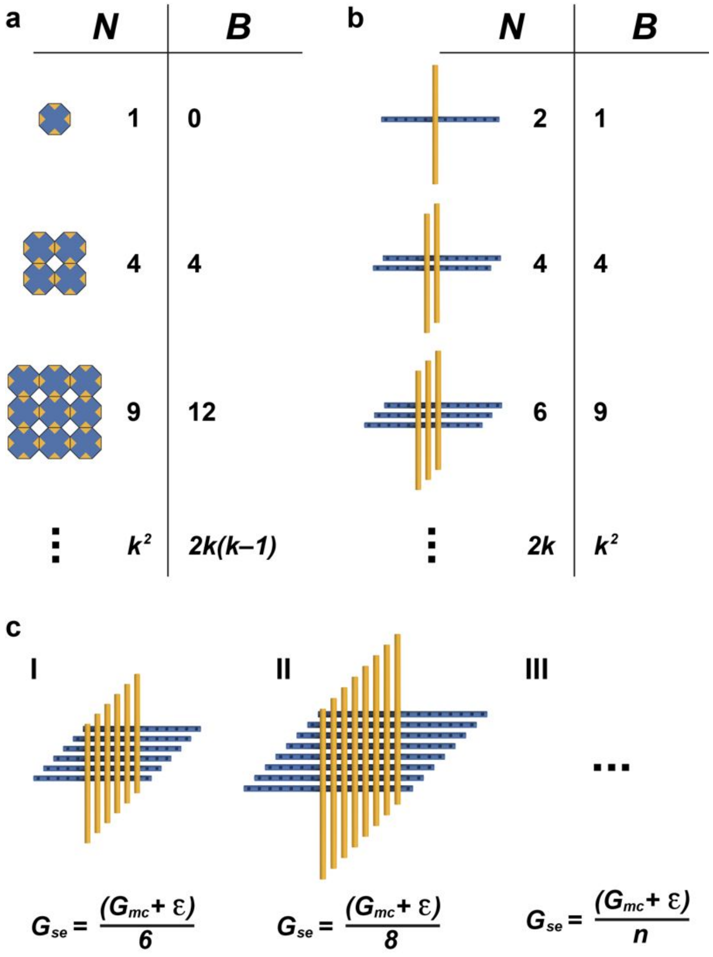


**Supplementary Figure S6**: Critical nuclei for square tile and crisscross assembly (figure adapted from section 4 Nucleation, figure 8, from Evans and Winfree[2]). **a**, Formation of a critical nucleus for square tile assembly shown as number of tiles (N) and bonds (B) formed. **b**, Formation of a critical nucleus for crisscross assembly shown as number of tiles (N) and bonds (B) formed. **c**, Relationship between $G_{se}$ and $G_{mc}$ and the size of the maximum critical nucleus ($k^*$) depending on the DNA slat length.

#### S2—ssDNA slat architecture

**Overview:** Here we clarify how the abstract cartoon representation of the slats and seed (as shown in **Figure 1a**, **1c**, and **Supplementary Information section S1**) was implemented experimentally as ssDNA slats in this work. **S2.1** shows ball-and-stick DNA models for the critical nuclei of the v6 and v8 designs, **S2.2** illustrates the design and successful folding of the DNA origami seed, and **S2.3** explains how sequence symmetry of the DNA slats was used so that sets of slats periodically bind to perpetuate ribbon growth.

##### S2.1—Critical nuclei comprised of ssDNA slats

###### *S2.1.1—v6 critical nucleus*

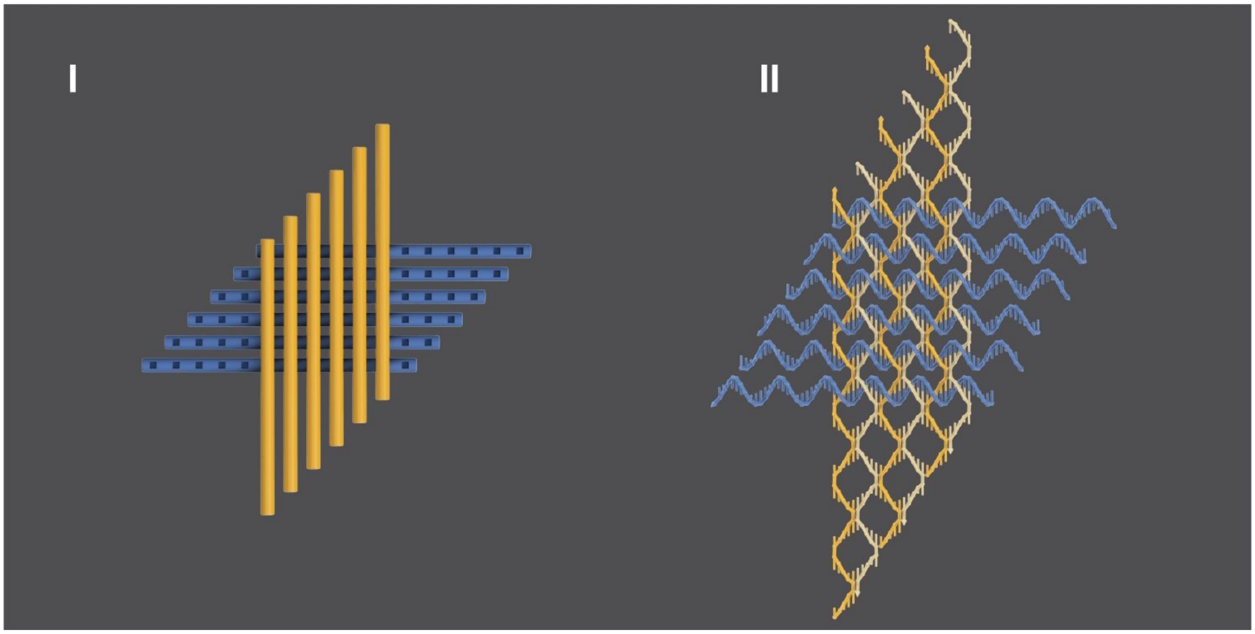


**Supplementary Figure S7**: Critical nucleus for $n$=6 (v6) sequence variants is composed of 12 individual slats. **I**, generalized cartoon model. **II**, specific implementation for ssDNA slats as shown with ball-and-stick DNA model. Each slat blue x-slat or gold y-slat is comprised of 12 half-turns of DNA, with all binding sites as shown in this particular model as 5 nt. However, sequences tested in this work also contained 6 nt binding sites.

##### *S2.1.2—v8 critical nucleus*

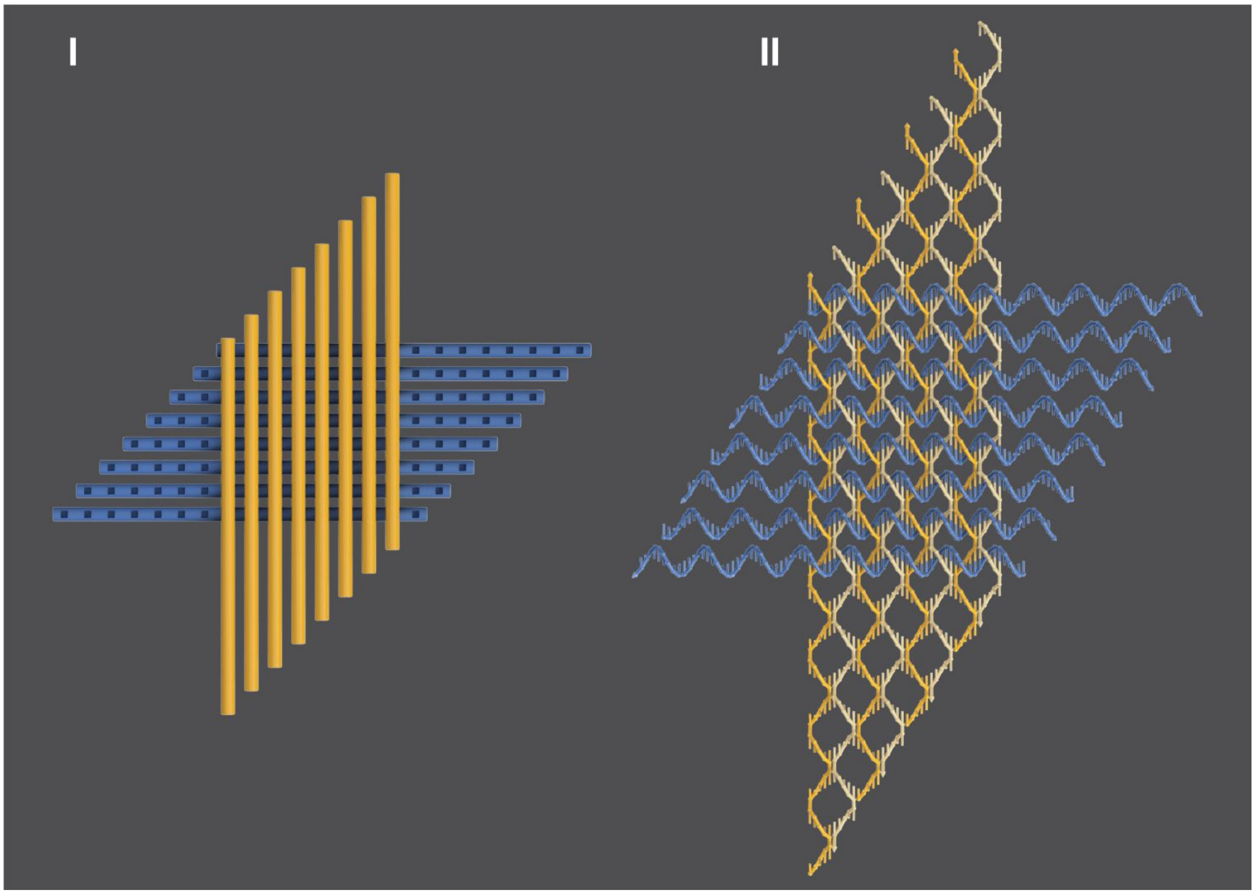


**Supplementary Figure S8**: Critical nucleus for *n*=8 (v8) sequence variants is composed of 16 individual slats. **I**, generalized cartoon model. **II**, specific implementation for ssDNA slats as shown with ball-and-stick DNA model. Each slat blue x-slat or gold y-slat is comprised of 16 half-turns of DNA, with all binding sites as shown in this particular model as 5 nt. However, sequences characterized in this work also contained 6 nt binding sites.

#### S2.2—DNA-origami seed design and characterization

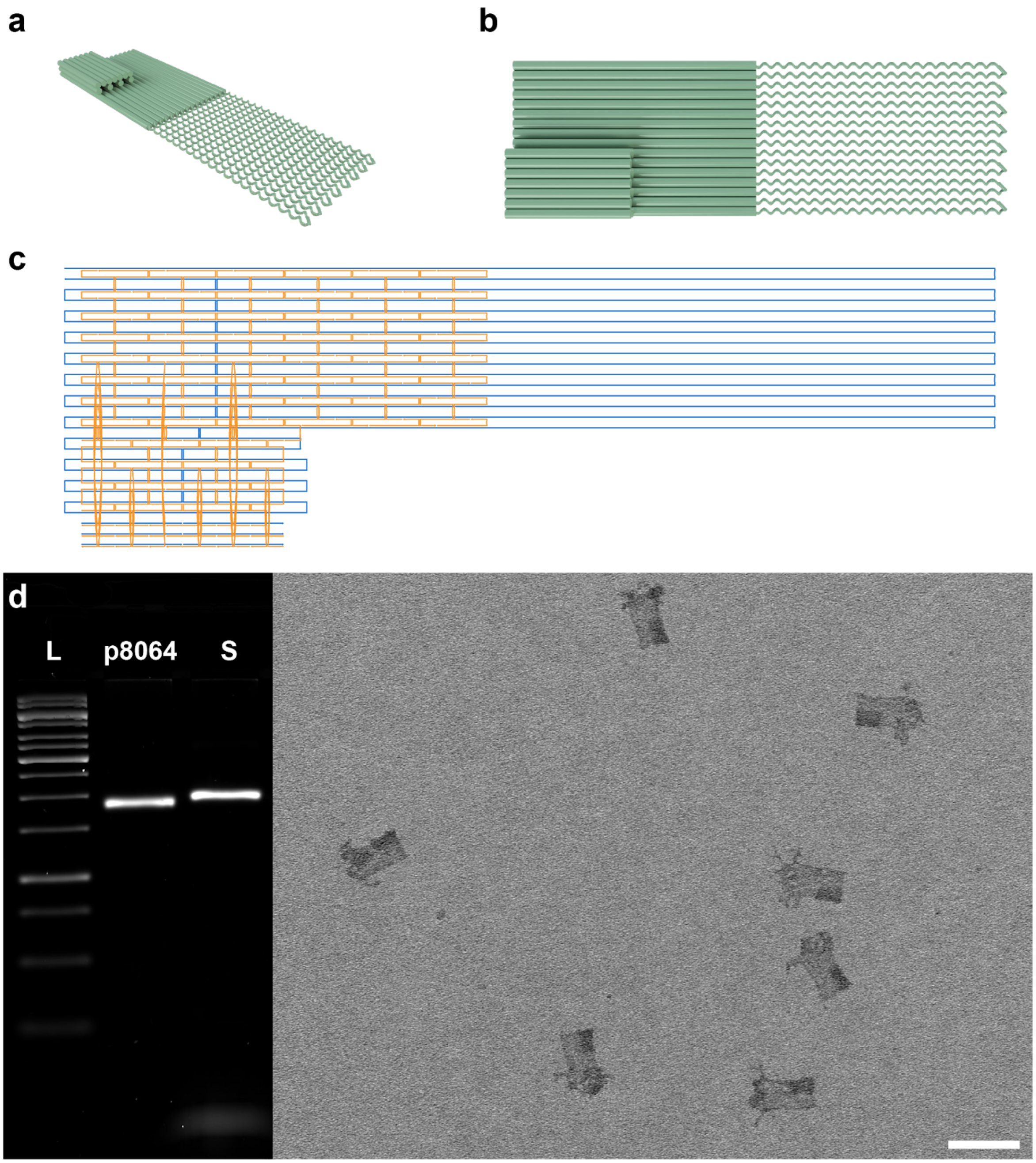


**Supplementary Figure S9**: DNA origami seed design and assembly. **a** and **b**, rendering of cylindrical extrusion of design from corner and top views respectively. Sixteen bare scaffold DNA loops that bind the nuc-y-slats are shown as twisted thin cylinders, versus the dsDNA body and reference square shown as straight thick cylinders. **c**, DNA origami scaffold routing diagram as generated with caDNAno[9] with the bare scaffold shown in the right-half of the diagram. **d**, Agarose-gel characterization (left) of folded DNA origami seed (S), with 1 kb Ladder (L), and M13 p8064 scaffold. TEM micrograph (right) of folded DNA origami seeds. Note that bare ssDNA scaffold loops appear unstructured and twisted. Scale bar is 100 nm.

##### S2.3—Sequence design of DNA slats

DNA slat sequences for v6 and v8 were designed using custom Python scripts (available upon request). We assessed base-pairing energies using Unafold[10] and designed all sequences to have minimal self-structure. We further designed DNA slat sequences (shown in **Supplementary Table S1**) to have stronger base-stacking in the y-direction than in the x-direction (using Unafold[10] or base-stacking energies reported in Protozanova et al[11]), which resulted in more uniform length distributions of assembled ribbons, coupled with greater spurious nucleation (see section **S3.2** for experimental results). This behavior was puzzling to us, as we initially hypothesized that stronger base-stacking in the x-direction than in the y-direction would give rise to more robust growth, and yet we observed the opposite trend (not shown). Currently we are unable to rationalize the basis for these trends. It is noteworthy that we were not able to resolve helical directionality using our negative-stain TEM imaging conditions; this difficulty in discriminating individual helices may arise from the dense crossover patterns within the ribbons holding those helices very close together. As a consequence, we were unable to resolve whether the stacking direction has the ability to switch spontaneously along the length of ribbons[12] or whether the intended design of stronger base-stacking in y- over x-direction actually gave rise to a preferred stacking direction in the assembled ribbons. In the future, this should be resolvable by designing a contrast-heavy marker feature into the ribbon structure that clearly reports on the directionality of the helices in negative-stain TEM. Looking to the future, we believe that tuning base-stacking energies is one of potentially many parameters that can be optimized for DNA slat sequences. However, due to time and cost limitations we were unable to explore further the large DNA slat sequence design space in this initial study.

| DNA slat sequence variant | stronger base-stacking in y- over x-direction |
|---|---|
| 6.1 | Yes |
| 6.2 | No |
| 6.3 | Yes |
| 8.1 | Yes |
| 8.2 | No |

**Supplementary Table S1**: DNA slat sequence variants designed to have stronger base-stacking in the y-over x-direction.

#### S2.4—Sequence symmetry of DNA slats versus number of repeating slats

Growth of ribbons is perpetuated by sequential binding of repeating sets of x and y-slats, as introduced in **Figure 1a** and **1d**. The domains (i.e. ssDNA sequences) overhanging from slats at the terminus of a growing ribbon determine the specific slats that bind next. The number of repeating slats in the set for a given variant is arbitrary and programmed into the sequence design of the slats themselves. We exploited sequence symmetry in certain variants so that ribbon growth was attainable with fewer unique x- and y-slats, as shown in **Supplementary Table S2**. We were interested in decreasing the number of unique slats per design because purchasing fewer slats per design tested made it more economically feasible to test a larger number of designs. Symmetry, in context of a given x- or y-slat, refers to the number of copies of a specific perpendicular slat to which it is bound in the final ribbon. The "low" sequence symmetries for v6 and v8 had 12 and 16 unique pairs of x- and y-slats, respectively. For v8, we also tested "medium" and "high" sequence symmetry so that that ribbon growth could be attained with 8 and 4 unique pairs of x- and y-slats, respectively.

| Relative symmetry | Sequence variants | Cartoon unit cell | Domain layout in example slat |
|---|---|---|---|
| **Low**<br>12 x-slats<br>12 y-slats | **v6.1**<br>**v6.2**<br>**v6.3** | 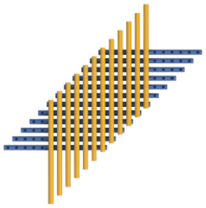 | 1 2 3 4 5 6 7 8 9 10 11 12 |
| **Low**<br>16 x-slats<br>16 y-slats | **v8.2** | 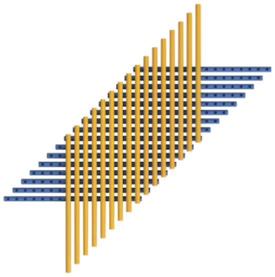 | 1 2 3 4 5 6 7 8 9 10 11 12 13 14 15 16 |
| **Medium**<br>8 x-slats<br>8 y-slats | **v8.1**<br>**v8.3**<br>**v8.4**<br>**v8.5**<br>**v8.6** | 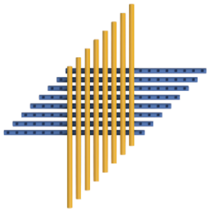 | 1 2 3 4 5 6 7 8 1 2 3 4 5 6 7 8 |
| **High**<br>4 x-slats<br>4 y-slats | **v8.7** | 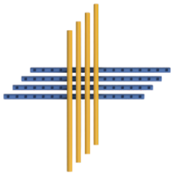 | 1 2 3 4 1 2 3 4 1 2 3 4 1 2 3 4 |

Unique domain

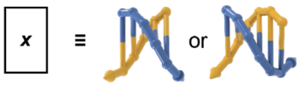


**Supplementary Table S2**: Sequence symmetry of specific variants for v6 and v8 designs. Increasing the sequence symmetry allows growth to be attained with fewer unique x- and y-slats. The concept of symmetry in slat design is illustrated in the rightward domain layout diagram. Each boxed cell is a unique half-turn binding sequence that is programmed to bind a specific perpendicular slat. The numbering '*x*' in each box refers to some specific complementary slat in the opposing orientation.

#### S3—Determining optimal growth conditions for seeded ribbons

**Overview:** In **S3**, we describe the characterization of experimental conditions for seeded growth of ribbons with ssDNA slats. This section does not consider spurious assembly of ribbons from just the slats themselves. In **S3.1**, we show additional TEM images of seeded v6 and v8 ribbons. Then, we use agarose gels in **S3.2** to determine optimal isothermal temperature and $MgCl_2$ parameters for seeded growth with either 0.2 or 1 µM each slat. We further show that the copy-number of ribbons is linearly controllable by the number of seeds added (see **S3.3**) and that blockage of nuc-y strands is a strong impediment to growth (see **S3.4**).

##### S3.1—Additional TEM images of v6 and v8 ribbons

###### *S3.1.1—v6.1 ribbons*

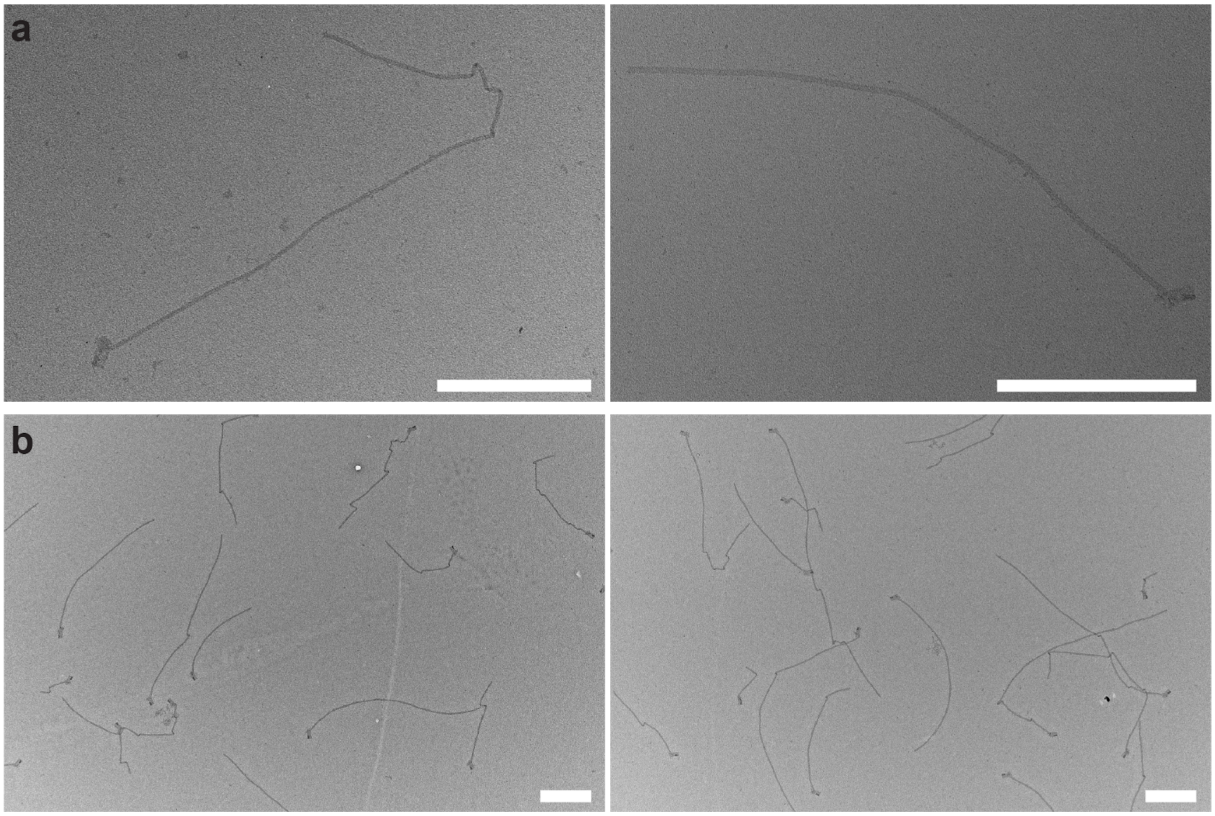


**Supplementary Figure S10**: TEM images depicting seeded growth of v6.1 ribbons. Growth was conducted for ~16 hours at 50°C, with 14 mM $MgCl_2$, 2 nM seed, and 0.2 µM each slat. Seeds are visible on all structures shown. **a**, Close-up of single ribbons. Left ribbon is ~2.3 µm long, and the right ribbon ~1.5 µm long. **b**, Lower magnification shows population of ribbons. Scale bars are 500 nm.

##### *S3.1.2—v8.2 ribbons*

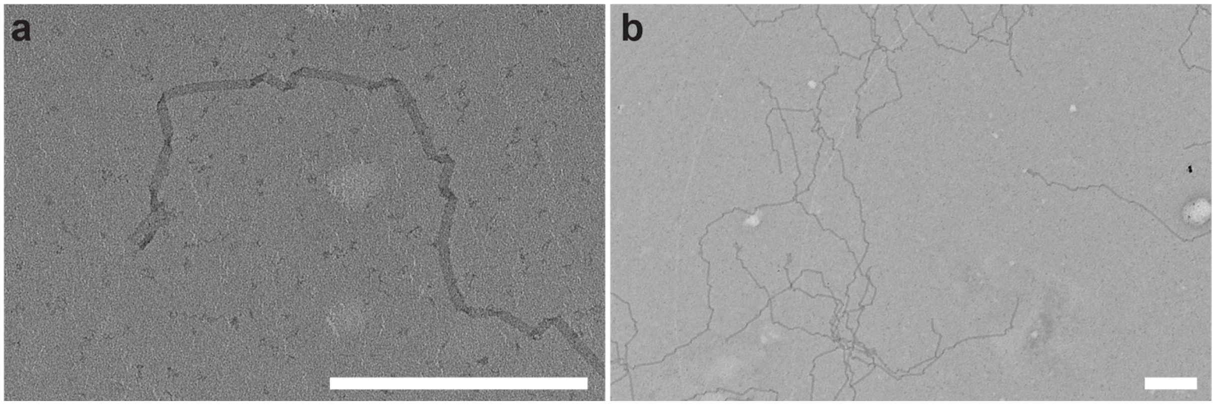


**Supplementary Figure S11**: TEM images depicting seeded growth of v8.2 ribbons. Growth was conducted for ~16 hours at 50°C, with 16 mM MgCl2, 2 nM seed, and 0.5 µM slats. Seeds are generally visible on structures shown. **a**, Close-up of single ribbon with ~1.6 µm length of growth shown. **b**, Lower magnification shows population of overlapping ribbons. Scale bars are 500 nm.

##### *S3.1.3—v8.7 high-sequence symmetry ribbons*

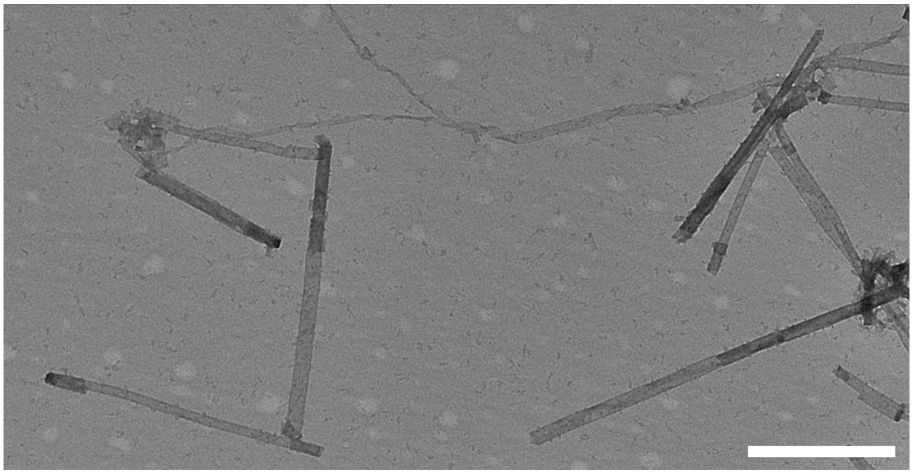

**Supplementary Figure S12**: TEM images depicting seeded growth of the high symmetry v8.7 slats. These ribbons are prone to blunt-end stacking because they lack T-brush passivation on the ends of the slats, and are thus forming tubes and aggregates. This particular design uses 10 bp/turn, which also alters ribbon morphology (see **S8** for more details on the formation of tubes). Scale bar is 500 nm.

#### S3.2—Growth of ribbons under optimal conditions

##### *S3.2.1—v6 designs*

###### *S3.2.1.1—v6.1, v6.2, v6.3 0.2 µM each slat*

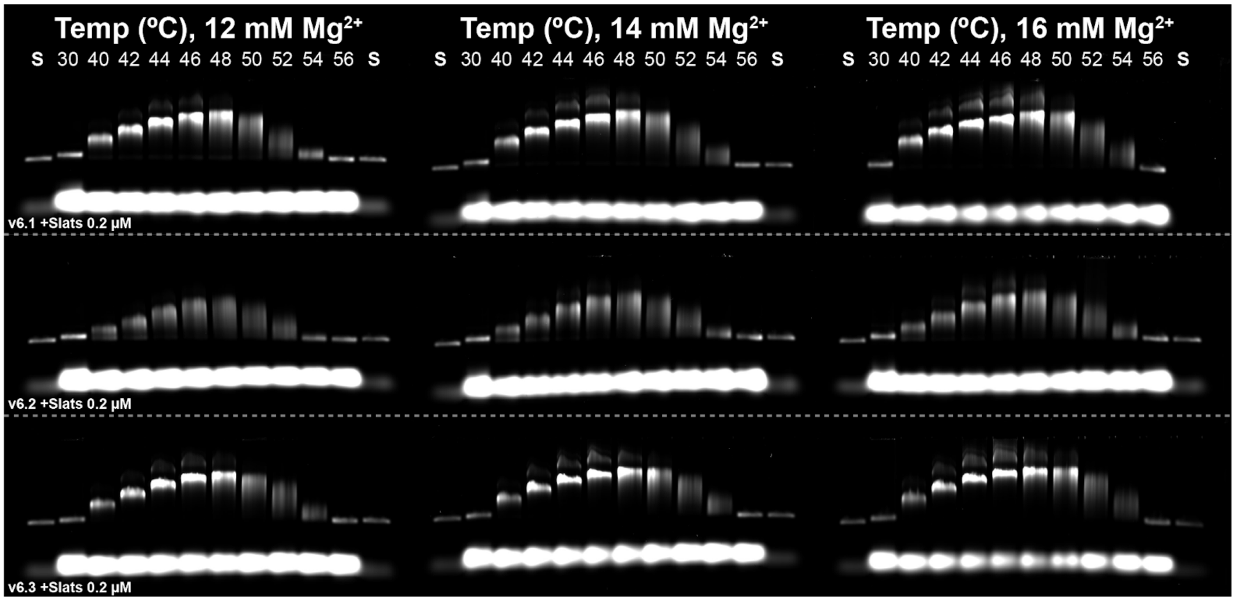


**Supplementary Figure S13:** Agarose-gel characterization of seeded growth of v6.1, v6.2, and v6.3 to find optimal seeded growth temperatures and $MgCl_2$. Growth was conducted for ~16 hours at temperature listed above each lane, with either 12, 14, or 16 mM $MgCl_2$. Seed was at 2 nM and slats at 0.2 µM each. Lane (S) is the seed only control. Note that the overall slat population was significantly depleted for v6.3 16 mM $MgCl_2$ at the optimal temperature, even though the nuc-y-slats (i.e. ⅓ of the added total) could not be consumed.

*S3.2.1.2—v6.1 1 µM each slat*

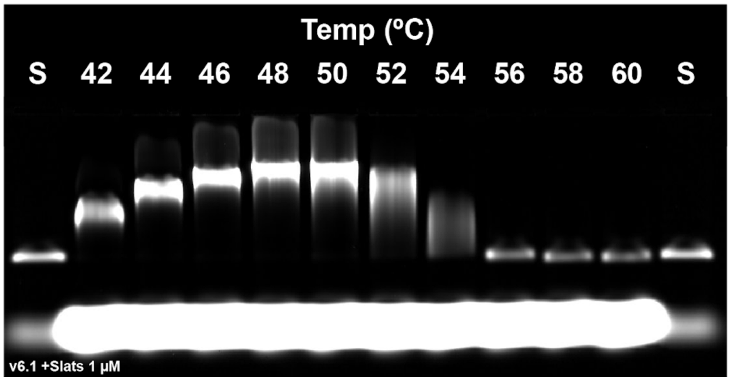


**Supplementary Figure S14:** Agarose-gel characterization of seeded growth of v6.1 to find optimal seeded growth temperature with 1.0 µM each slat. Growth was conducted for ~16 hours at temperature listed above each lane, with 16 mM $MgCl_2$. Seed was at 2 nM and slats at 1 µM. Lane (S) is the seed only control.

***S3.2.2—v8 designs***

*S3.2.2.1—v8.2 0.2 µM each slat*

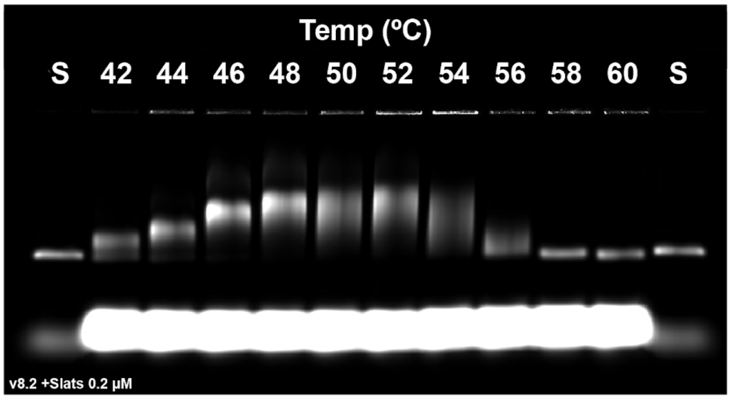


**Supplementary Figure S15:** Agarose-gel characterization of seeded growth of v8.2 to find optimal seeded growth temperature with 0.2 µM each slat. Growth was conducted for ~16 hours at temperature listed above each lane, with 14 mM $MgCl_2$. Seed was at 2 nM and slats at 1 µM each. Lane (S) is the seed only control.

*S3.2.2.2—v8.1 1 μM each slat*

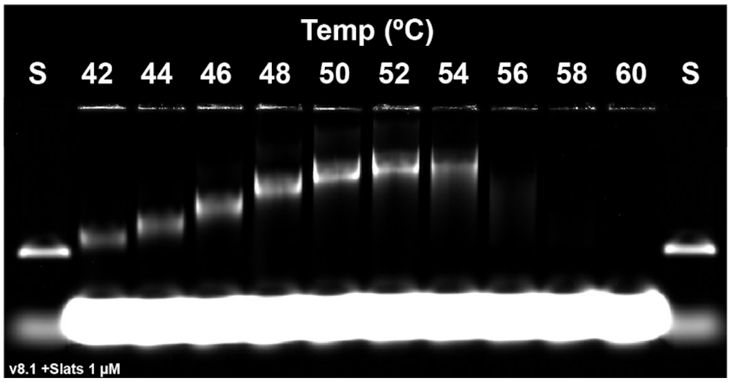


**Supplementary Figure S16:** Agarose-gel characterization of seeded growth of v8.1 to find optimal seeded growth temperature with 1.0 μM each slat. Growth was conducted for ~16 hours at temperature listed above each lane, with 16 mM $MgCl_2$. Seed was at 2 nM and slats at 1 μM each. Lane (S) is the seed only control. Long filaments of this design appear to aggregate at temperatures above 54ºC; presumably this could be mitigated with even longer T-brushes for greater passivation.

#### S3.3—Copy number control of ribbons with the seed

We expect the copy number of ribbons in final reaction to be linearly proportional to the number of seeds added to the starting reaction. This property of DNA slats is analogous to DNA origami in how the number of scaffold strands precisely controls the copy number of folded structures. In **Supplementary Figure S17**, we show experimental validation of this concept by assembling the v6.1 ribbons with a three-fold dilution series of seed at a temperature where spurious nucleation was not observed.

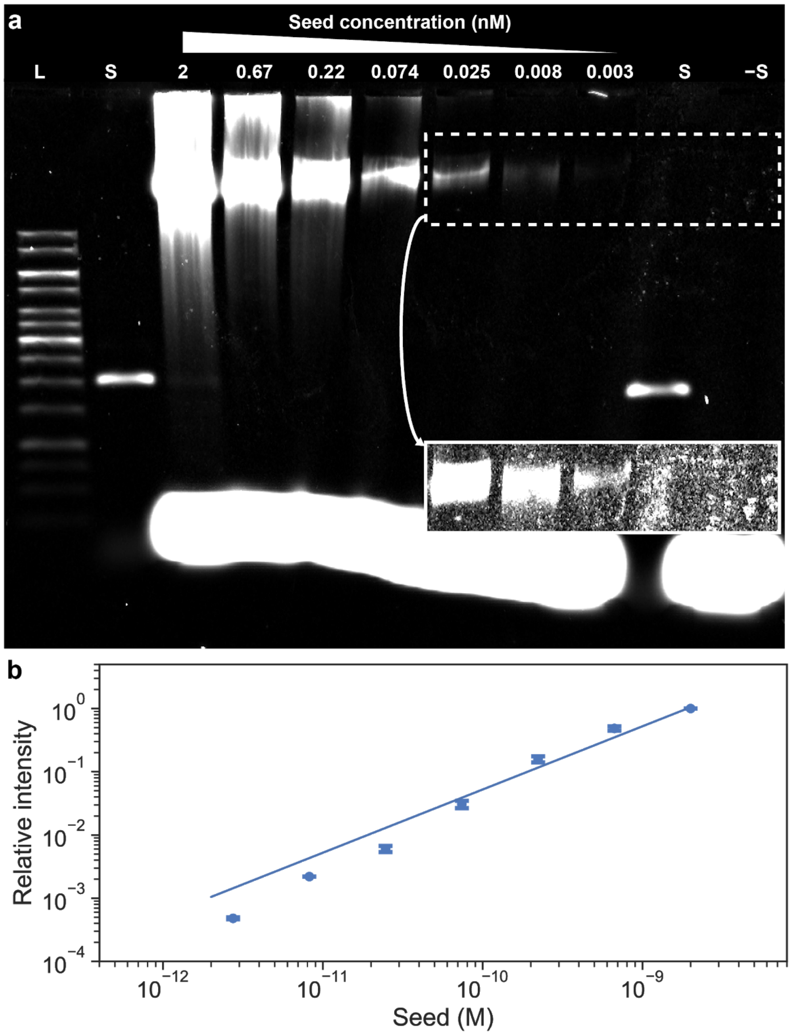


**Supplementary Figure S17:** Growth of v6.1 ribbons with different initial concentrations of seed. Growth was conducted for ~12 hours at 50°C, with 16 mM $MgCl_2$ and slats at 1 μM each. Insert shows contrast and brightness adjusted gel for better visualization. **a**, Agarose gel of single replicate of experiment. Lane (L) is the ladder and (S) is the seed only control. **b**, Plot of relative gel densitometry (measured with respect to the conditions with the highest 2 nM concentration of seed) versus initial concentration of seed, with the line showing a linear fitting of the data. Each data point is the mean of a duplicate experiment. Error bars are ±SD.

##### S3.4—Blocking nuc-y slats impedes ribbon assembly

We hypothesized that ribbon assembly requires sequential addition of every single designed slat for growth. To test whether ribbon growth could be attained if specific slats in assembly mixture are impeded, we added and 5-fold excess of blocking strands (i.e. reverse complement sequences of a slat) to mask one or more specific nuc-y slats. In **Supplementary Figure S18**, we show that blockage of only a single nuc-y slats completely terminated ribbon growth with 0.2 μM each slat. Using higher concentration (i.e. 1 μM each) slats allowed limited recovery of growth (i.e. skipping over the impeded nuc-y slats) for when up to two nuc-y slats were blocked, although growth was completely terminated when three nuc-y slats were blocked.

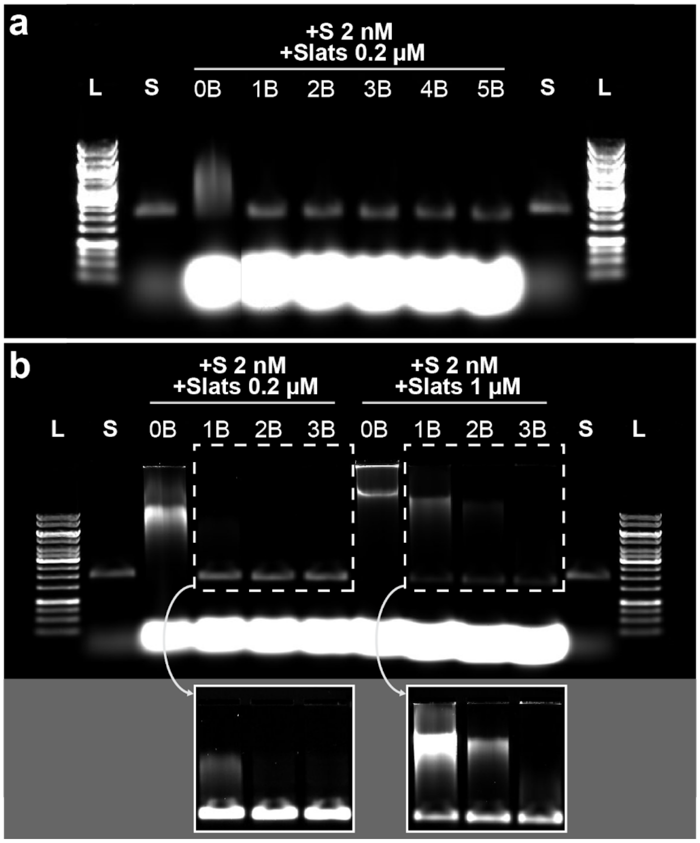

**Supplementary Figure S18:** Seeded DNA slats assembly with sequence v6.1 and blocking strands (B) added at 5X molar excess under various assembly conditions. **a**, DNA slats assembly with 0.2 µM each slat and 2 nM seed, in 12 mM $MgCl_2$ at 50ºC for ~16 hours. No blocking strand (0B) and 1–5 blocking strands (1B–5B, i.e. blocking 1, 2, 3, 4, or 5 nuc-y-slats simultaneously), with each at 1 µM concentration. Only 0B displayed visible growth. **b**, Increasing concentration of the slats from 0.2 µM each (left) to 1 µM (right) with 2 nM seed, in 14 mM $MgCl_2$ at 45ºC for ~16 hours. Blocking strand were added at either 1 µM (left) or 5 µM (right). The combination of lower temperature, higher concentration of slats, and more $MgCl_2$ in the right-half of *b* was sufficient to allow partial recovery of ribbon growth for up-to blockage of two distinct nuc-y strands, as shown in the cut-away contrast and brightness adjusted section of the gel. (L) 1 kb ladder and (S) seed only control.

#### S4—Assay for spontaneous nucleation under optimal growth conditions

**Overview:** In **S4**, we describe the measured amount of spurious ribbon assembly under optimal growth conditions when no seed was added to the reaction, as a function of the slat design. We challenged the DNA slats by carrying out growth with higher 1 μM each slat concentrations using the optimal growth conditions from **Figure 2 a−b** and **S3**. We show in **S4.1** that the SYBR-gold detection limit of ribbons is ~200 fM. We found that v6.1 (**Figure 2c**), v6.2 (**S4.2**), and v8.1 and v8.2 (**S4.4**) do not show any observable spuriously formed ribbons on gels after overnight incubation. We also show that sequence variant v6.3 (**S4.3**) exhibits spurious growth for selected conditions after overnight incubation, indicating that certain sequences are more prone to spurious assembly than others. Furthermore, we found limited to no evidence of spurious ribbon formation even after prolonged 100 hour incubation of 1.0 μM each slat with 16 mM $MgCl_2$ for v6.1 and v6.2 (**S4.5**): v6.1 showed a faint gel band near the SYBR-gold detection limit at the lowest 46°C temperature tested, versus v6.2 which showed no assembly. Finally, we find that the reversible temperature for slat binding with v6.1 and v6.2 is ~56.2°C under these salt and slat concentrations (**S4.6**). Taken together, this section shows that strictly seed-initiated ribbon assembly, as measurable to the 200 fM gel detection limit, can be attained with high 1 μM concentrations of each slat and high 16 mM $Mg^{2+}$ concentrations at growth temperatures that are ~10°C below the reversible temperature.

##### S4.1—SYBR-Gold gel detection limit of ribbons

To determine the sensitivity limit of SYBR-Gold, we loaded a dilution series of ribbons on agarose gels to determine the minimal concentration where a gel band could be identified. We prepared a typical ribbon assembly reaction (1 μM each v6.1 slat, 16 mM $MgCl_2$, 50ºC for ~16 hr) where 2 nM initial concentration of seed was added. The ribbons were diluted 1 in 250 in 1X reaction buffer. Then, a Labcyte Echo 525 acoustic liquid handler was used to transfer volumes (from 0.025–6.4 μL) of diluted reaction into 10 μL pools of agarose gel loading buffer. The volumes transferred corresponded to the absolute number of ribbons as labeled above each gel well in **Supplementary Figure S19a**, where we assumed one-for-one conversion of each seed into a ribbon. Given the typical 4 μL loading volume of assembly reactions for gel characterization, the absolute loading amounts correspond to the fM molarity annotated above each gel well. A faint, barely discernible band is seen for 200 fM (~$4.8*10^5$ ribbons). We further use the assumption that the mean length of ribbons in the reaction to be ~5 μm based on TEM observations and the model in **Figure 2d**, such that the picogram mass can be estimated. We assume each slat contributes 1.5 nm to the total length of the ribbon, such that there are an average of ~3333 slats (with each slat comprising 71 nt) extending from each 8064 bp seed. Thus, we calculate that we can detect ribbons to about 65 pg of assembled material (i.e. ~200 fM ribbons in 4 µL reaction). Triplicate gel results showing the gel densitometry versus the pg mass and fM amounts of ribbon are shown in **Supplementary Figure S19 b–c** respectively.

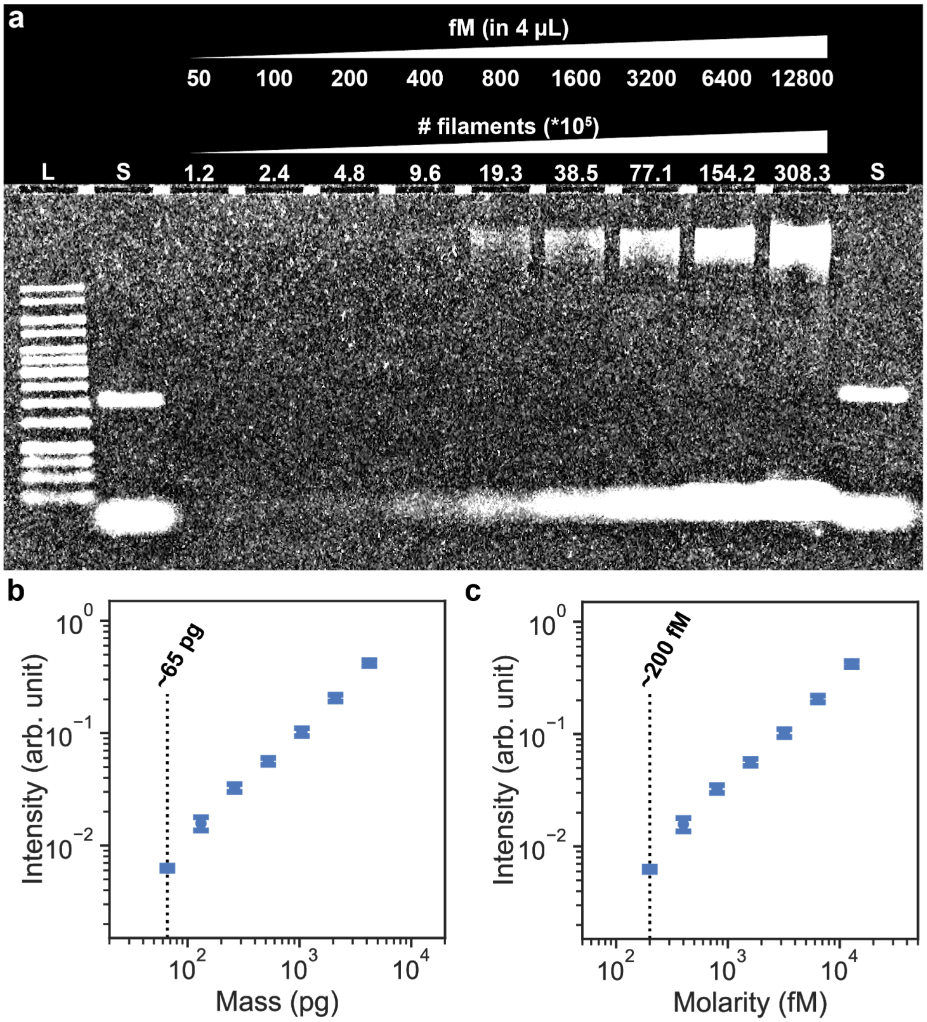


**Supplementary Figure S19**: Determining the sensitivity limit of SYBR-Gold agarose gel detection in our gel setup with a dilution series of ribbons. **a**, Agarose gel of a single dilution series, where (L) is the ladder and (S) the seed control, with a faint barely discernible band seen for 200 fM. **b–c**, Plots of a triplicate gel dilution series experiment showing mean gel intensities vs estimated mass and molarity of ribbons loaded, with error bars showing ±SD. The SYBR-Gold detection limits are labeled with a dotted line on each plot.

#### S4.2—Spurious nucleation not observable for v6.2 with optimal growth conditions

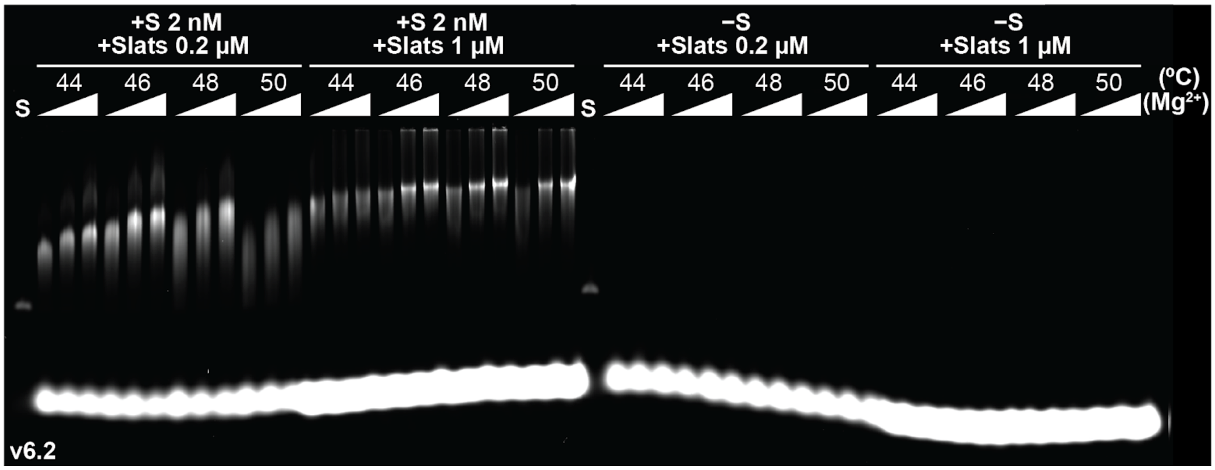


**Supplementary Figure S20**: Gel characterization of seeded (left) and unseeded (right) growth of DNA v6.2 slats. Reactions with either 0.2 or 1.0 µM each slat were incubated isothermally at one of four temperatures in either 12, 14, or 16 mM $MgCl_2$ for ~16 hours. Notably, unseeded reactions yielded no observable assembly at all conditions tested. (S) is the seed only control.

#### S4.3—Spurious nucleation is observable for v6.3 under select optimal growth conditions

##### *S4.3.1—Gel characterization of spurious nucleation of v6.3*

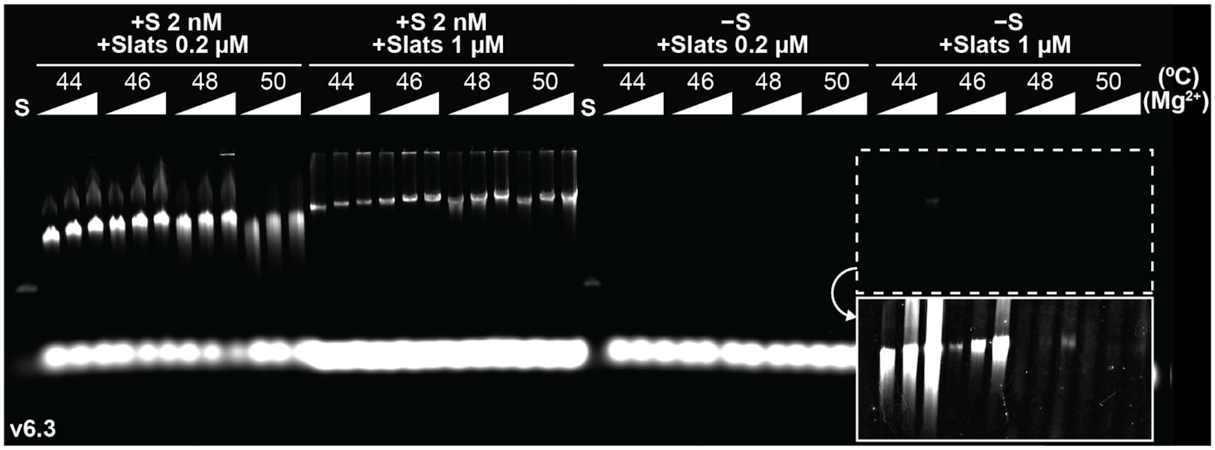

**Supplementary Figure S21**: Gel characterization of seeded (left) and unseeded (right) growth of DNA v6.3 slats. Reactions with either 0.2 or 1.0 µM each slat were incubated isothermally at one of four temperatures in either 12, 14, or 16 mM $MgCl_2$ for ~16 hours. Unseeded reactions yielded no observable assembly with 0.2 µM each slat. However, unseeded reactions with 1.0 µM each slat yielded some observable assembly as shown in the contrast and brightness adjusted cutaway from the gel. (S) is the seed only control.

##### *S4.3.2—TEM validation that spuriously formed v6.3 gel bands are indeed ribbons*

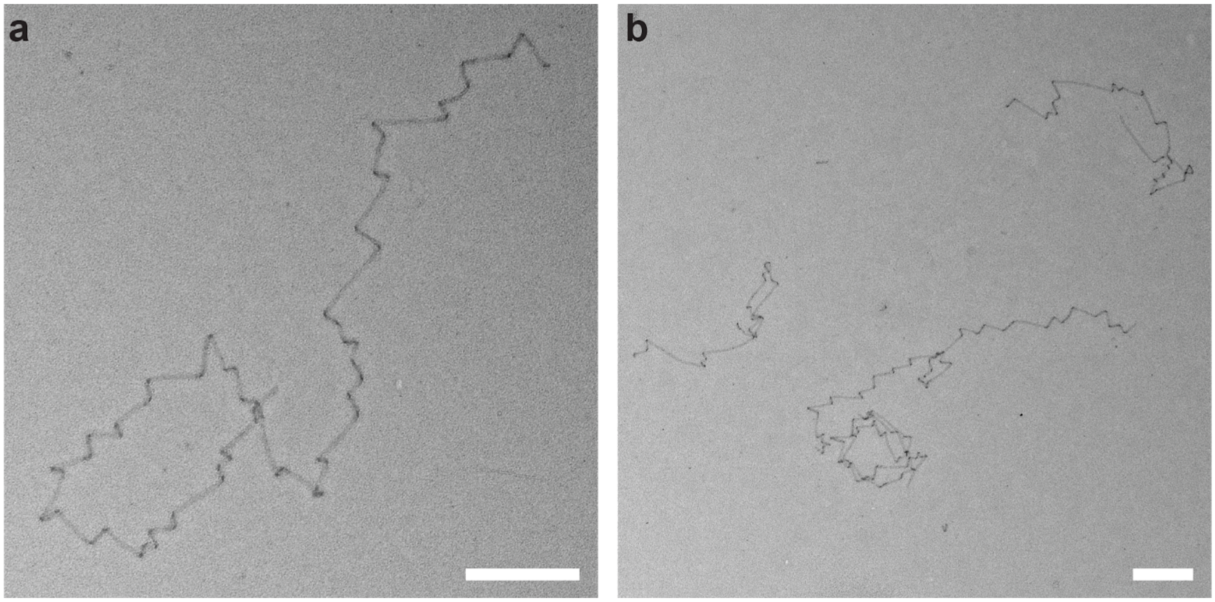


**Supplementary Figure S22**: **a** and **b**, TEM images of the spuriously formed gel band for v6.3 showing that the assemblies are well-formed ribbons, as opposed to some other undesirable artifact. No seed was observed on any of the ribbons. The band from the gel in **Supplementary Figure S21** for one condition (−seed, 16 mM $MgCl_2$, 1 µM each slat, 44ºC) was trimmed with a razor blade, crushed, and deposited onto a TEM grid. Scale bars are 500 nm.

##### S4.4—Spurious nucleation is not observable for v8.1 and v8.2 under optimal growth conditions

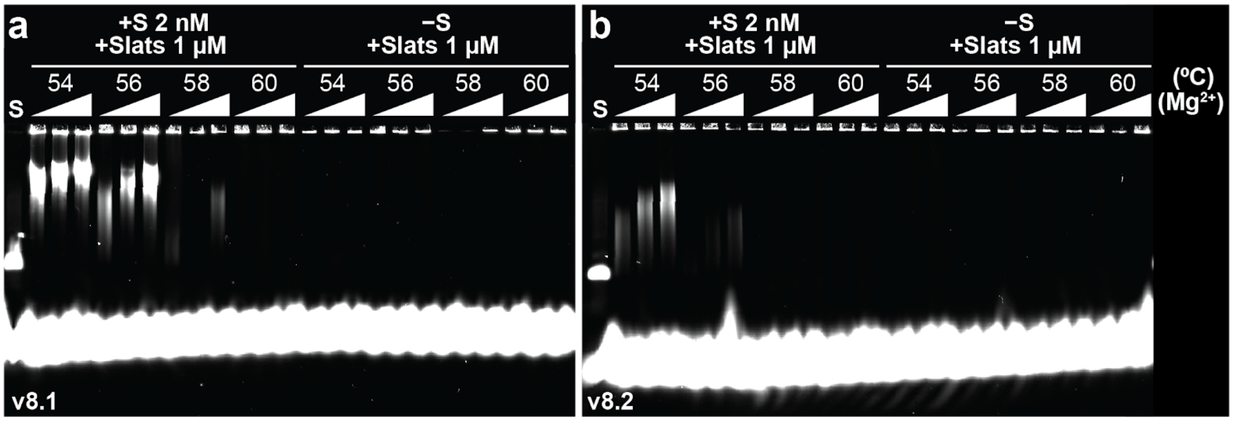


**Supplementary Figure S23**: Gel characterization of seeded and unseeded growth of DNA slats for v8.1 and v8.2 in **a** and **b** respectively. Reactions with 1 µM each slat were incubated isothermally at one of four temperatures in either 12, 14, or 16 mM $MgCl_2$ for ~16 hours. Reactions without a seed yielded no observable assembly at all conditions tested. (S) seed only control.

##### S4.5—Extended 100 hour incubation of v6.1 and v6.2 under optimal growth conditions

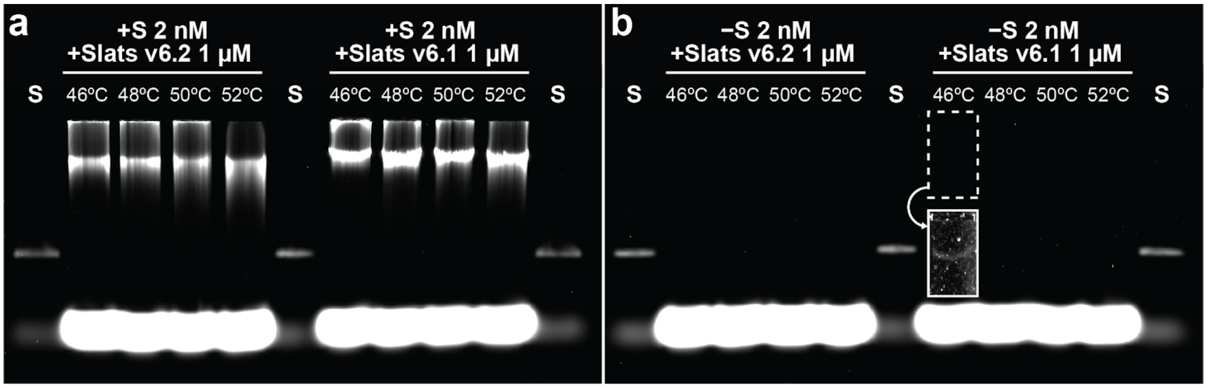


**Supplementary Figure S24**: Gel characterization of v6.2 and v6.1 slats with prolonged 100 hour incubation at one of four temperatures with 16 mM $MgCl_2$. **a**, Seeded reactions containing 1 µM each slat and 2 nM seed with ribbon growth at all conditions. **b**, Unseeded reactions containing 1 µM each slat. Notably, unseeded growth was only noted for v6.1 at the lowest temperature (46ºC) tested, with a faint band near the SYBR-Gold detection limit. See **S6.3** for more detailed analysis using mathematical modelling.

#### S4.6—Determination of reversible temperature for binding v6.1 and v6.2 slats

We determined the approximate reversible temperature for slat binding (for v6.1 and v6.2) at 16 mM $MgCl_2$ and 1 μM each slat by agarose gel in **Supplementary Figure S25**. Ribbons were grown for 1 hour at 48ºC, and then the temperature was increased for 3.5 hours across a gradient ranging 52–59ºC to find the melt temperature where growth ends and ribbons fall apart. Gel bands for the ribbons were compared with respect to a control grown for 1 hour at 48ºC, where the temperature with the most similar position gel band to the control is the approximate reversible temperature. Additional controls at lower-optimal (48 ºC) and higher (55ºC) temperatures for the entire 4.5 hour duration of the experiment were also included, to show how the rate of assembly is lessened as equilibrium conditions (i.e. close to the reversible temperature) are approached.

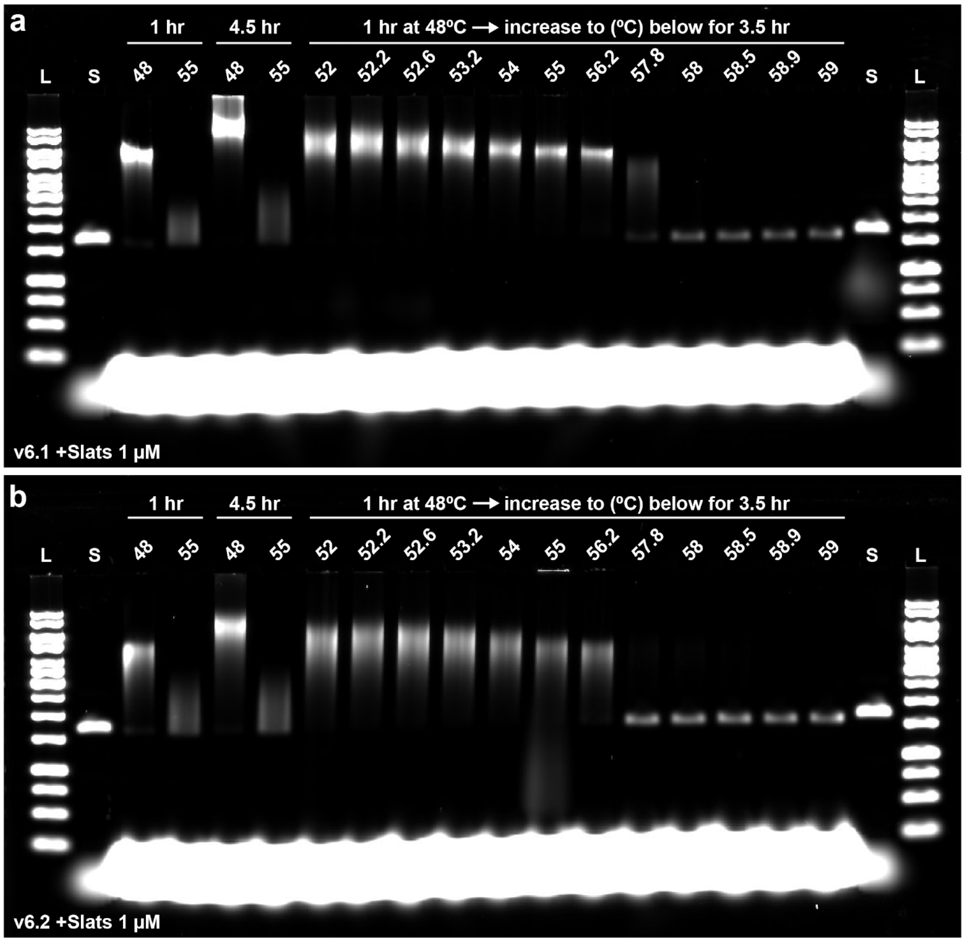


**Supplementary Figure S25**: **a–b**, Agarose gels showing the approximate reversible temperature of slat binding for v6.1 and v6.2 respectively. All assemblies contained 16 mM $MgCl_2$ and 1 µM each slat. The 56.2ºC lane for both versions most closely resemble the gel position of the 1 hour 48ºC control, indicating that assembly from the initial 1 hour incubation was negligible. This suggests the reversible temperature is close to 56.2ºC for v6.1 and v6.2 at these reaction conditions. Furthermore, growth of ribbons was much more minimal when the entire 4.5 hour assembly was carried out near the reversible temperature (55ºC) versus much lower than the reversible temperature (48ºC). We highlight that the far-from-equilibrium 48ºC temperature did not show evidence of spurious ribbon formation for either v6.1 or v6.2 (see **Figure 2c** and **Supplementary Figure S20**).

#### S5—Quantification of spontaneous nucleation under suboptimal low-temperature growth conditions

**Overview:** In **S5**, we show the relationship between spontaneous nucleation and reaction parameters. We focus specifically on v6.1 and v6.2. First, **S5.1** shows that spontaneous nucleation of ribbons becomes more prevalent at temperatures below the optimal seeded growth temperatures. Second, **S5.2** and **S5.3** quantifies how changes to reaction parameters (i.e. increasing temperature, lowering the concentration of slats, lessening $MgCl_2$, using a different sequence variant) lessens spurious ribbon formation. We determine that marginally increasing the nucleation temperature (i.e. from 41 to 44.7ºC), decreasing the concentration of the slats (i.e. from 1 to 0.2 µM each), or decreasing $MgCl_2$ (i.e. from 16 to 10 mM) could each lessen relative spurious assembly between 100–1000 fold (see **S5.2**). Moreover, we find that v6.2 is particularly resistant to spontaneous nucleation, with 10–100 fold fewer spontaneous ribbons versus v6.1 for the temperatures tested (see **S5.3**). We hypothesize that these changes to parameters can be combined to multiply the relative decreases in spontaneous nucleation. However, this was not tested because the small numbers of ribbons in question would be below the SYBR-Gold detection limit.

It is noted that all quantitative measures of spontaneous nucleation were measured at suboptimal growth temperatures. Spuriously formed ribbons, to the extent they may exist, could not be measured at optimal temperatures because the small number was below SYBR-Gold detection limit. Again, we hypothesize that the changes to parameters as noted in this section could be applied to lessen spontaneous nucleation at optimal growth temperatures, although it was not tested here.

##### S5.1—Increased spurious nucleation observed at low temperatures

In **Supplementary Figure S26a**, incubation of v6.1 at suboptimal growth temperatures (ranging 34–44ºC for 16 hours, 1.0 µM each slat, 16 mM $MgCl_2$) showed increased spuriously formed ribbons, compared to what was noted at 46ºC in **Supplementary Figure S24**. However, the molecular mass of ribbons became lower as the temperature was lessened, suggesting that assembly was slower at lower temperature. In **Supplementary Figure S26b**, assembly was performed using two temperature steps: first, the reaction was incubated at a lower "nucleation" temperature (i.e. below 46ºC for ~6 hours); second, the reaction was heated to a higher optimal "growth" temperature (i.e. 50ºC for ~10 hours). We found that two-step incubation caused spontaneous nuclei to grow into ribbons of similar length, thus normalizing spontaneous nucleation between various suboptimal temperatures. Moreover, **Supplementary Figure**

**S26c** shows that the number of spontaneous nuclei was relatively consistent and within a single order of magnitude for the v6.1 slats at suboptimal temperatures ranging 37–42.1ºC. Additionally, two-step incubation in **Supplementary Figure S26b** revealed limited spurious assembly at 4, 25, and 44.1ºC, though the absolute number of ribbons were far fewer compared to the aforementioned temperature range. The observations in **S5.1** taken together suggest the following about how the slats behave: (1) Spurious nucleation is slower as the optimal growth temperature (i.e. ≥ 46ºC) is approached; (2) Formation of spurious nuclei is relatively constant for some range of temperatures (e.g. 37–42.1ºC) below the growth optimum. However, assembly of these nuclei into ribbons is progressively slower as temperature is decreased; and (3), The slats at temperatures much lower (i.e. 4ºC and 25ºC) than the growth optimum form relatively few spurious assemblies.

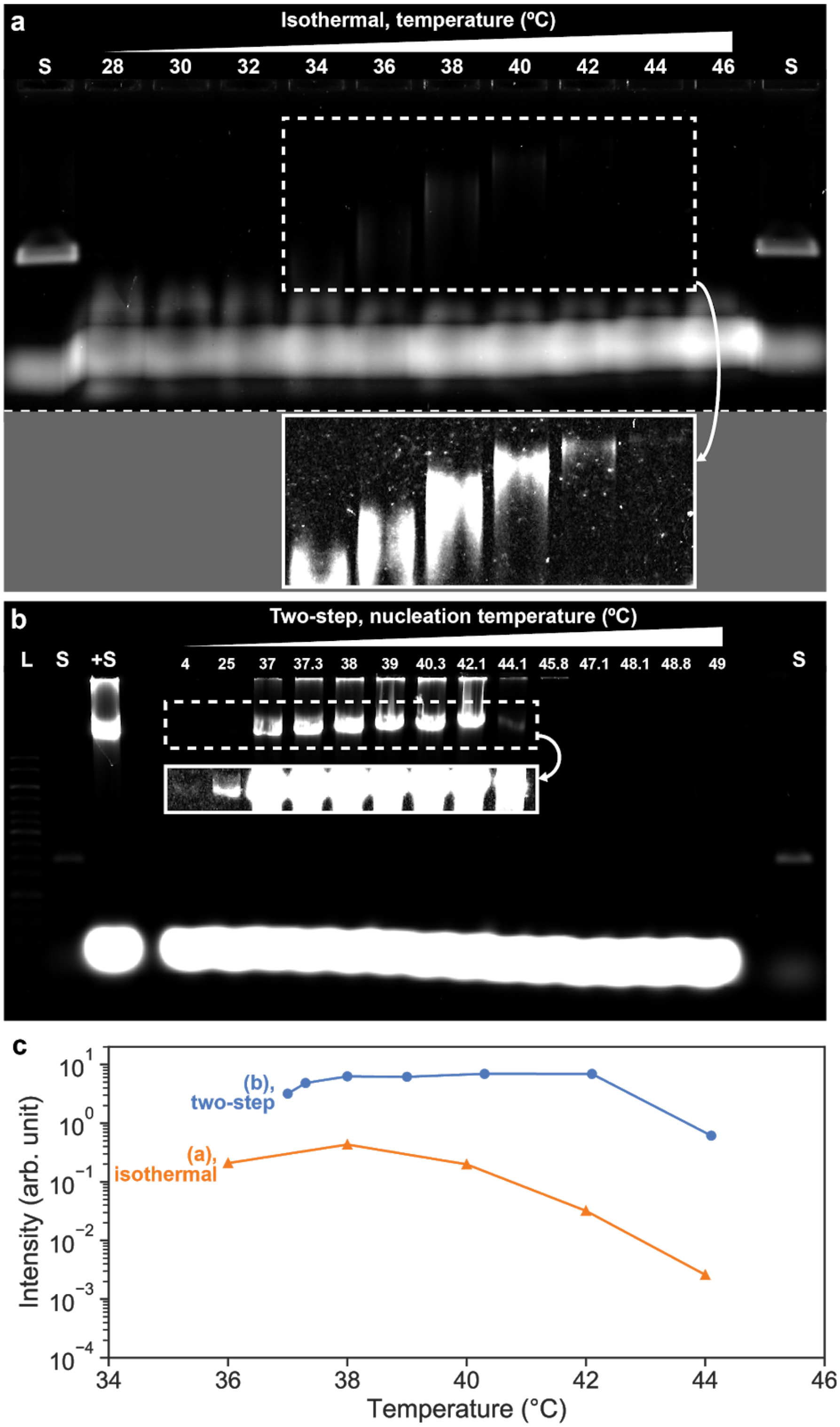


**Supplementary Figure S26:** Seedless assembly of sequence variant 6.1 at suboptimal growth temperatures (16 mM $MgCl_2$, 1 µM each slat). **a**, Agarose gel showing no-seed reactions with isothermal incubation for ~16 hours at the temperatures shown above each well. Spuriously nucleated ribbons of various lengths can be observed for 34 to 44ºC, seen more clearly in the contrast and brightness adjusted inset. **b**, Agarose gel showing no-seed reactions incubated at two different temperatures, with the contrast and brightness adjusted inset showing the fainter bands more clearly. The relative length of the spuriously assembled ribbons is similar at all temperature points with bands at similar positions in the gel. The two-step temperature incubation can therefore be used to visualize and normalize lengths of nuclei formed at lower temperatures. **c**, Plot of gel densitometry measurements, showing more assembly in *b* with two-step growth versus isothermal growth in *a*.

#### S5.2—Reducing spurious nucleation by increasing temperature, decreasing concentration of slats, or decreasing $MgCl_2$ concentration

In **S5.2**, we quantify how varying experimental parameters including temperature, concentration of slats, and $MgCl_2$ changes the number of spuriously nucleated ribbons for v6.1. We apply the variations of the two-step isothermal growth protocol noted in **S5.1** to normalize lengths of the spurious ribbons for quantification. In **Supplementary Figure S27**, reactions are first incubated at a nucleation-prone temperature (40ºC for 4–6 hours, unless otherwise noted) and then normalized at an optimal growth temperature (50ºC for 10–16 hours). Unless otherwise noted, these trials contained 1.0 µM each slat and 16 mM $MgCl_2$. All experiments also included seed-initiated controls (+S), where growth was initiated from 2 nM seed, with growth conducted at 50ºC only for the latter stage of the two-step incubation.

In **Supplementary Figure S27 a–b**, we nucleated the slats at temperatures from 41–45ºC to see how spontaneous nucleation is lessened as optimal growth temperatures are approached. We found that increasing the nucleation temperature from 41ºC to 44.7ºC lessened spontaneous nucleation 1000-fold.

In **Supplementary Figure S27 c–d**, we nucleated the slats at 40ºC with slat concentrations ranging from 0.2–2 µM. Before proceeding to the growth normalization step, one-volume of reaction was transferred to one-volume of preheated denatured 50ºC reaction buffer containing additional slats or buffer to equalize concentration of the slats to 1 µM each for growth normalization. This particular experiment necessitated preparation of different buffers with small variations in the concentration of slats. To eliminate possible manual pipetting errors that could have impacted the final concentration of slats, the reaction mixtures were prepared using an Echo 525 acoustic liquid handler. We found that decreasing the concentration of slats from 1 µM to 0.2 µM each lessened spontaneous nucleation nearly 1000-fold.

In **Supplementary Figure S27 e–f**, we nucleated the slats at 40ºC in 10–16 $MgCl_2$. Prior to the growth normalization step, three-volumes of reaction was transferred to one-volume of preheated 50ºC reaction buffer containing additional $MgCl_2$ to equalize $MgCl_2$ to 16 mM for final growth. Slats in the final growth become diluted to 0.75 µM because of this dilution step. We found that decreasing the concentration of $MgCl_2$ from 16 mM to 10 mM lessened spontaneous nucleation nearly 1000-fold.

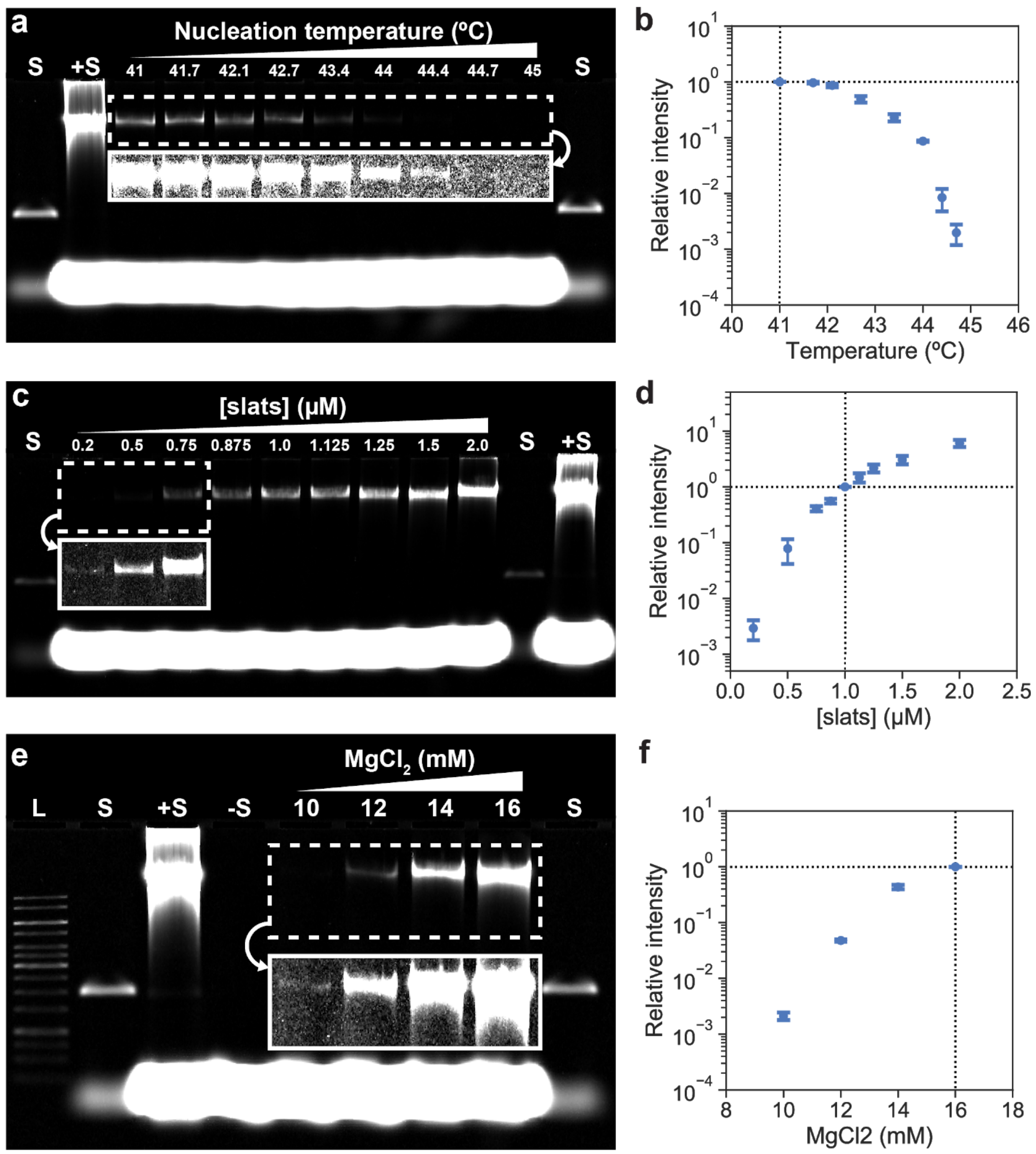


**Supplementary Figure S27:** Characterization of spurious nucleation by agarose gel of variant 6.1 at various temperatures, concentrations of slats, and $MgCl_2$. Gel lanes (+S) are control reactions where growth was initiated with the seed, (S) are seed normalization controls, and (L) is the ladder. **a**, Single gel of triplicate experiment where slats were nucleated at temperatures from 41–45ºC. Nucleation at 45ºC was below the sensitivity limit of gel detection. **b**, Plot of mean relative band intensity with respect to the

41ºC condition. There is nearly a 1000-fold decrease in relative intensity (i.e. number of nuclei) at 44.7ºC vs 41ºC. **c**, Single gel of duplicate experiment where initial concentration of slats during nucleation was varied from 0.2–2 µM each. **d**, Plot of mean relative band intensity with respect to the 1 µM each condition. There was nearly a 1000-fold decrease in the number of nuclei formed at 0.2 µM vs 1 µM each slat. **e,** Single gel of triplicate experiment where slats were nucleated in 10–16 mM $MgCl_2$. Gel lane (-S) is a no-seed control where three-parts of freshly denatured slats in 16 mM $MgCl_2$ was pipetted into one-part preheated 50ºC normalization buffer to show that momentary temperature decrease during reaction transfer did not lead to measurable spurious assembly. **f,** Plot of mean relative band intensity with respect to the 16 mM $MgCl_2$ condition. There was more than a 10-fold decrease in the number of nuclei formed at 12 mM vs 16 mM $MgCl_2$, and nearly 1000-fold decrease for 10 mM vs 16 mM $MgCl_2$. Inset sections of gels in white are contrast and brightness adjusted to better-show samples with lower concentrations of ribbons. All error bars are ±SD.

##### S5.3—Reducing spurious nucleation by changing the sequence design from v6.1 to v6.2

In **S5.3**, we quantify the relative amount of spurious ribbons for v6.2 versus v6.1 in an attempt to address how sequence design influences spontaneous nucleation. This section expands the results from **S4.5** with more readily quantified gel densitometry from lower nucleation temperatures. Again, we apply the variations of the two-step isothermal growth protocol noted in **S5.1** to normalize lengths of the spurious ribbons for quantification. In **Supplementary Figure S28**, reactions are first incubated at a nucleation-prone temperature (either 4, 25, 30, 37, or 40–44.3ºC for 6 hours) and then normalized at an optimal growth temperature (50ºC for 17 hours). These trials contained 1.0 µM each slat and 16 mM $MgCl_2$. All experiments also included seed-initiated controls (+S), where growth was initiated from 2 nM seed, with growth conducted at 50ºC only for the latter stage of the two-step incubation. We determine that v6.2 has 10–100 fold less spontaneous nucleation compared to v6.1, depending on the temperature tested.

It is noted that making a quantitative comparison between v6.1 and v6.2 requires considering that the average lengths of ribbons assembled were shorter for v6.2 (see **S6.1.1** and **S6.1.3**). To account for the length differences between versions, all gel intensity values for v6.2 were scaled by the ratio of gel intensities between the (+S) controls of each version.

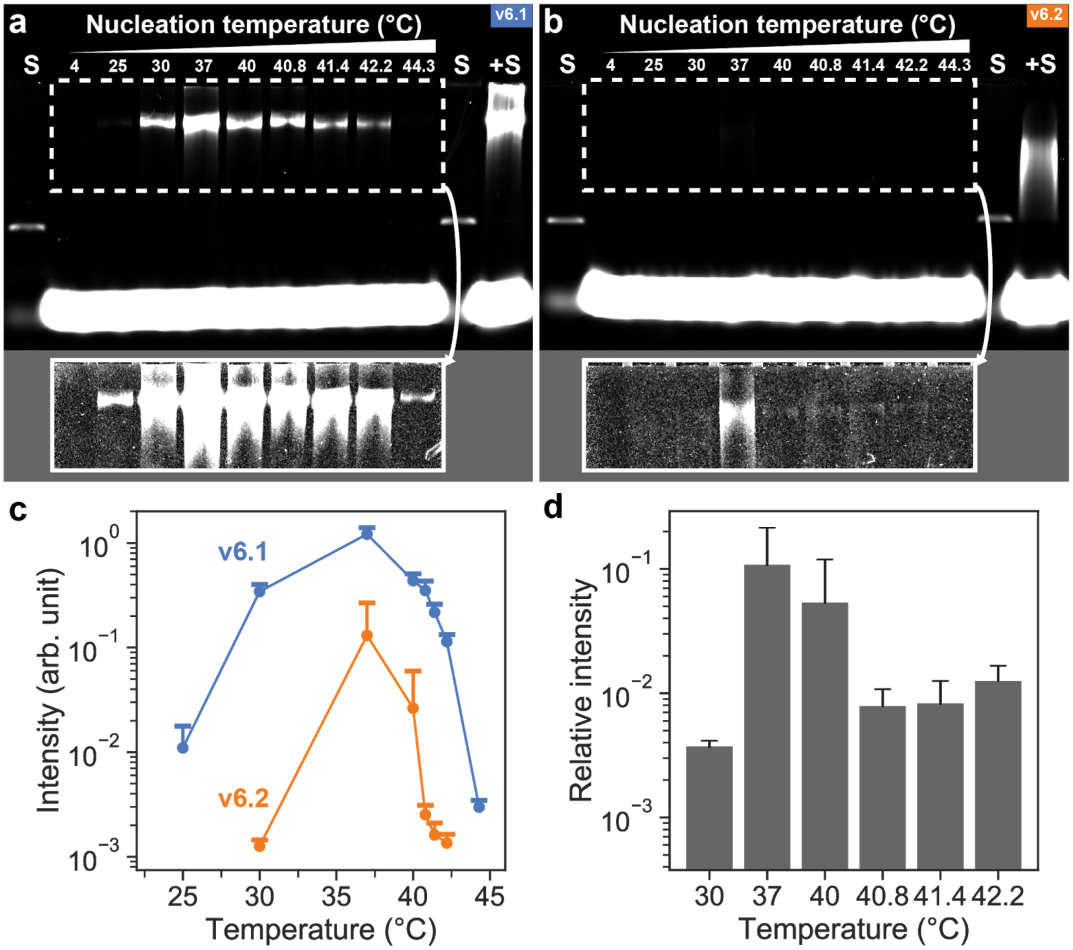


**Supplementary Figure S28**: Characterization of spurious nucleation for v6.1 and v6.2 at select temperature points. **a**, Single gel of triplicate experiment of nucleation of v6.1 slats. **b**, Single gel of triplicate experiment of nucleation of v6.2 slats. Inset sections of the gel in white are contrast and brightness adjusted to better-show samples with lower concentrations of ribbons. **c**, Plot of mean gel intensity measurements from *a–b*, where less spontaneous nucleation is seen for v6.2. **d**, Plot of mean relative gel intensity for v6.2 with respect to v6.1 at nucleation temperatures where a band could be measured for each version. There are ~10–100 fold fewer spontaneous nuclei for v6.2 versus v6.1, depending on the nucleation temperature. Gel intensity values for v6.2 were corrected as noted in the **S5.3** text. All error bars are the +SD.

#### S6—Ribbon assembly characteristics by experimental observation and modelling

**Overview:** To validate the mechanism by which slats assemble into crisscross ribbons, we built a stochastic model to simulate their assembly, and developed an analytical solution of this model. We tested the model by fitting to length distribution data obtained by TEM and agarose gel densitometry.

We postulate that there are three distinct phases of assembly: initiation, growth, and termination. Initiation can occur from either a DNA seed, or from a spuriously formed nucleus. Growth subsequently proceeds at a relatively constant rate until ribbon growth terminates due to accumulation of defects or impurities. The growth may occasionally stall and subsequently recover, for example due to an incorrect slat temporarily binding to the terminus of a ribbon.

From these mechanistic assumptions, we reproduce experimentally measured ribbon length distributions over time in a stochastic model considering only nucleation rate, growth rate, stalling probability, and termination probability. Each available seed has a fixed probability of nucleating a ribbon at each timepoint. Spurious nucleation occurs at a fixed rate throughout the assembly time. Once initiated, a ribbon grows at a fixed rate until its growth is terminated with a fixed probability of termination at each timepoint. At each timepoint in ribbon growth, the ribbon has a fixed probability of stalling (stopping growth), or if stalled, of restarting growth.

We assume that effects from changes in concentration of the slats from monomer depletion during growth are negligible because most experiments had large excesses of slats (1–1.5 μM for each slat). Additionally, this model simplifies growth as a continuous coarse-grained increase of ribbon length, as opposed to discrete incorporation of single slats.

| **Parameter** | **Notation** | **Fitting** |
|---|---|---|
| Growth rate | $l_{growth}$ | TEM data |
| Probability of termination | $p_{term}$ | TEM data |
| Probability of stalling and recovery | $p_{stall}$ | TEM data |
| Probability of seed nucleation | $p_{seed}$ | Gel data |

| | | |
|---|---|---|
| Spurious nucleation rate | $r_{nuc}$ | Gel data |
| Number of seeds | $N_{seed}$ | Constant/gel data |
| Total assembly duration | $T_{final}$ | Constant |
| Timepoint | $t$ | – |
| Length of ribbon $k$ | $l_k$ | – |
| State of ribbon $k$ (seed/growing/terminated) | $s_k$ | – |

**Supplementary Table S3:** List of parameters used in stochastic model and analytical solution, and the data used to fit each parameter (if applicable). Probabilities of termination, stalling and seed nucleation were used as rates in the analytical solution, and hence referred to as $\lambda_{term}$, $\lambda_{stall}$ and $\lambda_{seed}$ respectively in **S6.4**. The number of seeds is taken as constant for the stochastic simulations but is fitted in the gel data analysis.

**Stochastic model algorithm:** Begin with a population of $N_{seed}$ seeds (i.e. uninitiated ribbons) by initialising the ribbon state $s_k \leftarrow$ "*growing*" and length $l_k \leftarrow 0$ for ribbons $k \in \{1, \ldots, N_{seed}\}$.

For every timepoint $t \in \{1, \ldots, T_{final}\}$:

1. Generate $r_{nuc}$ new ribbons by initialising each of these ribbons with state $s_k \leftarrow$ "*growing*" and length $l_k \leftarrow 0$ for ribbons $k \in \{1, \ldots, r_{nuc}\}$.
2. For every stalled ribbon $k$ such that $s_k ==$ "*stalled*", set $s_k \leftarrow$ "*growing*" with a probability of $p_{stall}$.
3. For every growing ribbon $k$ such that $s_k ==$ "*growing*", add a constant length $l_{growth}$ to $l_k$, i.e. $l_k \leftarrow l_k + l_{growth}$.
4. For every ribbon $k$ such that $s_k ==$ "*growing*" or $s_k ==$ "*stalled*", set $s_k \leftarrow$ "*terminated*" with a probability of $p_{term}$.

5. For every growing ribbon $k$ such that $s_k$ == "*growing*", set $s_k \leftarrow$ "*stalled*" with a probability of $p_{stall}$.

The output of the algorithm is the sample of lengths $l_k$ for all the growing and terminated ribbons such that $s_k$ == "*growing*", $s_k$ == "*stalled*", or $s_k$ == "*terminated*".

When considering only seeded growth (for example in the TEM analyses in **Supplementary Figures S29−S32**), step 1 was omitted, effectively meaning $r_{nuc} = 0$. Similarly, when considering spurious assembly only, $N_{seed}$ was set to zero (i.e. run the algorithm without an initial population of ribbons).

#### S6.1—Kinetics by TEM observation versus model fit

The lengths of ribbons nucleated from a seed were measured using TEM for different assembly conditions and assembly times. For each timepoint we measured approximately 150 ribbons (actual N=130–168) , and so the number of seeds used for the simulations was also 150. Only ribbons that had a visible seed attached to one end were measured. Therefore, in the stochastic simulations we can ignore the contributions of spurious nucleation. We also assume that stalling and recovery occur by similar mechanisms (for example the reversible binding of incorrect slats), and therefore we take the probability of stalling and of recovery to be equal for simplicity. To justify this assumption, we tried fitting these two parameters separately, which resulted in very similar parameter values for stalling and recovery – having them as two separate variables also did not significantly improve the fit.

The $p_{stall}$, $p_{term}$, and $l_{growth}$ parameters in the stochastic model were fit to the TEM data using the Kolmogorov-Smirnov (KS) statistic[13] to compare the cumulative distribution functions of ribbon lengths of the model and the data. For every experimental condition, the mean of the KS statistics for each assembly time was minimised with the Nelder-Mead method[14]. Using a timestep of one minute, this resulted in a stalling and recovery probability of ~0.03-0.1 per timestep, a termination probability of ~0.00005-0.005 per timestep, and a growth rate of ~4-30 nm/minute (**Supplementary Figures S29−S32**). Fitted parameters for each simulation are shown in the respective figure legends.

*S6.1.1—v6.1 at optimal growth conditions*

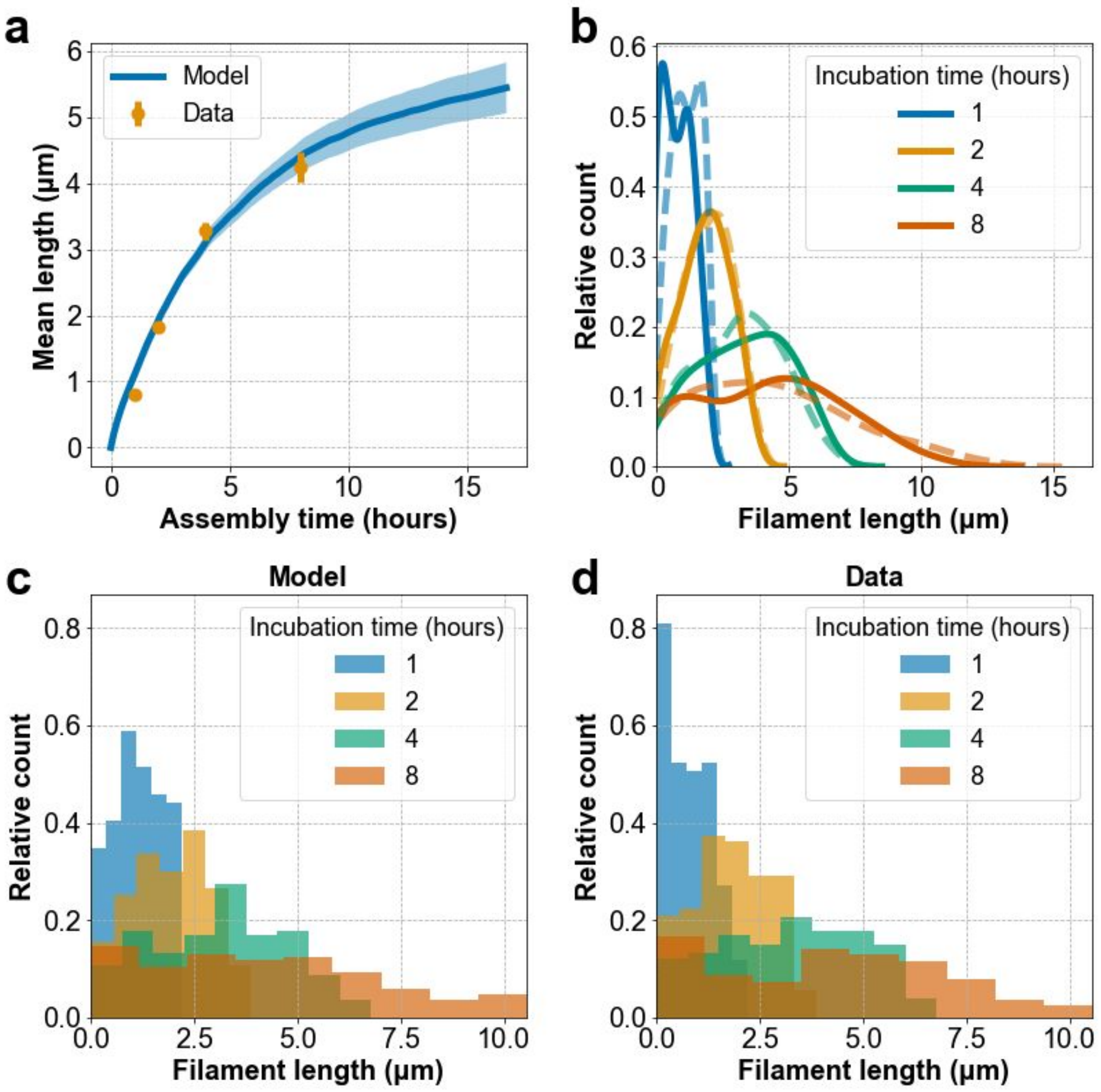


**Supplementary Figure S29:** Stochastic model fitting for sequence variant 6.1 at experimental conditions of 50°C, 16 mM $MgCl_2$, and 1 μM each slat. Estimated parameters are $p_{stall} = 0.037$ per minute, $p_{term} = 0.0030$ per minute, and $l_{growth} = 31.37$ nm/minute. **a**, Mean lengths of the distributions of model vs data. Error bars represent the standard error of the mean and are hidden by the markers for some of the points. **b**, Gaussian kernel density estimates for model (dashed line) vs data (solid line). **c–d**, Histograms of the unprocessed length distributions for the model and data respectively.

*S6.1.2—v6.1 at sub-optimal growth conditions*

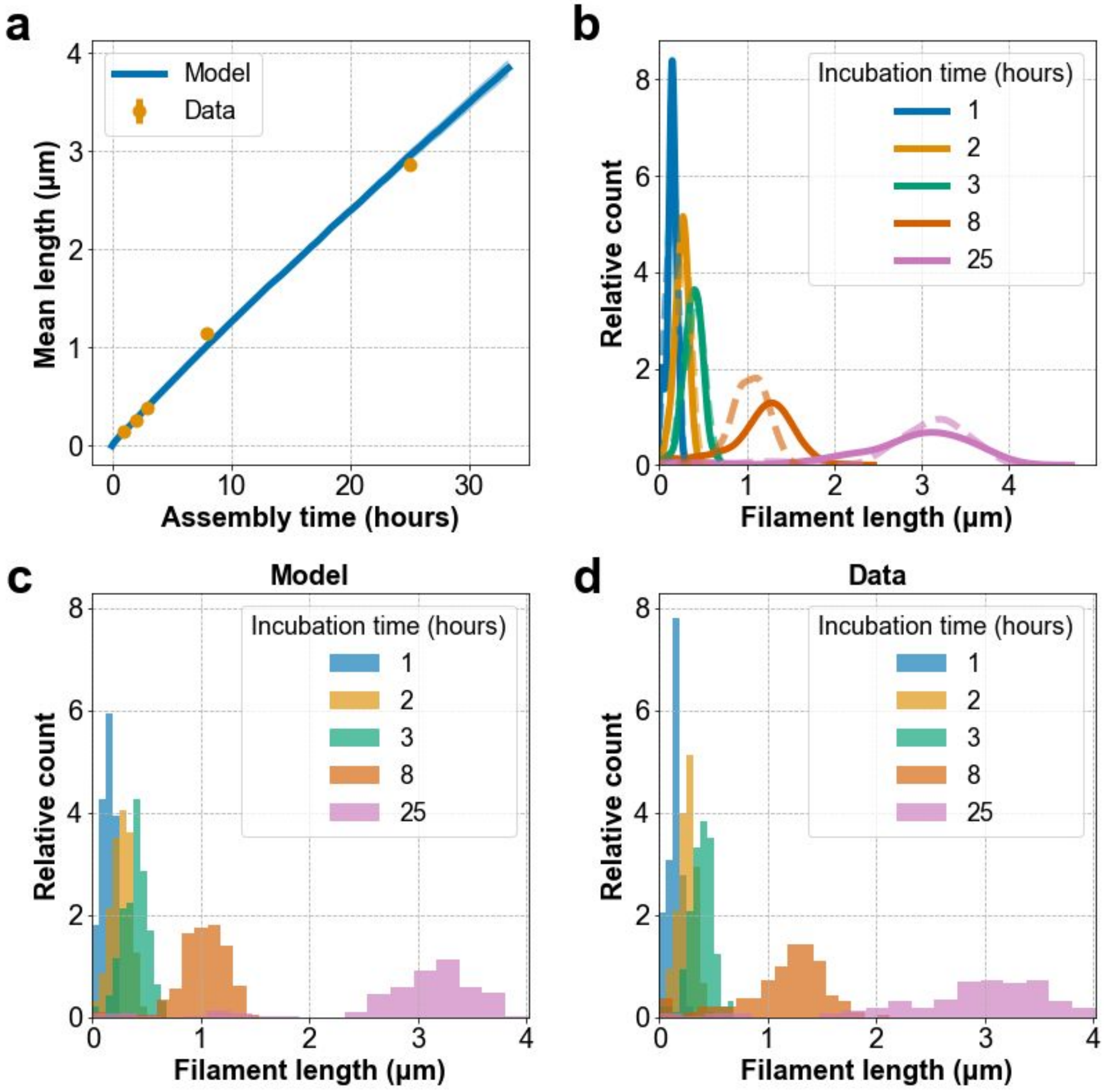


**Supplementary Figure S30:** Stochastic model fitting for sequence variant 6.1 at experimental conditions of 40°C, 20 mM $MgCl_2$, and 1 μM each slat. Estimated parameters are $p_{stall}$ = 0.063 per minute, $p_{term}$ = 0.000056 per minute, and $l_{growth}$ = 4.12 nm/minute. **a**, Mean lengths of the distributions of model vs data. Error bars represent the standard error of the mean and are hidden by the markers for some of the points. **b**, Gaussian kernel density estimates for model (dashed line) vs data (solid line). **c–d**, Histograms of the unprocessed length distributions for the model and data respectively.

*S6.1.3—v6.2 at optimal growth conditions*

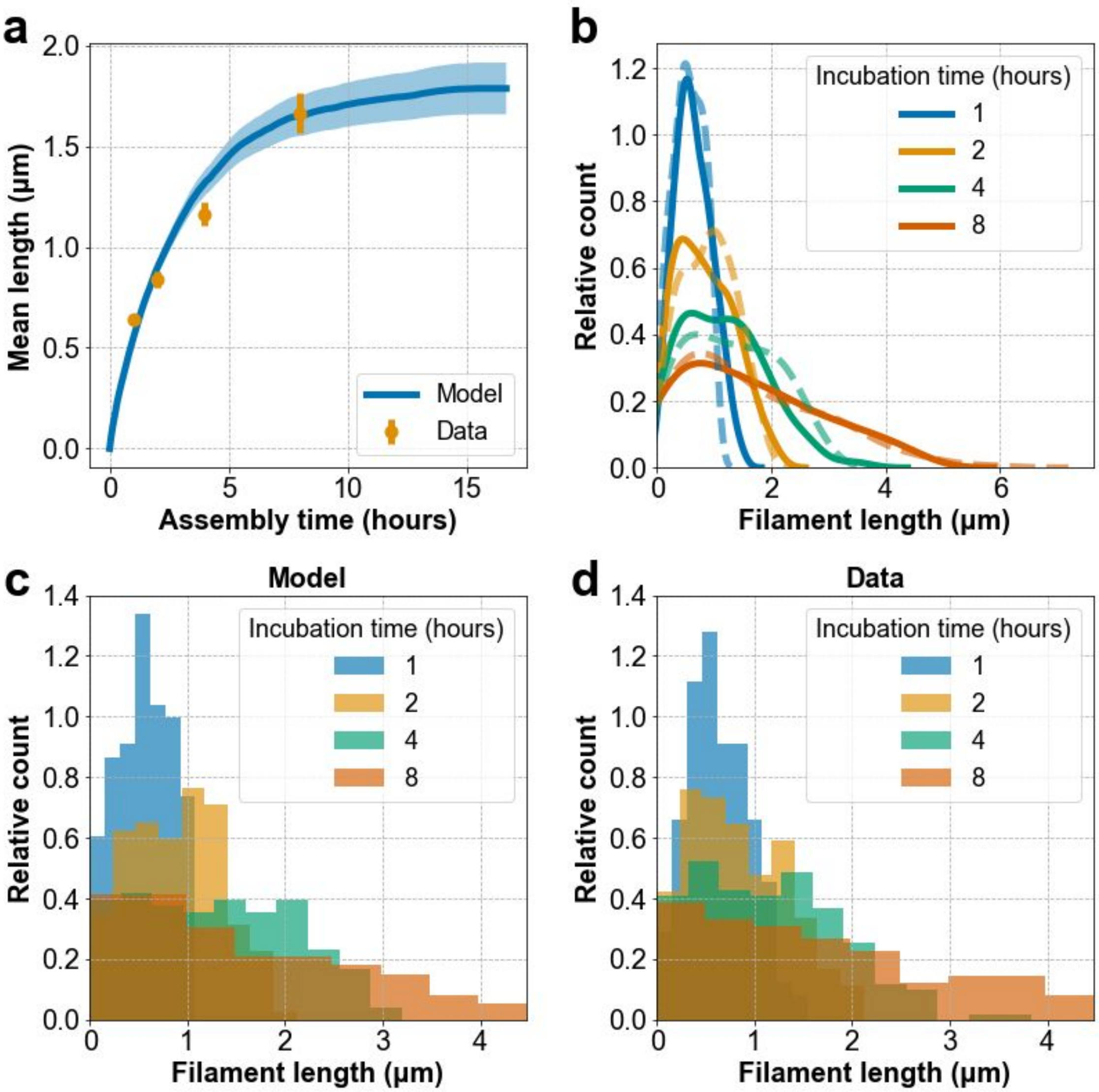


**Supplementary Figure S31:** Stochastic model fitting for sequence variant 6.2 at experimental conditions of 50°C, 16 mM $MgCl_2$, and 1 µM each slat. Estimated parameters are $p_{stall} = 0.047$ per minute, $p_{term} = 0.0050$ per minute, and $l_{growth} = 8.53$ nm/minute. **a**, Mean lengths of the distributions of model vs data. Error bars represent the standard error of the mean and are hidden by the markers for some of the points. **b**, Gaussian kernel density estimates for model (dashed line) vs data (solid line). **c–d**, Histograms of the unprocessed length distributions for the model and data respectively.

*S6.1.4—v6.2 at alternative optimal growth conditions with 1.5 µM each slat*

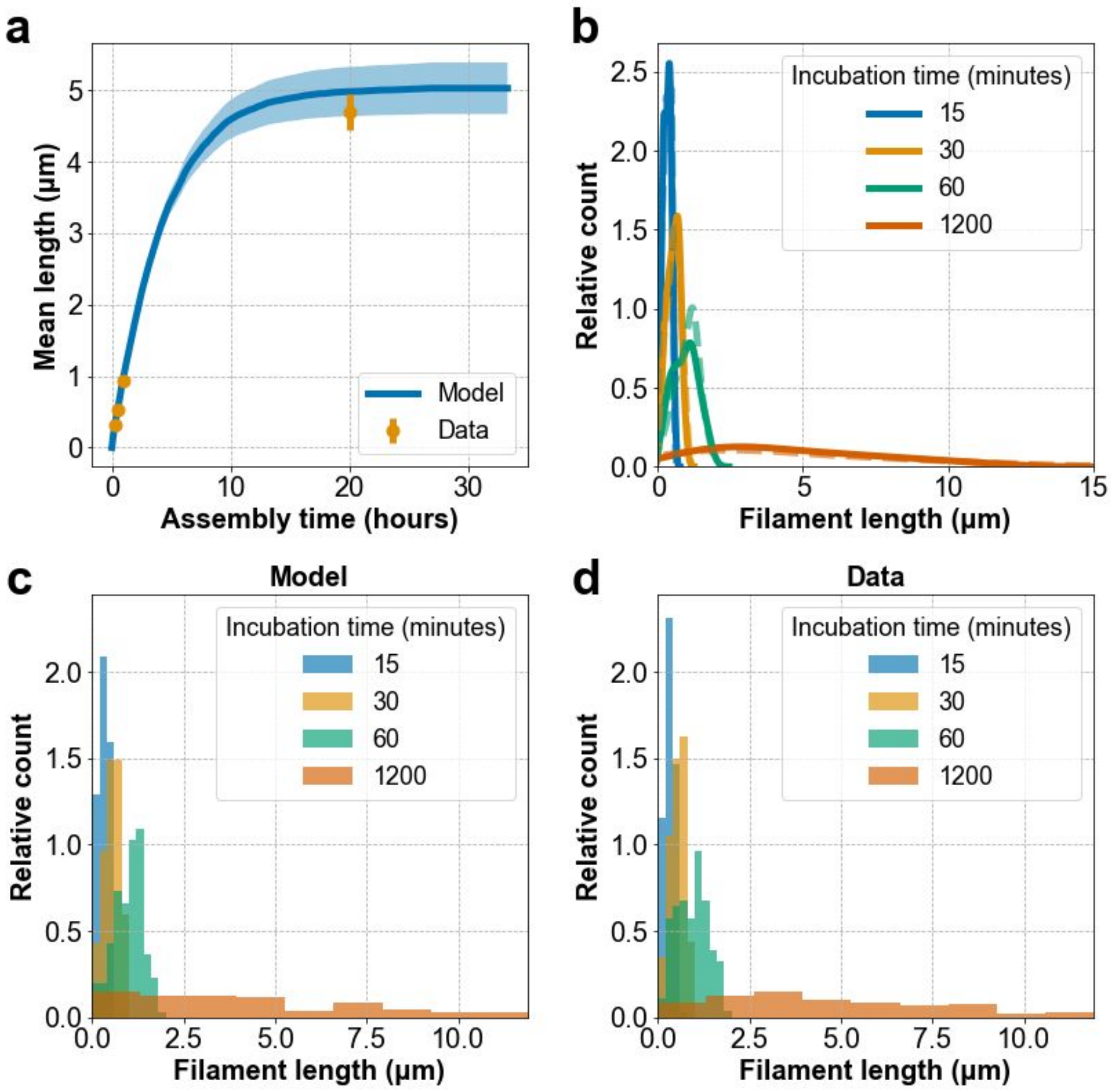


**Supplementary Figure S32:** Stochastic model fitting for sequence variant 6.2 at experimental conditions of 50°C, 14 mM $MgCl_2$, and 1.5 µM each slat. Estimated parameters are $p_{stall} = 0.116$ per minute, $p_{term} = 0.0031$ per minute, and $l_{growth} = 31.84$ nm/minute. **a**, Mean lengths of the distributions of model vs data. Error bars represent the standard error of the mean and are hidden by the markers for some of the points. **b**, Gaussian kernel density estimates for model (dashed line) vs data (solid line). **c–d**, Histograms of the unprocessed length distributions for the model and data respectively.

#### S6.2—Observation of spurious ribbons on agarose gels versus model fit

We were unclear whether the spurious assembly observed in experiments using certain reaction conditions was the result of nuclei that had formed during some early stage when the reagents were mixed, or if they were forming throughout the growth period at some constant spurious nucleation rate (i.e. $r_{nuc}$). To elucidate this process, we compared spurious and seed-initiated growth of sequence variant 6.1 on agarose gels versus time using reaction conditions that were either permissive to spontaneous growth (i.e. 40ºC, 20 mM $MgCl_2$, 1 μM each slat) or not (i.e. 52ºC, 20 mM $MgCl_2$, 1 μM each slat). We used the stochastic model and analytical solution (which were developed with the hypothesis that spurious nucleation occurs at constant rate $r_{nuc}$) to see if observed gel densitometry data for spurious and seeded-initiated assembly could be recreated *in silico*.

We assume measured intensity of a gel band at a given position on the gel is proportional to the number of ribbons where the relative SYBR-Gold gel stain fluorescence per ribbon scales linearly with its length (i.e. accounting for the amount of stain bound to the net amount of DNA in each ribbon). To apply the model to recreate the gel densitometry profiles, we scaled the ribbon counts by their corresponding length. The total intensity in both the experiment and model is the integration of the intensity vs gel position (i.e. ribbon length). For experimental data, gel intensity profiles were extracted using ImageJ and total intensity was calculated as the area under the curve.

We estimated $p_{stall}$, $p_{term}$, and $l_{growth}$ parameters from the TEM data (**Supplementary Figure S30**). These parameters were fixed in the analytical solution of the model (see **S6.3** for further motivation and **S6.4** for derivation), which was used to fit the $r_{nuc}$ parameter to the total intensities of assembly when no seed was present. The fitted $r_{nuc}$ parameter was subsequently used to fit the number of seeds $N_{seed}$ to the total intensities of assembly in seeded conditions based on the analytical solution.

To recreate the "seed" peak in the gel intensity profiles for seeded assembly, we needed to assign leftover seeds a certain size (relative to ribbon length). Thus, we took seed size to be 85.2 nm, and added Gaussian noise with a mean of 0 nm and standard deviation of 50 nm to prevent singularities in plotting. While it is not possible to directly quantify the effective size of the DNA seed relative to ribbon length in terms of how it would run on a gel, we estimated its effective size by considering that the seed contributes 4032 nucleotides. The average length of a single slat is 71 nucleotides, and we can approximate that each slat contributes around 1.5 nm to the length of the ribbons. Thus, the effective seed size can be estimated to be

85.2 nm, though in practice simulations are robust to variations in the seed size used since it simply determines the starting position of the simulated gel intensity profile.

Using gel data for experimental conditions permissive to spurious nucleation (40°C, 20 mM $MgCl_2$, sequence variant 6.1), we fit the analytical equation for total gel intensity to the mean areas of the gel intensity profiles (**Supplementary Figure S33a** for gel, **Supplementary Figure S33b** for fit). The ratio between the number of seeds present and the spurious nucleation rate (per minute) was estimated to be 1148. This would mean that for a concentration of 2 nM seed at these conditions, we would expect a spurious nucleation rate of 1.7 pM per minute. Note that we are unable to make similar estimates for other experimental conditions as spurious nucleation is too low to be detectable by gel analysis.

The stochastic model was executed with $N_{seed} = 22961$ and $r_{nuc} = 20$ (corresponding to the ratio calculated through the analytical solution fit) and the parameters estimated in **Supplementary Figure S30**. A continuous distribution of ribbon lengths was generated using Epanechnikov kernel density estimation based on the model outputs. Given that gels separate linear DNA logarithmically based on length, we assumed that logarithmic separation based on ribbon length would be a reasonable qualitative approximation of how ribbons may run on the gel. Thus, the products of the relative counts and lengths were plotted on a logarithmic scale to recreate the expected gel intensity profiles for different assembly durations, giving good qualitative agreement between the model and the data (**Supplementary Figure S33c**).

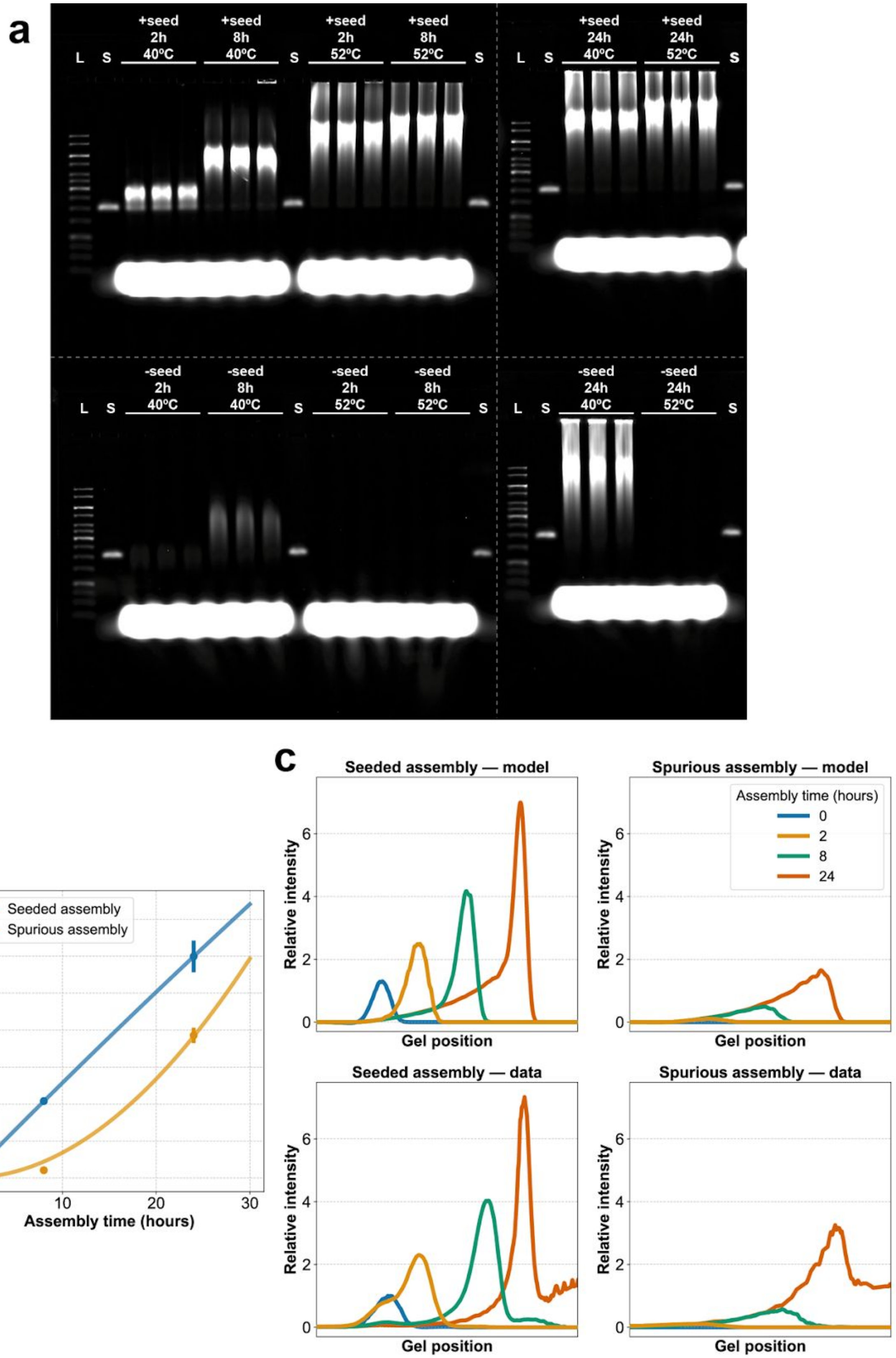
a
L
S
+seed 2h 40ºC
+seed 8h 40ºC
+seed 2h 52ºC
+seed 8h 52ºC
+seed 24h 40ºC
+seed 24h 52ºC
-seed 2h 40ºC
-seed 8h 40ºC
-seed 2h 52ºC
-seed 8h 52ºC
-seed 24h 40ºC
-seed 24h 52ºC
b
Seeded assembly
Spurious assembly
Total intensity relative to seed
Assembly time (hours)
0
10
20
30
40
50
60
70
c
Seeded assembly — model
Spurious assembly — model
Seeded assembly — data
Spurious assembly — data
Assembly time (hours)
0
2
8
24
Relative intensity
Gel position

**Supplementary Figure S33:** Reproduction of gel intensity profiles using the analytical solution. **a,** Gel data for sequence variant 6.1 at experimental conditions of 40 or 52°C, 20 mM $MgCl_2$, and 1 µM each slat, displaying spurious and seeded growth at 40°C and no observable spurious nucleation at 52°C. **b**, Analytical solution to the stochastic model (solid line) compared to the 40°C data from the gel in *a* using parameters estimated in the TEM data fit of **Supplementary Figure S30**. Error bars represent the standard error of the mean and are hidden by the markers for some of the points. Note that here seeded assembly is the sum of assembly from seed nucleation as well as spurious nucleation. **c**, Averaged gel intensity profiles for 40°C data from gel *a* (bottom row) and the stochastic model using the parameters estimated in *b* (top row).

#### S6.3—Analytical solution of ratiometric gel analysis

In order to be able to calculate the expected ratio of gel intensities (proportional to the total mass of ribbons present) resulting from spurious nucleation of a 100-hour incubation relative to a 1-hour incubation (the experiment in **S4.5**), we derived an analytical solution to our probabilistic model (see **S6.4** for derivation). This analytical solution was found to be in good agreement with the stochastic model simulations (**Supplementary Figure S34a**).

Under this model, the expected ratio solely depends on stalling and termination probability (**Supplementary Figures S34b−c**). Based on our estimates of these probabilities from fitting the TEM data, through the analytical solution we can expect an approximately 500- to 8000-fold increase in mass (corresponding to total gel intensity) due to spurious nucleation for a 100-hour incubation relative to a 1-hour incubation. When comparing the increase in mass after 100 hours to that after 1 hour and ignoring stalling, with zero termination we would expect a 10,000-fold increase in mass (quadratic growth as a result of a linear increase in the number of ribbons and linear growth), while with 100% termination at each step we would expect a 100-fold increase in mass (linear increase in number of ribbons).

The effect of stalling is minor in this case when modelling unseeded growth, though it is most pronounced for intermediate stalling probabilities. Intuitively, there would be a substantial proportion of ribbons that would not have stalled after 1 hour, while after 100 hours they would have likely reached a steady state of about 50% of non-terminated ribbons being stalled. At high stalling probabilities, the majority of ribbons at both 100 hours and 1 hour are likely to have reached the steady state of growing 50% of the time, and thus the ratio between 100 hours and 1 hour increases again. This is manifested as a small dip in **Supplementary Figure S34c**.

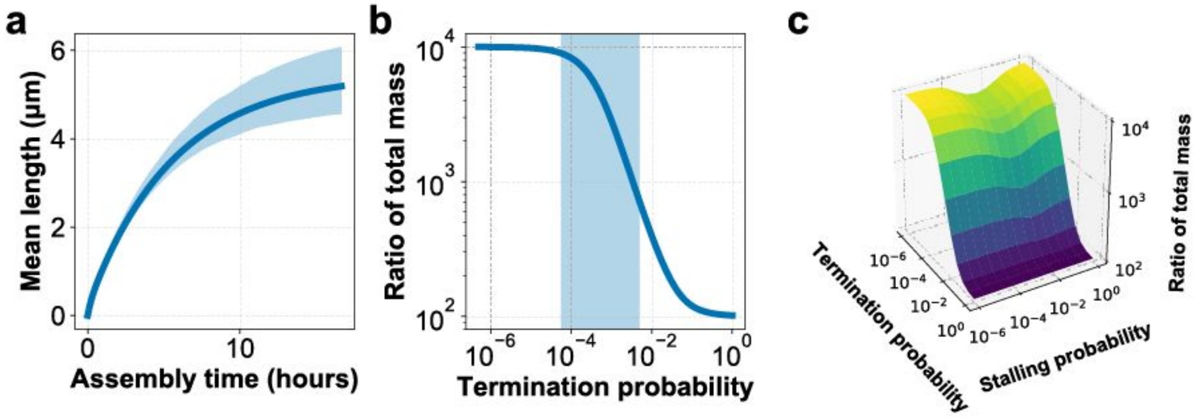


**Supplementary Figure S34:** Characterisation of analytical solution. **a**, Mean length versus time derived from the analytical solution (solid line) compared to the 95% confidence interval of 500 runs of the stochastic model (shaded area), using the parameters from the fit in **Supplementary Figure S29**. **b–c**, Expected ratio of the total mass of spuriously nucleated ribbons for a 100-hour incubation compared to a 1-hour incubation (equivalent to total integrated intensity on a gel) based on the analytical solution, assuming no stalling in *b* and with variable stalling in *c*. Shaded area in *b* corresponds to termination probabilities observed in **Supplementary Figures S29−S32**.

#### S6.4—Analytical solution derivation

In order to enable the ratiometric analysis discussed in **S6.3**, we derived an analytical solution to our stochastic model. Every growing ribbon is assumed to have a fixed probability of terminating at each timestep. Thus, for ribbons nucleated at $t = 0$, the proportion of ribbons terminated at each timestep *t* is given by the probability density function of an exponential distribution with rate parameter $\lambda_{term}$. If we ignore stalling, given a fixed growth rate $l_{growth}$, the length of a ribbon that has grown for $T$ timesteps is $l_{growth}\, T$. With a fixed probability of stalling and recovery from a stall $\lambda_{stall}$, the proportion of ribbons growing at timestep $t$ is given by $0.5 + 0.5\, e^{-2\,\lambda_{stall}\, t}$. Hence the total length of a ribbon growing for $T$ timesteps is the definite integral of this expression from $t = 0$ to $t = T$, which gives a total length of $\left(\frac{T}{2} - \frac{e^{-2\,\lambda_{stall}\, T} - 1}{4\,\lambda_{stall}}\right) l_{growth}$. Thus, for $N_{seed}$ growing ribbons, the total length of ribbons whose growth has terminated by $T_{final}$ is given by:

$$\sum_{\substack{k\in\{1\ldots N_{\text{seed}}\}\\ s_k=\text{"terminated"}}} l_k = \sum_{t=0}^{T_{\text{final}}} \left(\frac{t}{2} - \frac{e^{-2\,\lambda_{\text{stall}}\,t} - 1}{4\,\lambda_{\text{stall}}}\right) l_{\text{growth}}\, N_{\text{seed}}\, \lambda_{\text{term}}\, e^{-\lambda_{\text{term}}\,t} \tag{1}$$

And the total length of ribbons still growing at $T_{final}$ is given by:

$$\sum_{\substack{k\in\{1\ldots N_{\text{seed}}\}\\ s_k=\text{"growing"}}} l_k = \left(\frac{T_{\text{final}}}{2} - \frac{e^{-2\,\lambda_{\text{stall}}\,T_{\text{final}}} - 1}{4\,\lambda_{\text{stall}}}\right) l_{\text{growth}}\, N_{\text{seed}}\, e^{-\lambda_{\text{term}}\,T_{\text{final}}} \tag{2}$$

Equations (1) and (2) can be summed to give the total lengths for seeded assembly.

By comparison, for spurious assembly, the number of ribbons formed at each timestep corresponds to the fixed rate $r_{nuc}$. Thus, at the end of the incubation, there are $T_{final}r_{nuc}$ ribbons present. Therefore, equation (1) becomes:

$$\sum_{\substack{k\in\{1\ldots T_{\text{final}}r_{\text{nuc}}\}\\ s_k=\text{"terminated"}}} l_k = \sum_{t_{\text{nuc}}=0}^{T_{\text{final}}} \sum_{t=0}^{T_{\text{final}}-t_{\text{nuc}}} \left(\frac{t}{2} - \frac{e^{-2\,\lambda_{\text{stall}}\,t} - 1}{4\,\lambda_{\text{stall}}}\right) l_{\text{growth}}\, r_{\text{nuc}}\, \lambda_{\text{term}}\, e^{-\lambda_{\text{term}}\,t} \tag{3}$$

Similarly, equation (2) becomes:

$$\sum_{\substack{k\in\{1\ldots T_{\text{final}}r_{\text{nuc}}\}\\ s_k=\text{"growing"}}} l_k = \sum_{t_{\text{nuc}}=0}^{T_{\text{final}}} \left(\frac{(T_{\text{final}} - t_{\text{nuc}})}{2} - \frac{e^{-2\,\lambda_{\text{stall}}\,(T_{\text{final}}-t_{\text{nuc}})} - 1}{4\,\lambda_{\text{stall}}}\right) l_{\text{growth}} r_{\text{nuc}}\, e^{-\lambda_{\text{term}}\,(T_{\text{final}}-t_{\text{nuc}})} \tag{4}$$

Equations (3) and (4) can be summed to give the total lengths for spurious assembly. Correspondingly, seeded and spurious assembly can be summed to give total assembly in the presence of a seed (for example in the "Seeded assembly" conditions in **Supplementary Figure S33**).

#### S6.5—Consideration of delayed seed initiation

In the simulations described in previous sections, we assumed that all the seeds nucleate ribbons at the beginning of the incubation. However, for seeded assembly, we can still see a prominent band corresponding to uninitiated seed in certain experimental conditions (for example, the 2-hour gel data in

**Supplementary Figure S33**). Thus, we can conclude that the seeds do not all initiate ribbons at $t = 0$. While the approximation that seed initiation is instantaneous is sufficient to get good agreement with both TEM and gel data, in some instances we might be interested in describing the behaviour of the seeds themselves. In this case, we can assume that seed nucleation occurs at a constant rate, and therefore the fraction of ribbons nucleated at timestep $t$ follows an exponential distribution with rate $\lambda_{seed}$, i.e. there will be $N_{seed}\,\lambda_{seed}e^{-\lambda_{seed}\,t_{nuc}}$ ribbons that have nucleated at the timestep $t_{nuc}$. Thus, equation (1) becomes:

$$\sum_{\substack{k\in\{1\ldots N_{\mathrm{seed}}\}\\ s_k=\text{"terminated"}}} l_k = \sum_{t_{\mathrm{nuc}}=0}^{T_{\mathrm{final}}} \sum_{t=0}^{T_{\mathrm{final}}-t_{\mathrm{nuc}}} \left(\frac{t}{2} - \frac{e^{-2\,\lambda_{\mathrm{stall}}\,t} - 1}{4\,\lambda_{\mathrm{stall}}}\right) l_{\mathrm{growth}}\, N_{\mathrm{seed}}\lambda_{\mathrm{seed}}\, e^{-\lambda_{\mathrm{seed}}\,t_{\mathrm{nuc}}}\lambda_{\mathrm{term}}\, e^{-\lambda_{\mathrm{term}}\,t} \quad (5)$$

Similarly, equation (2) becomes:

$$\sum_{\substack{k\in\{1\ldots N_{\mathrm{seed}}\}\\ s_k=\text{"growing"}}} l_k = \sum_{t_{\mathrm{nuc}}=0}^{T_{\mathrm{final}}} \left(\frac{(T_{\mathrm{final}} - t_{\mathrm{nuc}})}{2} - \frac{e^{-2\,\lambda_{\mathrm{stall}}\,(T_{\mathrm{final}}-t_{\mathrm{nuc}})} - 1}{4\,\lambda_{\mathrm{stall}}}\right) l_{\mathrm{growth}}\, N_{\mathrm{seed}}\,\lambda_{\mathrm{seed}}e^{-\lambda_{\mathrm{seed}}\,t_{\mathrm{nuc}}}\, e^{-\lambda_{\mathrm{term}}\,(T_{\mathrm{final}}-t_{\mathrm{nuc}})} \quad (6)$$

In the stochastic simulation of the model, this can be implemented by beginning with a population of $N_{seed}$ seeds (i.e. uninitiated ribbons) by initialising the ribbon state $s_k \leftarrow$ "*seed*" and length $l_k \leftarrow 0$ for ribbons $k \in \{1, \ldots, N_{seed}\}$. Subsequently, at every timepoint $t \in \{1, \ldots, T_{final}\}$, for every uninitiated ribbon $k$ such that $s_k ==$ "*seed*", set $s_k \leftarrow$ "*growing*" with a probability of $p_{seed}$.

From the gel data in **Supplementary Figure S33**, we can estimate that $p_{seed}$ for these specific experimental conditions is around 0.003 per minute (**Supplementary Figure S35a**). Incorporation of seed initiation into the model is capable of reproducing a large portion of the variance in the data even when stalling is excluded from the model (**Supplementary Figures S35b−d**). However, since seed initiation is not capable of reproducing the true shape and symmetry of the length distributions, preference was given for stalling as the main source of variance in ribbon length, and for simplicity seed initiation was assumed to be instantaneous in all of the analyses in this paper.

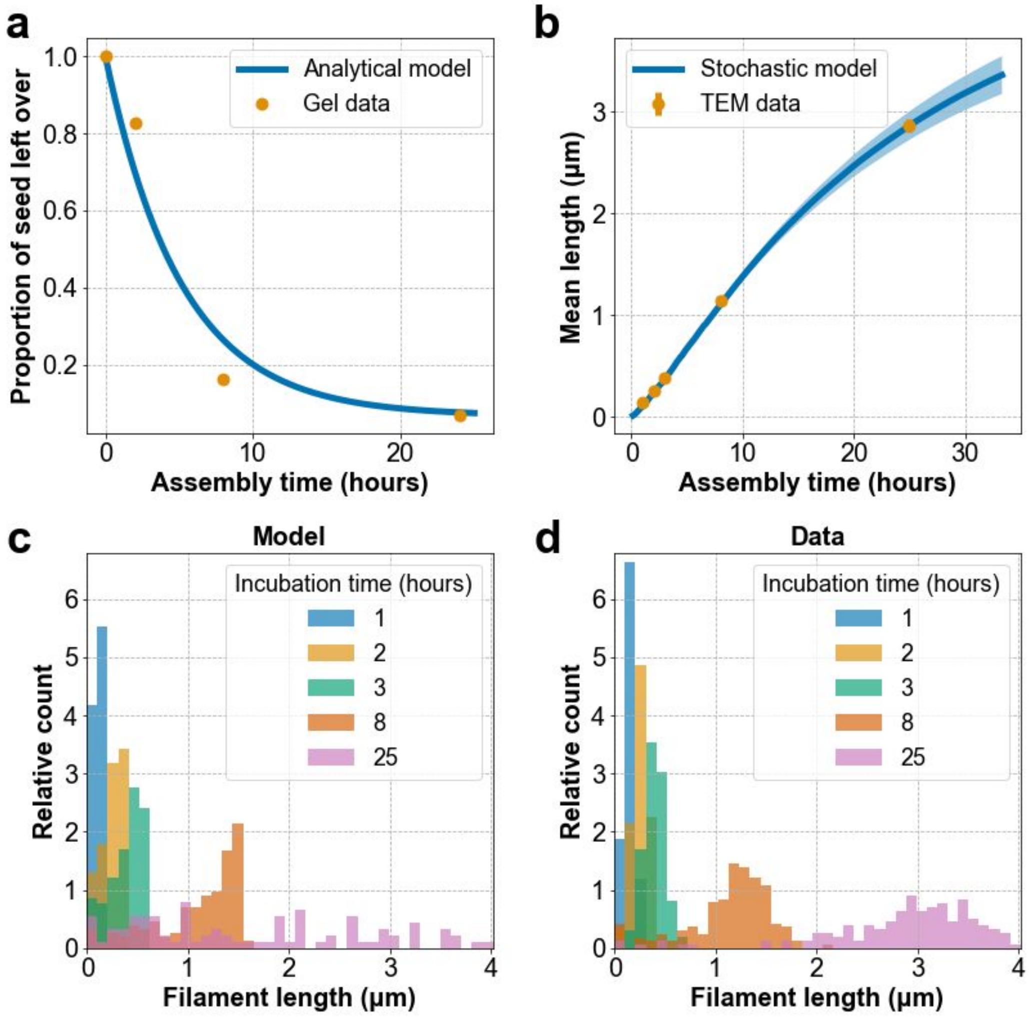


**Supplementary Figure S35:** Estimation of seed initiation delay and execution of model considering delayed seed initiation but not stalling. **a**, Single exponential decay corresponding to seed initiation at a rate of 0.0032 per minute compared to the seed peak height from averaged data in the gel from **Supplementary Figure S33**. **b−d**, Stochastic model fitting for sequence variant 6.1 at experimental conditions of 40°C, 20 mM $MgCl_2$, and 1 μM each slat (as in **Supplementary Figure S30**). Estimated parameters are $p_{seed}$ = 0.013 per minute, $p_{term}$ = 0.00067 per minute, and $l_{growth}$ = 3.22 nm/minute ( $p_{stall}$ taken as zero). **b**, Mean lengths of the distributions of model vs data. Error bars represent the standard error of the mean and are hidden by the markers for some of the points. **c**–**d**, Histograms of the unprocessed length distributions for the model and data respectively.

#### S7—Crisscross polymerization to sense nucleic acid sequences

**Overview:** In **S7** we generalize the seed architecture from a DNA origami structure (**Supplementary Figure S9**) to a 192 nt region comprised of six loops[15] of the M13 p8064 scaffold (**Supplementary Figure S36a**). We designed two different sets of nuc-y-slats (6 different length slat sequences) that respectively are able to use two specific regions of 192 nt length from the p8064 scaffold as a seed and subsequently trigger DNA slat ribbon growth with v6.3 x- and y-slats. We show that nuc-y-slats are able to trigger the ribbon polymerization with and without a specially designed cinch strand. Most importantly, we show that without the target p8064 ssDNA sequence we see no spurious ribbon assembly—i.e. no signal—on the agarose gel (**Supplementary Figure S36b**). In the future different sensors could be designed as pre-formed crisscross seeds that first are incubated with the analyte solution for target capture. Each sensor seed would be programmed to disintegrate during a subsequent timed destruction phase in the case that did not remain bound to the target for the full duration. Thus surviving seeds and subsequently polymerized ribbons would represent positive identifications.

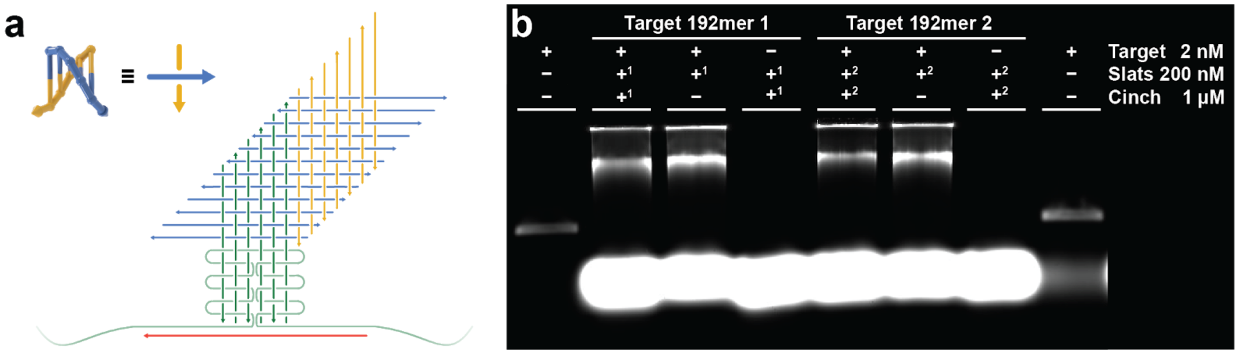


**Supplementary Figure S36**: Crisscross polymerization of DNA slats can be generalized to amplify ssDNA sequences into ribbons. **a**, Abstract strand diagram depicting how nuc-y-slats (dark green) bind to a 192 nt short region of the p8064 scaffold (light green) and subsequently trigger crisscross ribbon formation (gold y-slats and blue x-slats). The cinch strand is shown in red. **b**, Agarose gel showing crisscross polymerization seeded by two 192 nt long target regions of the M13 p8064 scaffold with and without a cinch strand. Superscripts on the + sign denotes that certain strands in the pool (i.e. nuc-y slats or the cinch strand) were customized to target a specific region of the target. $+^1$ denotes target region 1 of the p8064 scaffold, and $+^2$ denotes target region 2 of the p8064 scaffold.

#### S8—Twisted ribbons and tubes

**Overview:** In this section we expand the DNA slats flat ribbon architecture to form coiled ribbons and tubes. A DNA slat consists of multiples of four consecutive binding domains (i.e. $n = 6,\ 3 \times 4 = 12$ binding domains ***or*** $n = 8,\ 4 \times 4 = 16$ binding domains) that are each comprised of 5 or 6 bp long individual binding domains. Each four binding domain segment has either three 5 bp and one 6 bp binding domain (totalling 21 bp or 10.5 bp/turn), or two 5 bp and two 6 bp binding domain (totalling 22 bp or 11 bp/turn underwound DNA). The order of the 5 or 6 bp binding domain arrangement within the four binding domain long segments gives rise to ribbons with different coiled morphologies. We tested a total of six different binding domain arrangements shown in **S8.1**. The design shown in **Supplementary Figure S37a** is our standard flat ribbon architecture, whereas coiled ribbon designs are shown in **Supplementary Figure S37b–f**. In **S8.2** we show how the binding domain arrangements from the schematic in **Supplementary Figure S37a,b,f** result in ribbon morphologies that have very little coiling versus loosely or tightly coiled ribbons. We further show in **S8.3** which ribbons from **Supplementary figure S37b–f** yield the best tube morphologies once we add short sticky ends to the edge of the coiled ribbons in order to form closed continuous tubes.

In addition we note in **S8.4**, that besides the different ribbon morphologies that arise through the designs, particularly v8.3 exhibited the fastest rate of slat addition, with an estimated second-order rate constant of $10^6\ M^{-1}s^{-1}$ (**Supplementary Figure S40** and **Table S4**). In **S8.5** we test the assembly of v8.3 ribbons with and without seeds at low (0.1 μM each slat) and high (1 μM each slat) concentration of slats from 46–56ºC in 16 mM $Mg^{2+}$.

#### S8.1—Design notes on arrangement of binding sites to change ribbon morphology

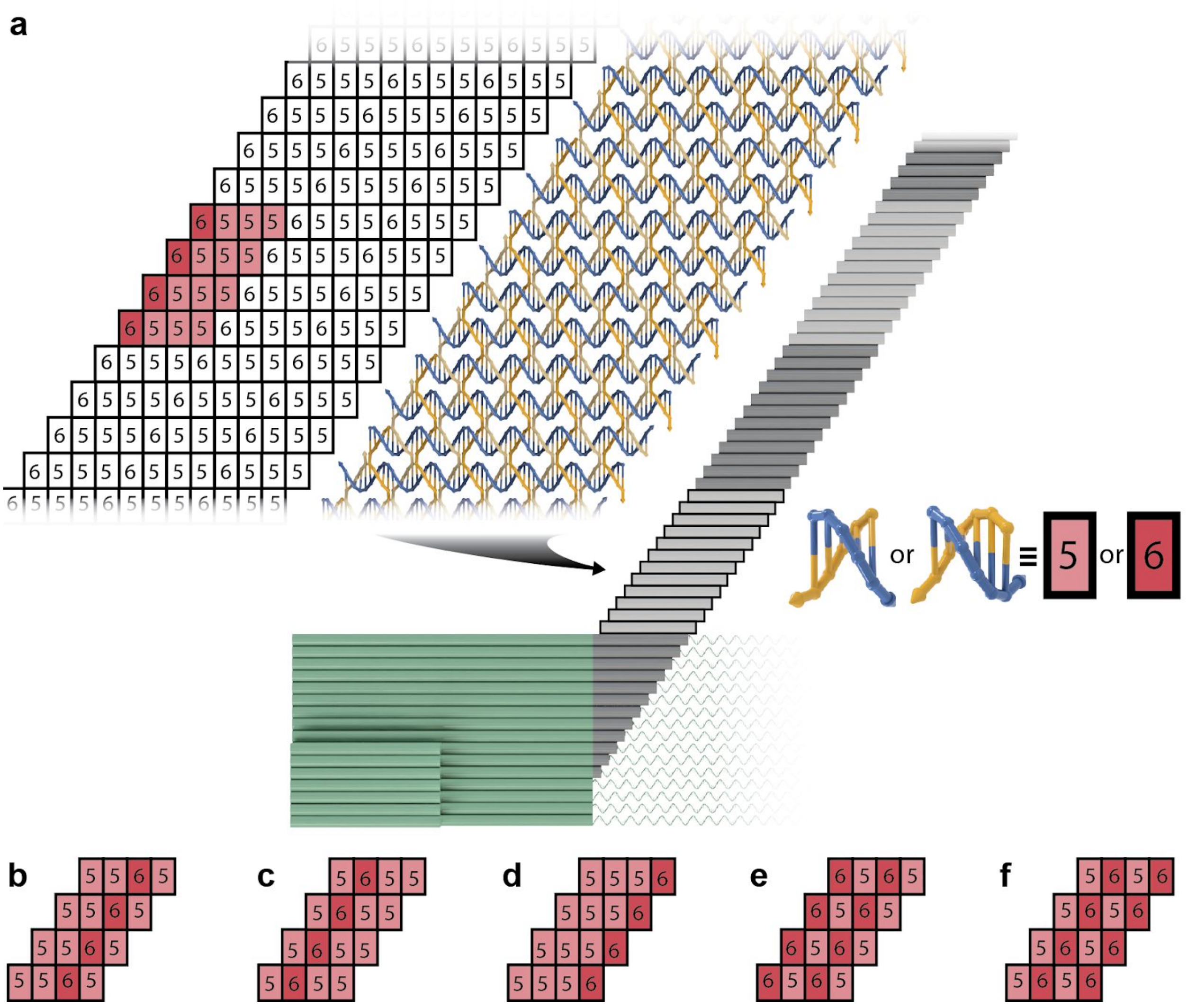


**Supplementary Figure S37**: DNA slats individual binding domain length and pattern arrangement. **a**, Pattern of rectangles with corresponding binding domains of length 5 or 6 bp are displayed next to the 3D DNA ball and stick model. The pattern of four sequential binding domains constituting of either 21 or 22 bp per two helical turns is highlighted in red. The 6555 pattern is used in sequence variants 6.1, 6.2, 6.3, and 8.2. These slat variants all have 10.5 bp/turn. **b**, Binding domain pattern in sequence variant 8.1. v8.1 has 10.5 bp/turn. **c**, Binding domain pattern in sequence variant 8.4. v8.4 has 10.5 bp/turn. **d**, Binding domain pattern in sequence variant 8.5. v8.5 has 10.5 bp/turn. **e**, Binding domain pattern in sequence variant 8.6. v8.6 has 11 bp/turn. **f**, Binding domain pattern in sequence variant 8.3. v8.3 has 11 bp/turn.

#### S8.2—Results from shifting arrangement of binding sites

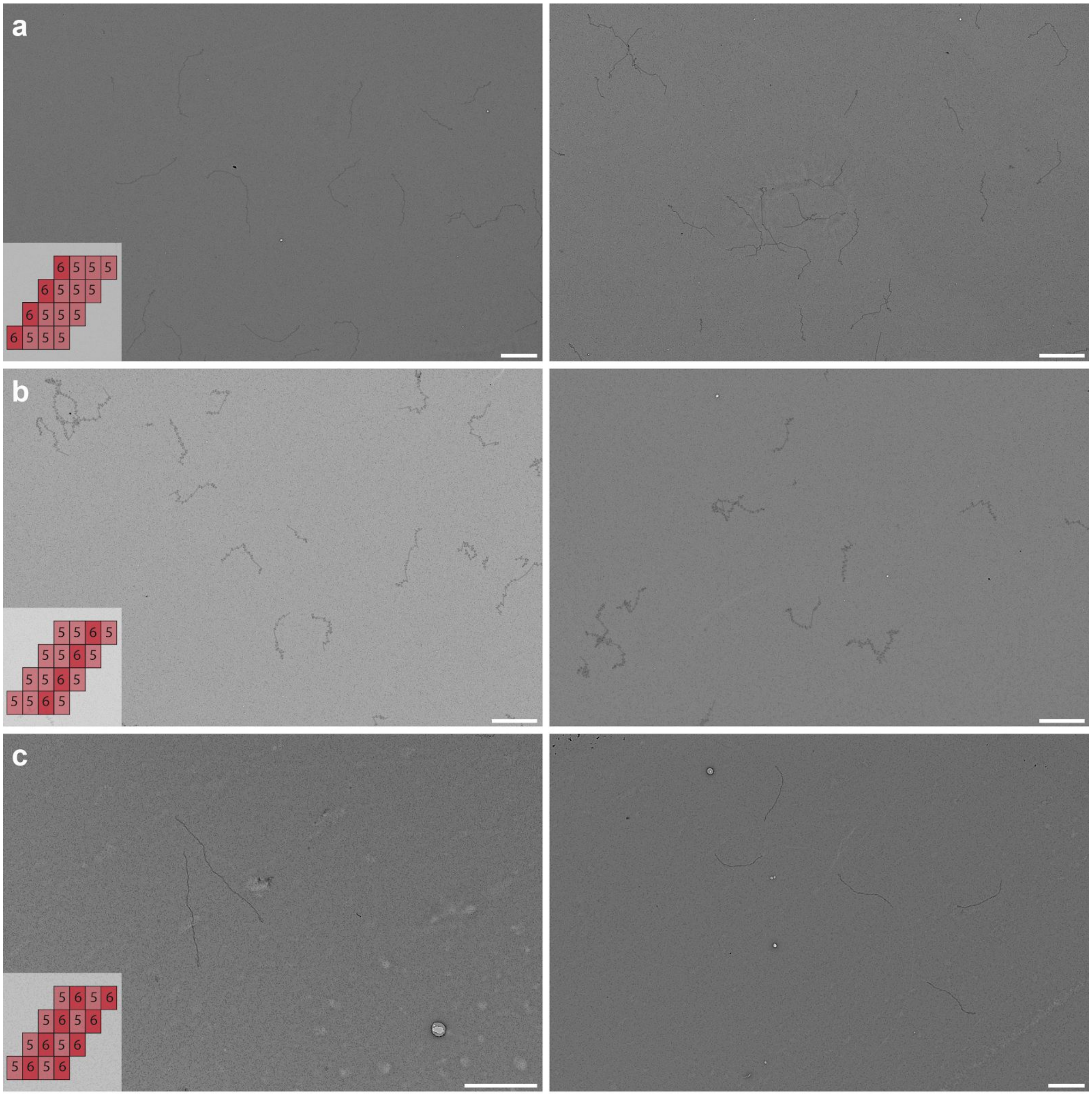


**Supplementary Figure S38**: Negative-stain TEM micrographs showing DNA slats sequence variant 8.1, 8.2, and 8.3 without sticky ends. **a**, Binding domain pattern in sequence variant sequence variant 8.2. DNA slat ribbons grown with v8.2 display very little coiling. We hypothesize that variant 8.2 still has some inherent coiling in solution, which results in kinked ribbons once deposited onto a carbon grid for TEM imaging. However, the kinks are of a much lesser extent in this variant when compared to all of the other tested designs. **b**, Binding domain pattern in sequence variant sequence variant 8.1. This variant displayed loosely coiled ribbons on the TEM micrographs. **c**, Binding domain pattern in sequence variant sequence variant 8.3. This variant displayed tightly coiled ribbons on the TEM micrographs. Both variants shown in panels *a* and *b* were used to assemble tubes. Scale bars are 1 μm.

#### S8.3—Results from adding short sticky ends to close the coiled sheets into tubes

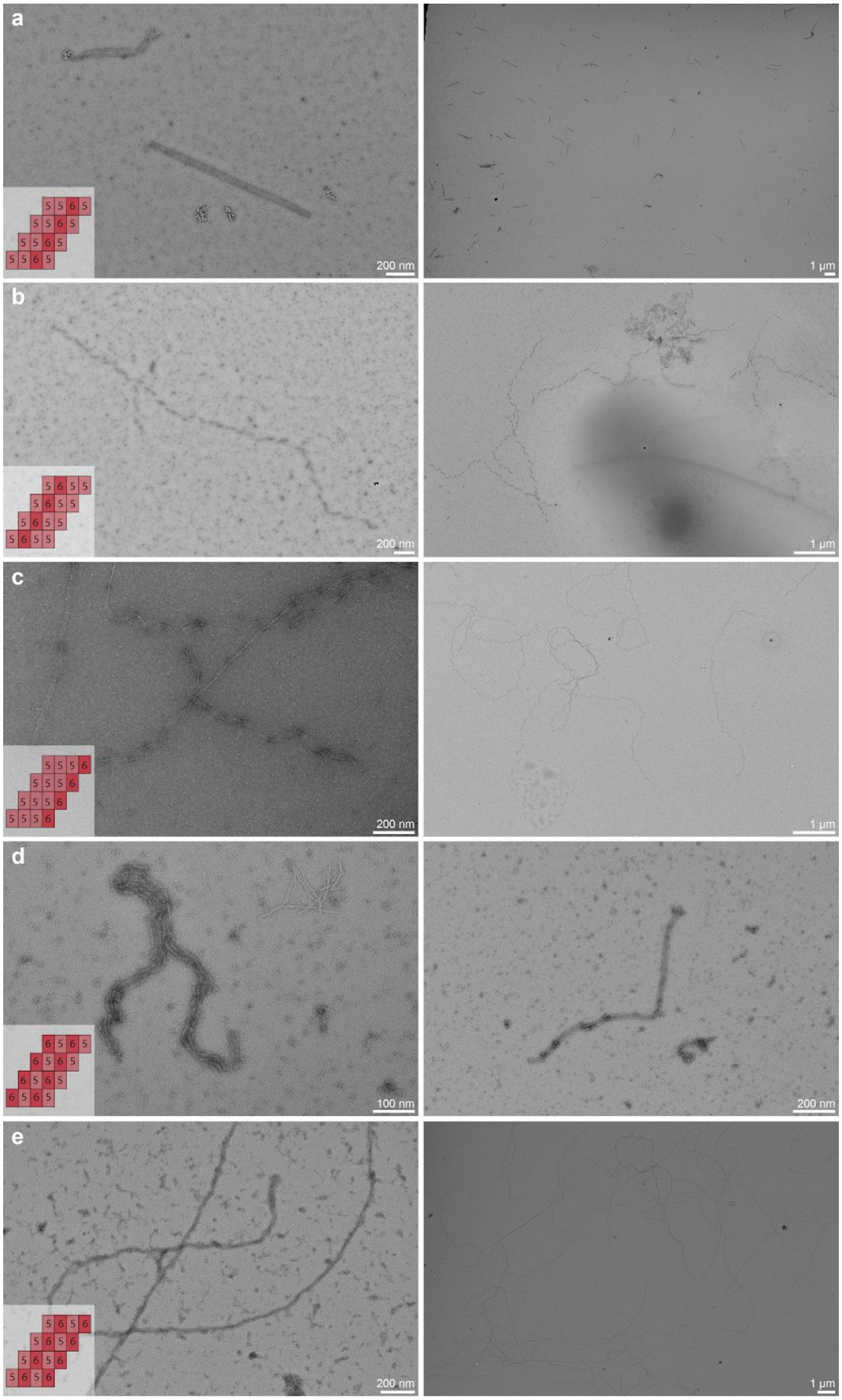

**Supplementary Figure S39**: Negative-stain TEM micrographs showing DNA slats sequence variant 8.1, 8.3, 8.4, 8.5, and 8.6 with 2 nt sticky ends along the edge of the ribbon. **a**, Sequence variant 8.1 with 2 nt sticky ends. This variant forms thick tubes of various diameters in a single reaction. We hypothesize that the loosely coiled nature of v8.1 combined with 2 nt sticky ends, results in the formation of tubes of various diameters. The sticky ends aid in forming a specific diameter at the start of the reaction, at which point the tube continues to grow with a relatively constant diameter. However, we also observed tubes with slight diameter variations along the axial of the tube. **b**, Sequence variant 8.4. This variant resulted in misformed tube structures. **c**, Sequence variant 8.5. This variant resulted in misformed tube structures. **d**, Sequence variant 8.6. This variant resulted in misformed tube structures. **e**, Sequence variant 8.3. This variant forms thin tubes of constant diameter. We hypothesize that the tightly coiled nature of v8.3 combined with 2 nt sticky ends, results in the formation of tubes with constant diameters.

#### S8.4—Faster rate of assembly for 11.0 bp/turn design for underwound v8.3

We approximate the rate of slat addition for v8.3 based on ribbon length measurements shown in **Supplementary Figure S40**. We estimate the second-order rate constant with

$$\tau = t_{1/2}/ln(2) = 1/(kC_0)$$

Where $\tau$ is the e-folding lifetime, $t_{1/2}$ is the half-life, $C_0$ the initial DNA slats concentration, and $k$ the second-order rate constant. If we assume $k = 10^6 M^{-1}s^{-1}$ and set $C_0 = 0.5 \times 10^{-6} M$ (as per experimental results shown in **Supplementary Figure S40**), $\tau = 1/\ (10^6 M^{-1}s^{-1} \times 0.5 \times 10^{-6} M) = 2\ s$. Thus, with the assumption of 1 slat addition every 2 seconds we get a close approximation of the $10^6 M^{-1}s^{-1}$ second-order rate constant (see **Supplementary Table S4**).

| Assem-bly time [s] | Mean length measured [nm] | SD of lengths [nm] | Mean # of slat additions* | SD # of slat additions* | # slat additions assuming $\tau = 2$ and $k = 10^6 M^{-1} s^{-1}$ |
|---|---|---|---|---|---|
| 300 | 275 | 44 | 183.33 | 29.33 | 150 |
| 900 | 770 | 55 | 513.33 | 36.67 | 450 |
| 1800 | 1460 | 207 | 973.33 | 138 | 900 |

* We are assuming 1.5 nm length increase per single slat addition.

**Supplementary Table S4**: Length measurements form **Supplementary Figure S40** used to approximate the second-order rate constant of v8.3 DNA slat assembly.

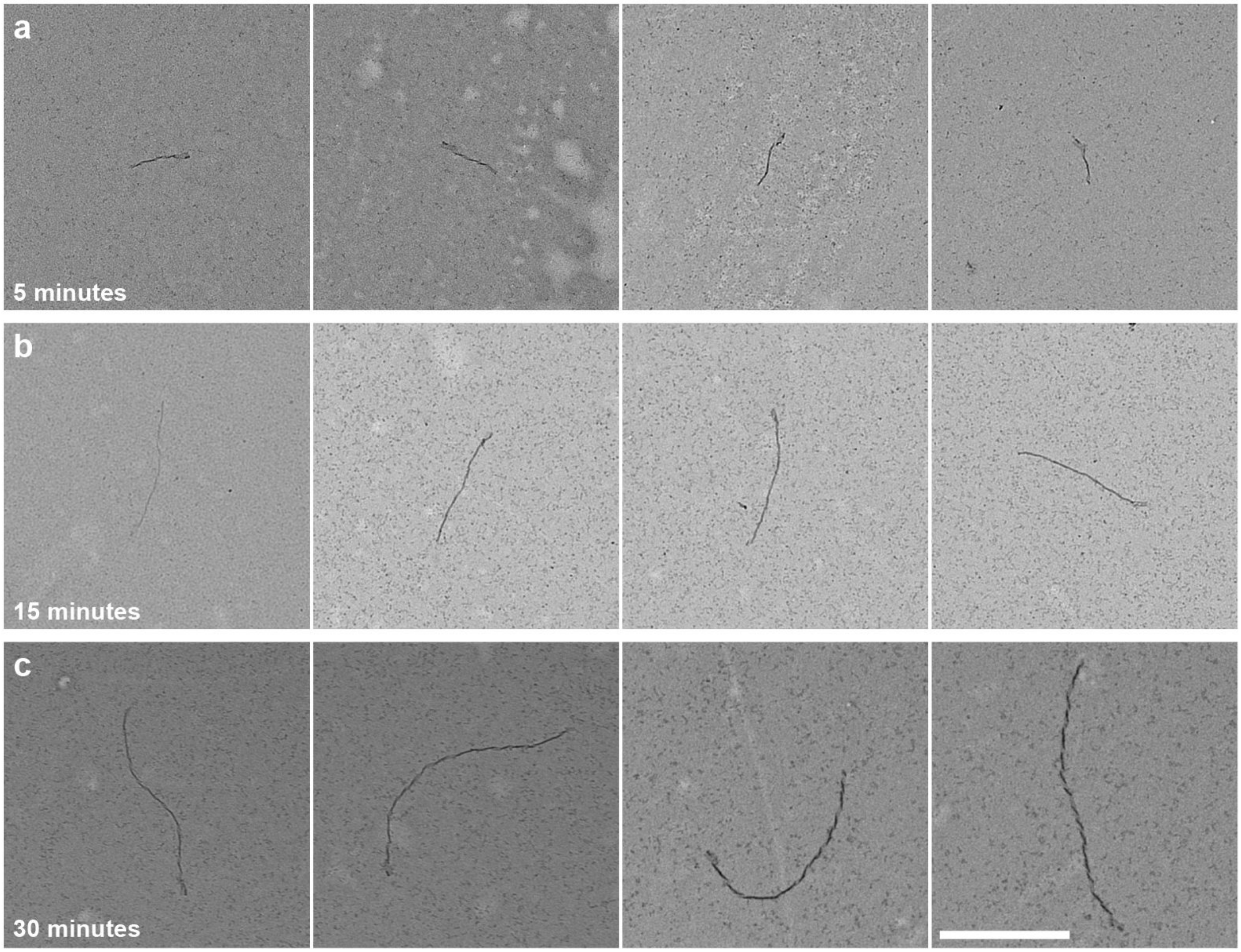


**Supplementary Figure S40**: Negative-stain TEM micrographs for sequence variant 8.3 without sticky ends. DNA slat at 0.5μM were assembled with 0.02 nM seed in 14 mM $Mg^{2+}$ at 52ºC. **a**, 5 minutes of assembly yield 275 nm ± 44 nm in length. **b**, 15 minutes of assembly yield 770 nm ± 55 nm in length. **c**, 30 minutes of assembly yield 1460 nm ± 207 nm in length. We estimated the second-order rate constant to be approx. $10^6$ $M^{-1}s^{-1}$. Mean ± SD for N=4 was computed. Scale bar are 800 nm.

#### S8.5—No observable spurious nucleation for v8.3 under optimal growth conditions (≥52ºC for 1 μM each slat)

In this subsection we determine optimal growth conditions for v8.3. This variant displayed an approximately five-times faster second-order rate constant (see **S8.4**) in comparison to v6.1, v6.2, v6.3, v8.1, and v8.2. To determine the optimal growth conditions we isothermally assembled v8.4 from 46ºC–56ºC, ~16 hours in 16 mM $MgCl_2$ for 0.1 and 1 μM each slat. Optimal growth conditions were especially hard to determine for high (i.e. 1 µM each) slat concentrations, since the v8.3 ribbons grow so long (over the course of 16 hours) that they aggregate in the well of each lane. We therefore added a condition with 0.1 µM each slat and used this to further determine that 52ºC–54ºC seemed to yield optimal assembly for v8.3.

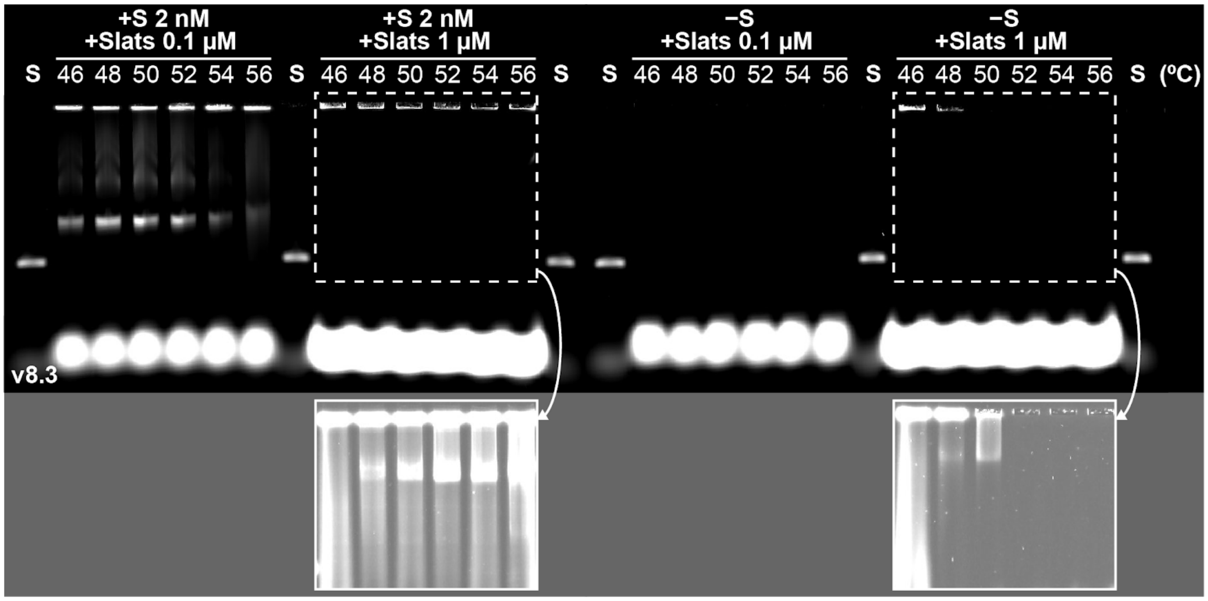


**Supplementary Figure S41**: Gel characterization of seeded (left) and unseeded (right) growth of DNA v8.3 slats. Reactions with either 0.1 or 1.0 µM each slat were incubated isothermally at one of six temperatures in 16 mM $MgCl_2$ for ~16 hours. Notably, unseeded reactions for 0.1 µM each slat yielded no observable assembly at all conditions tested. Unseeded reactions for 1 µM each slat yielded in spurious assembly (as shown in inserts where contrast and brightness were adjusted for better visualization) only for 50ºC and below. (S) is the seed only control.
